# Proteogenomic Reprogramming to a Functional Human Totipotent Stem Cell State via a PARP-DUX4 Regulatory Axis

**DOI:** 10.1101/2024.06.14.598510

**Authors:** Ludovic Zimmerlin, Ariana Angarita, Tea Soon Park, Rebecca Evans-Moses, Justin Thomas, Sirui Yan, Isabel Uribe, Isabella Vegas, Clara Kochendoerfer, Willem Buys, Anthony K L Leung, Elias T. Zambidis

## Abstract

PARP1 (ARTD1) and Tankyrases (TNKS1/TNKS2; PARP5a/5b) are poly-ADP-ribose polymerases (PARPs) with catalytic and non-catalytic functions that regulate both the genome and proteome during zygotic genome activation (ZGA), totipotent, and pluripotent embryonic stages. Here, we show that primed, conventional human pluripotent stem cells (hPSC) cultured continuously under non-specific TNKS1/TNKS2/PARP1-inhibited chemical naïve reversion conditions underwent epigenetic reprogramming to clonal blastomere-like stem cells. TIRN stem cells (TIRN-SC) concurrently expressed hundreds of gene targets of the ZGA-priming pioneer factor DUX4, as well as a panoply of four-cell (4C)-specific (*e.g*., *TPRXL, HOX* clusters), eight-cell (8C)-specific (*e.g*., *DUXA, GSC, GATA6*), primitive endoderm-specific (*e.g*., *GATA4, SOX17*), trophectoderm-specific (*e.g., CDX2, TFAP2C*), and naïve epiblast-specific (*e.g*., *DNMT3L, NANOG, POU5F1(OCT4)*) factors; all in a hybrid, combinatorial single-cell manner. Mapping of proteomic and single-cell expressions of TIRN-SC against human preimplantation embryo references identified them as relatively homogenous 4C-8C stage populations. Injection of TIRN cells into murine 8C-16C-staged embryos resulted in efficient totipotent-like single cell contributions of human cells to both extra-embryonic (trophectoderm, placenta) and embryonic (neural, fetal liver, hematopoietic) lineages in human-murine blastocyst and fetal chimeras. Pairing of proteome with ubiquitinome analyses of TIRN-SC revealed a global shutdown of ADP-ribosylation, and a perturbed TNKS/PARP1 equilibrium which not only impacted the protein levels of hundreds of TNKS/PARP1 substrates via a rewiring of the ubiquitin-proteosome system (UPS), but also de-repressed expression of hundreds of developmental genes associated with PARP1 suppression. ChIP-Seq analysis of core NANOG-SOX2-OCT4 (NSO) pluripotency factors in TIRN-SC identified reprogrammed DUX4-accessible distal and cis-regulatory enhancer regions that were co-bound by PARP1 (NSOP). These NSOP enhancer regions possessed co-binding motifs for hundreds of the same ZGA-associated, embryonic, and extraembryonic lineage-specifying pioneer factors (*e.g*., *HOX, FOX, GATA, SOX, TBX, CDX* families) that were concurrently co-expressed in TIRN-SC; suggesting that PARP1 and DUX4 cooperate with NSO pluripotency core factors to regulate the epigenetic plasticity of a human totipotency program. These findings provide the first demonstration that global, proteome-wide perturbations of post-translational modifications (*i.e*., ADP-ribosylation, ubiquitination) can regulate epigenetic reprogramming during human embryogenesis. Totipotent TIRN-SC will provide a valuable cell culture model for studying the proteogenomic regulation of lineage specification from human blastomere stages and may facilitate the efficient generation of human organs in interspecies chimeras.

## INTRODUCTION

PARP1 and TNKS1/2 are PARP family enzymes that catalyze post-translational poly-ADP-ribose (PAR)ylation of protein substrates and play formative roles in regulating chromatin structure, transcription, and ubiquitination in not only adult cells, but also during embryonic development (Imamura et al., 2004; Zimmerlin and Zambidis, 2020). Chemical inhibition of PARylation impairs the homeostasis and progression of preimplantation embryonic development (Imamura et al., 2004); failure to degrade PAR is embryonic lethal (Koh et al., 2004). TNKS1/2 activity controls developmental progression of zygotic genome activation (ZGA) (Gambini et al., 2020), and PARP1 regulates histone ubiquitination, DNA methylation, RNA processing, DNA repair during ZGA (Hamazaki et al., 2015; Osada et al., 2013; Tsai et al., 2023), cellular reprogramming (Chiou et al., 2013; Doege et al., 2012; Jiang et al., 2015), pluripotency (Roper et al., 2014), trophoblast specification (Hemberger et al., 2003) and chromatin plasticity (Ciccarone et al., 2017; Ko and Ren, 2012) during preimplantation embryogenesis. Although loss of a single PARP protein minimally impairs development (suggesting redundancy of PARP proteins), compound TNKS1/TNKS2 and PARP1/PARP2 knockouts, or dominant-negative PARP1 are embryonic lethal (Chiang et al., 2008; Hsiao et al., 2006; Menissier de Murcia et al., 2003; Shao et al., 2023).

PARP1 and TNKS1/2 both possess catalytic and non-catalytic activities via multifunctional protein domains. For example, PARP1 not only ADP-ribosylates SOX2 during cellular reprogramming (Weber et al., 2013) and pluripotency (Lai et al., 2012), but also binds chromatin at its non-catalytic DNA-binding domain to facilitate the pioneer activities of SOX2 (Liu and Kraus, 2017). TNKS1/2 proteins similarly mediate non-PAR-catalytic activities such as DNA repair regulation and pexophagy/autophagy (Bhardwaj et al., 2017; Li et al., 2017; Nagy et al., 2016). Furthermore, the catalytic activities of TNKS and PARP1 deeply impact homeostasis of the proteome (proteostasis) during embryogenesis, via extensive networks of post-translational PAR-dependent ubiquitination (PARdU) (Callow et al., 2011; Cho-Park and Steller, 2013; Rona et al., 2018; Vittal et al., 2015; Vivelo et al., 2019). PARdU impacts the longevity and stability of histones and transcription factors via the ubiquitin-proteasome system (UPS), thus indirectly regulating developmental progression (Choi and Baek, 2018; Strikoudis et al., 2014; Yan et al., 2020), ZGA (Higuchi et al., 2018), and pluripotency (Buckley et al., 2012; Schroter and Adjaye, 2014; Suresh et al., 2016; Vilchez et al., 2012; Wu and Zhang, 2021). For example, protein stability of the WNT signaling ligand Axin1 is regulated by TNKS1/2-mediated PARylation of E3 ligase RNF146, which ubiquitinates Axin1 for subsequent UPS degradation (Callow et al., 2011). Direct control of the stability and assembly of the proteasome 26S subunit itself by TNKS ADP-ribosylation has also been suggested (Cho-Park and Steller, 2013). Beyond the UPS, post-translational, UPS-independent (Mukhopadhyay and Riezman, 2007; Trulsson et al., 2022) ubiquitination structurally regulates chromatin/histone architecture (Vaughan et al., 2021), and transcription factor DNA binding (Mark and Rape, 2021) (including of the core pluripotency factors NANOG (Kim et al., 2014; Pei, 2017), OCT4(Baek et al., 2020; Li et al., 2018; Rhie et al., 2021) and SOX2(Cui et al., 2018; Fang et al., 2014a; Wang et al., 2016)). Indeed, the ubiquitin machinery itself (Rona et al., 2018; Vittal et al., 2015) (*e.g*., ubiquitin ligases and deubiquitinases (DUBs)) which write and erase the proteome’s ‘ubiquitin code’ (Dikic and Schulman, 2023) during embryonic development (Cruz Walma et al., 2022; Xie et al., 2019), are regulated by PARP1-mediated PARylation (Callow et al., 2011; Vivelo et al., 2019).

Several studies have employed chemical PARP inhibition to expand pluripotency and improve the chimeric contribution of mouse embryonic stem cells (mESC) into embryos (Gao et al., 2019; Liu et al., 2021; Yang et al., 2017a; Yang et al., 2017b), or to improve functional pluripotency (Park et al., 2020; Park et al., 2018; Thomas et al., 2021; Zimmerlin et al., 2016; Zimmerlin et al., 2017). Zimmerlin et al (Zimmerlin et al., 2016) first demonstrated that primed, conventional human pluripotent stem cells (hPSC) were chemically reverted to a naive epiblast-like pluripotent state with improved differentiation potential via a two-step chemical reprogramming system (‘LIF-5i -> LIF-3i’; **Fig. S1a**), that comprised of sequential culture with LIF and five small molecules (*i.e*., Hedgehog signalling activation, cAMP agonism, GSK3ϕ3 inhibition, MEK inhibition, and nonspecific TNKS1/TNKS2/PARP1 inhibition (XAV939). Sequential naïve reversion with continuous exposure to XAV939 reprogrammed a large repertoire of >25 conventional lineage-primed hPSC to tankyrase/PARP inhibitor-regulated naïve (TIRN) stem cells (Park et al., 2020; Park et al., 2018; Thomas et al., 2021; Zimmerlin et al., 2016; Zimmerlin et al., 2017). TIRN-SC possessed high clonal proliferation, phosphorylated-STAT3 signalling, MEK-ERK/bFGF signalling independence, modulation of ϕ3-catenin expression, and erasure of lineage-primed gene expression (Park et al., 2020; Zimmerlin et al., 2016). TIRN-SC also maintained normal karyotypes and imprinted CpG patterns, and were devoid of irreversible demethylation defects (Thomas et al., 2021; Zimmerlin et al., 2016) that were reported in naïve reversion systems (Pastor et al., 2016; Theunissen et al., 2016), and suggested to be secondary to prolonged MEK inhibition (Choi et al., 2017). Reversion to TIRN-SC eliminated interline variability and significantly enhanced the multi-lineage differentiation performance of isogenic primed hPSC differentiation; without requirement for transitioning back to primed culture conditions to restore differentiation potency (Lee et al., 2017; Sahakyan et al., 2017; Warrier et al., 2017). Although the mechanism of improved functional pluripotency of TIRN-SC was unclear, the pleiotropic effects of XAV939 likely played a role. Although XAV939 is a TNKS inhibitor of the canonical WNT pathway (*e.g*., via TNKS-mediated PARdU of Axin1), it is a promiscuous small molecule that inhibits *in vitro* PARylation activities of not only TNKS1 and TNKS2 (IC_50_ ∼5.2-94.6 nM), *but also* PARP1 and PARP2 (IC_50_ ∼26.9-169 nM) (Huang et al., 2009; Thorsell et al., 2017). Thus, XAV939 is expected to inhibit *both* PARP1 and TNKS1/2 at the supra-micromolar concentrations used by Zimmerlin *et al*; with potential impact on broad biological processes dependent on ADP-ribosylation.

Here, we show that XAV939-treated TIRN-SC were globally inhibited in both TNKS and PARP1 PARylating activities. TNKS1/2 protein levels were *increased* while PARP1 protein levels *decreased* in TIRN-SC; potentially defining a novel functional PARP protein equilibrium. PAR-deficient TIRN-SC underwent a global ubiquitinome reprogramming that was driven by hundreds of PARP1 and TNKS1/2 targets; including core factors NANOG and SOX2 and epigenetic modifiers. Unexpectedly, XAV939-mediated TIRN reprogramming activated simultaneous, single-cell co-expressions of ZGA-priming (e.g., *TPRXL)* and DUX4 gene targets, as well as a panoply of PARP1-regulated lineage-specifying transcriptional factors. TIRN-SC possessed totipotent-like functionality including efficient capacity to contribute differentiated human cells into both embryonic and extraembryonic tissues of human-murine chimeras, and efficient generation of trophoblast stem cells (TSC) with *in vivo* placental potency. TIRN-SC underwent a genome-wide epigenetic reprogramming driven by reduction and redistribution of PARP1 chromatin co-binding to core pluripotency factors NANOG/SOX2/OCT4 (NSO) at gained promoter, intragenic, and distal enhancer sites of 8C- and morula-associated genes (*e.g., EBF2*, *HOXB2*, *GATA6*). A cohort of gained NSO and PARP1 (NSOP) co-binding enhancer sites were identified near transcriptionally active, as well as distal regions of DUX4-accessible chromatin. Remarkably, these gained NSOP-DUX4 co-binding sites were further enriched in co-binding motifs for *hundreds* of the same embryonic and extraembryonic lineage-determining pioneer factors that were actively and concurrently already co-expressed in TIRN-SC, suggesting the discovery of candidate totipotency-regulating chromatin regions.

## RESULTS

### TIRN-SC expressed transcriptional and proteomic signatures of human cleavage-stage embryo cells

The LIF-5i->LIF-3i method mediates efficient reversion of conventional primed hPSC cultures into TIRN-SC with high functional pluripotency and reduction of interline variability of multi-lineage differentiation (Park et al., 2018; Zimmerlin et al., 2016) (**Fig. S1a,b**). An array of genetically independent TIRN-SC lines reproducibly and stably acquired activated phosphorylated STAT3 signaling, naïve epiblast-specific transcripts and markers, decreased ERK1/2 phosphorylation, and upregulated TNKS1/2 proteins for at least 30 passages in single-cell TIRN conditions (**Figs. 1a; S1c-e**). To investigate the impact of genetically-driven variabilities in the TIRN system (which is a key determinant in naïve pluripotent cultures (Ortmann et al., 2020; Skelly et al., 2020)), we performed comparative RNA-seq bioinformatics analysis of eight genetically-independent TIRN-reverted hESC and hiPSC lines, and their isogenic, primed counterparts. These studies revealed significant global differential expressions of developmentally important transcriptional regulators (**Fig. S1f, Table S1**) in TIRN-SC relative to their isogenic primed hPSC counterparts. TIRN-SC lines *concurrently expressed* high transcript levels of *not only* naïve epiblast-specific genes (*e.g*., *DNMT3L, NANOG, KLF17*), but also unexpectedly, a broad panoply of 2C-4C-specific maternal (*e.g*., *TPRXL, HOX A/B/C* clusters, *KHDC3L*), 8C-specific (*e.g*., *DUXA, EBF2, GSC, GATA6*), meso-endoderm lineage-specific (*e.g*., *GATA4, SOX17, EOMES, FOXA2, HAND1, MIXL1, TBXT*), and trophectoderm-specific (*e.g., CDX2, TFAP2C, GATA3*) genes; all in a combinatorial manner (**Fig. 1b**).

**Figure 1.**
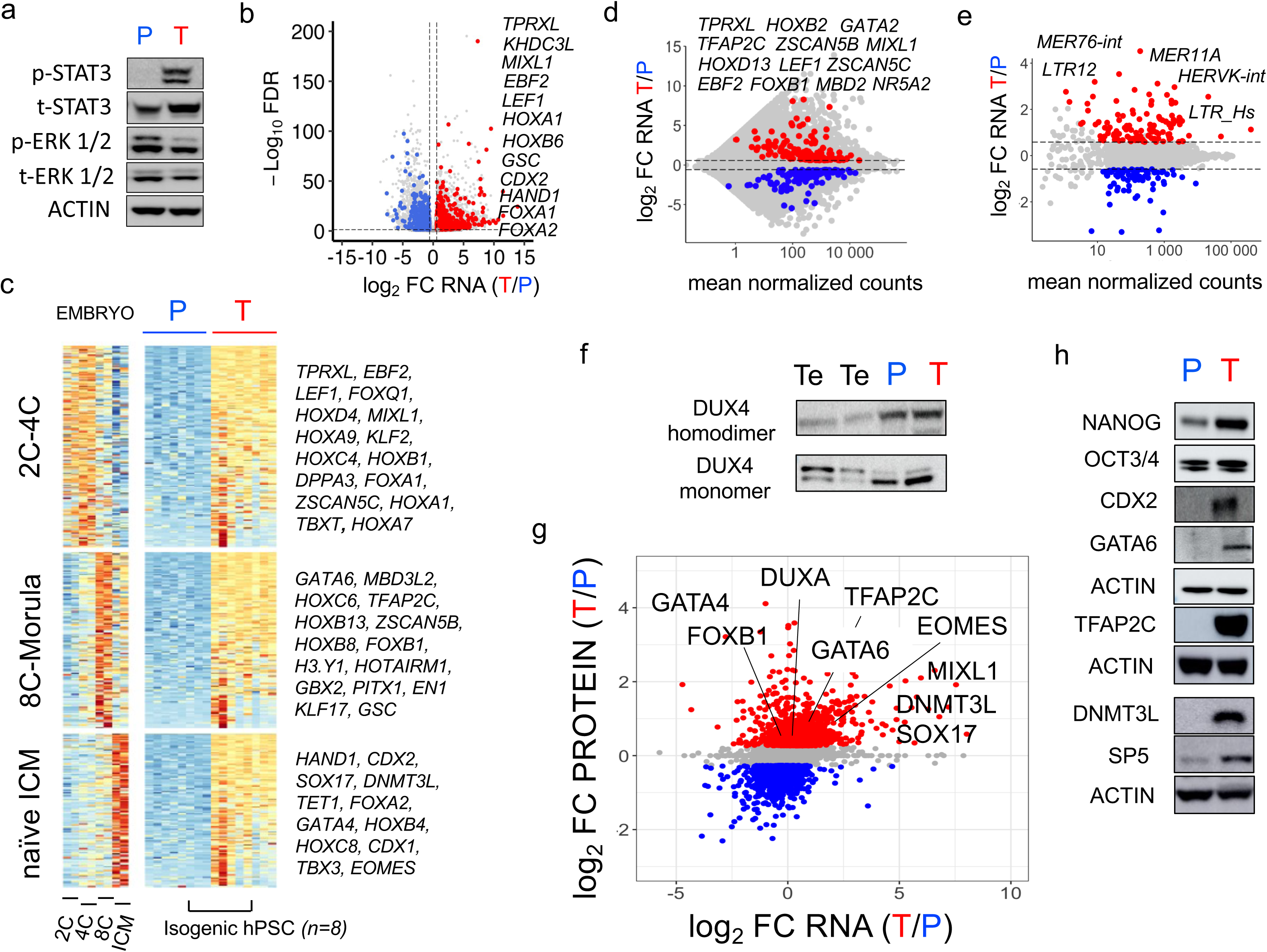
Activation of human cleavage stage embryonic expression programs in TIRN-SC. (**a**) Western blot analysis of phosphorylated (p) and total (t) STAT3 and ERK1/2 in isogenic primed (P) vs TIRN (T) stem cells. (**b**) Volcano plot showing log2 fold change (FC>1.5; p<0.05) RNA expression highlighting only differentially expressed transcriptional regulators (Lambert et al., 2018) (**Table S1**) between TIRN (red) vs. primed hPSC (n=8 independent cell lines); all other genes are greyed. (**c)** Heatmaps of RNA seq data (FPKM) comparing the most significantly expressed (FC>1.5, adjusted p-value <0.05) transcriptional regulators (Lambert et al., 2018) (n=392 genes) in 8 isogenic primed (P) and TIRN (T) hPSC lines expressed during indicated human preimplantation stages. Correlative expression of same genes from human embryo Ribo-Seq translatome data at indicated stages (Zou et al., 2022) (FPKM) is shown on the left heatmap as control. (**d**) MA plot showing bulk RNA expression in paired T (red) and P (blue) stem cells (n=8). Colored dots show transcription factors with a promoter (±3kb) bound by the pioneer factor DUX4 (Hendrickson et al., 2017) with the most significant expression (fold change >1.5, p < 0.05). (**e**) MA plot showing expression of transposable elements (TE) in T (red) and P (blue) stem cells (colored dots represent TE with the most significant expression (fold change >1.5, p < 0.05). (**f**) Western blot analysis of DUX4 protein expression in control testis (Te), primed (P) and TIRN (T) lysates using antibodies that recognize homodimer or monomer isoforms. (**g**) Crossplot of log2 fold change (FC) of RNA-seq (x-axis; RNA) vs whole proteome (y-axis; PROTEIN) data showing significantly expressed (FC>1.2, q<0.05) proteins in TIRN (T; red, 961 proteins) vs primed (P; blue, 992 proteins) hPSC (*n*=3-6 replicates for each condition). (**h**) Western blot confirmation of selected proteome findings of core pluripotency (NANOG, OCT3/4), 8C-specific (GATA6) trophectoderm-specific (CDX2, TFAP2C) and naïve epiblast-specific (DNMT3L, SP5) proteins overexpressed in TIRN-SC.

The hybrid expression of multiple lineage-specific pioneer factors expressed simultaneously along with pluripotency factors has been reported to be a characteristic of epigenetically plastic totipotent and pre-lineage morula stages in human, mouse, and zebrafish embryos (Farrell et al., 2018; Lee et al., 2013; Petropoulos et al., 2016; Redo-Riveiro et al., 2024; Thompson et al., 2022; Zou et al., 2022). To delineate the developmental stage of TIRN-SC overexpressing diverse pioneer factors that not only specify committed lineages during blastocyst and gastrulation stages, but are also expressed in a combinatorial fashion during totipotent 4C-8C pre-blastocyst embryonic stages (*i.e*., *HOX, GATA, T-BOX, FOX* families; **Fig. S1h, Table S1**), we generated gene clusters that specify pre-lineage embryonic stages from 2C to naïve inner cell mass (ICM) by k-mean clustering of a human preimplantation embryo Ribo-seq translatome data set (Zou et al., 2022). This approach revealed that TIRN-SC were highly enriched in hundreds of translatable multi-lineage-specifying pioneer factors that are co-expressed at pre-ZGA (2C-4C), ZGA/totipotent (4C-8C), and morula human embryonic stages (**Fig. 1c, Table S1**). Many of the 4C-8C-specific genes overexpressed in TIRN-SC (*e.g*., *TPRXL, ZSCAN5B, HOXB2, EBF2*) were targets reported to be bound at their promoters by the ZGA-priming pioneer factor DUX4 (Hendrickson et al., 2017; Vuoristo et al., 2022; Yoshihara et al., 2022). Indeed, we identified >200 cleavage stage transcriptional factors and >70 transposable elements (*i.e*., TE’s; *e.g*., *MER11A, MER76-int, HERVH-int*) differentially overexpressed in TIRN cells reported to be cleavage-stage DUX4 gene targets (**Fig. 1d,e Table S1**) (Hendrickson et al., 2017). However, both primed hPSC and TIRN-SC expressed comparably low endogenous levels of homodimeric and monodimeric isoforms of DUX4 protein (**Fig. 1f**); suggesting an alternate mechanism for the observed DUX4 gene target activation in TIRN cells.

The human embryo Ribo-seq reference data (Zou et al., 2022) we employed predicted that the ZGA-to-morula-specific transcripts expressed in TIRN RNA-seq data are potentially translated. To validate protein expression of TIRN-SC RNA-seq results, we performed whole proteome analysis of TIRN-SC vs isogenic primed hPSC (**Figs. 1g, S1g, Table S2**). These studies confirmed *simultaneous* protein expressions of developmentally disparate embryonic lineages in TIRN-SC (*e.g*., naïve epiblast proteins NANOG, OCT4 (POU5F1), DNMT3L, SP5, along with 8C-specific proteins DUXA, GATA6, primitive endoderm lineage-specific proteins GATA4, SOX17, and trophectoderm lineage-specific proteins CDX2, TFAP2C. Selected proteome results were confirmed by Western blotting of TIRN-SC lysates (**Fig. 1h**).

### Transient, inducible DUX4 (iDUX4) transgenesis augmented and completed the baseline blastomere-stage expression pattern of TIRN-SC to a 4C-8C totipotent cell signature

Ectopic, transient expression of full-length DUX4 was reported to be toxic in both primed and naïve hESC, albeit partially and transiently activated subsets of 8C/morula-specific target genes (Hendrickson et al., 2017; Vuoristo et al., 2022; Yoshihara et al., 2022). Although TIRN-SC overexpressed hundreds of 4C-8C-specific DUX4 target genes (**Fig. 1d,e, Tables S1, S3**), including *TPRXL*, *HOXB2*, *EBF2*, and *GATA6*, both primed and TIRN-SC expressed comparably low levels of endogenous germline-associated mRNA splice variants of *DUX4* (Das and Chadwick, 2016; Snider et al., 2010) (**Fig. S2a**). To define a role for DUX4 regulation of cleavage-stage TIRN-SC, we generated codon-altered doxycycline-inducible full-length DUX4 (iDUX4) lines (Jagannathan et al., 2016) from isogenic primed hPSC and TIRN-SC (**Fig. S2b,c**), and compared the transcriptomes of primed, primed-iDUX4, TIRN, and TIRN-iDUX4 cells (**Fig. 2a,b)**.

**Figure 2.**
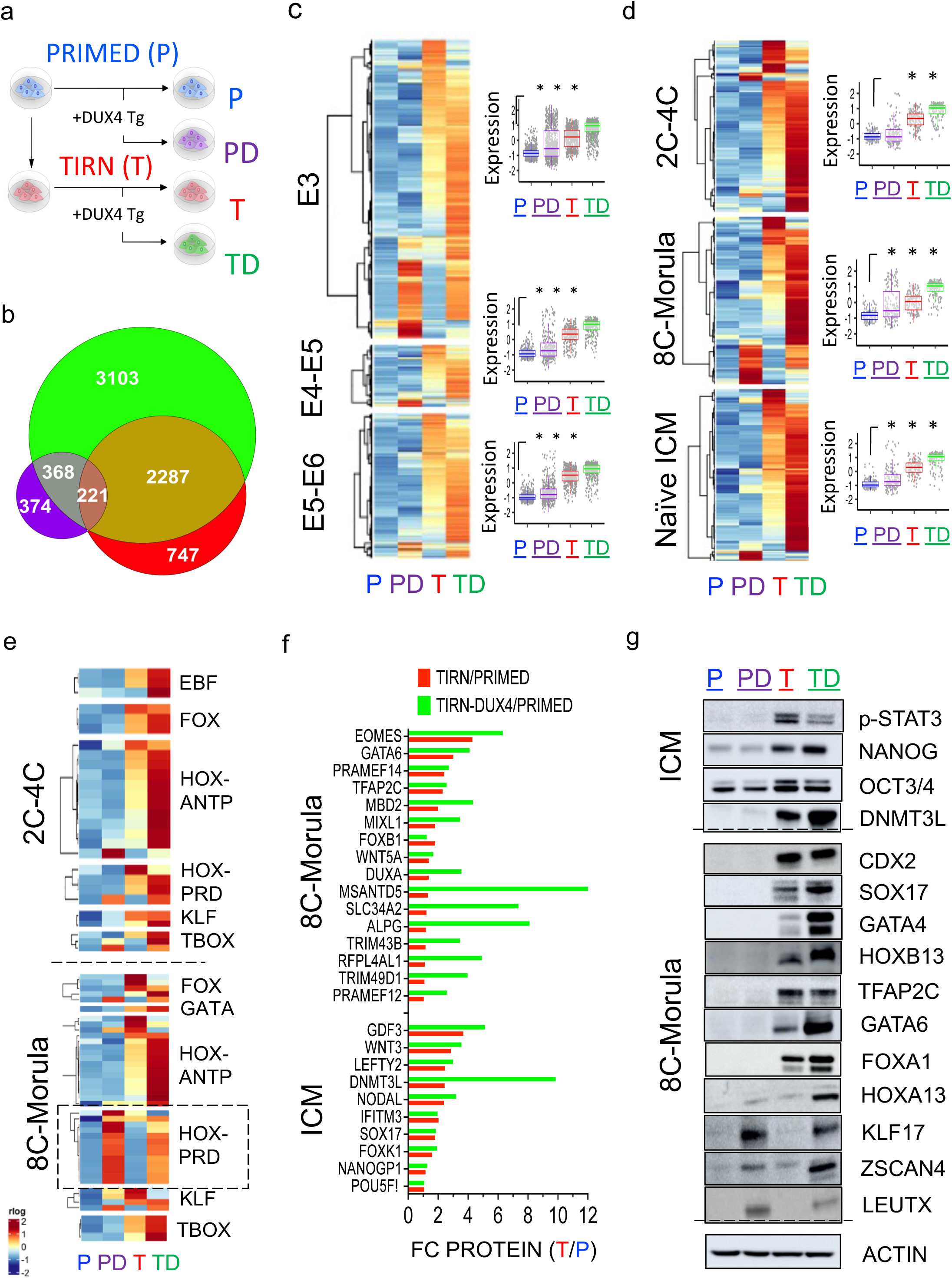
Reinforcement of a totipotent-like expression signature in TIRN-SC following iDUX4 expression. (**a**) Experimental design of 4 conditions to assess the contribution of transgenic inducible DUX4 expression (iDUX4): primed (P, blue), primed + iDUX4 (PD, purple), TIRN (T, red) and TIRN+ iDUX4 (TD, green) cells (**b**) Euler diagram of number of differentially upregulated genes from primed hPSC (n=3; fold change (FC) >1.5, p<0.05) in PD (purple; n=3), T (red; n=6) and TD (green; n=6) samples. Expression of iDUX4 in primed cells (PD) had a more limited impact than in TIRN cells (TD); TIRN conditions (T) had a strong overlap with TD-expressing cells. (**c**) Heatmap of RNA-seq data (regularized log counts) for mean expression in all 4 conditions split by gene expression modules that define the E3 (8C), E4-E5/early pre-lineage (cleavage-morula) stage and E5/late-E7 epiblast stages in human embryos (Petropoulos et al., 2016). Scaled mean-subtracted normalized count data is summarized as boxplots on right. *; ANOVA p<2e^-16^. (**d**) Heatmap of RNA-seq data (FPKM) for mean expression of transcriptional regulators in all 4 conditions split by gene expression modules that define the 2C-4C, 8C and naive ICM stages in human embryos (Zou et al., 2022). Scaled normalized count data is quantitated as boxplots, *; ANOVA p<2e^-16^. (**e**) Heatmap of RNA-Seq data (regularized log counts) for mean expression of pioneer factors in all 4 conditions split by transcription factor families for 2C-4C and 8C-morula stages. Although iDUX4 uniquely activated a small of subset of HOX-PRD factors in PD and TD conditions (dashed box), TIRN-SC already expressed a broad repertoire of ZGA-Morula-specific pioneer factor genes; these expressions were further enhanced by iDUX4. (**f**) Differential expression analysis of proteomics data for 8C and naïve ICM proteins in TIRN and TIRN + iDUX4 cells relative to primed control samples. TIRN-SC expressed proteins identified in the translatome of early human embryos (Zou et al., 2022); that were further enhanced by iDUX4. (**g**) Western blot protein confirmation of selected 8C/morula and naïve ICM factors identified in whole proteomics experiments in P/PD/T/TD lysates.

As reported (Hendrickson et al., 2017; Vuoristo et al., 2022; Yoshihara et al., 2022), high level transgenic DUX4 expression was toxic in primed hESC +iDUX4 and TIRN +iDUX4 at >36 hours following induction, but at 12-24 hours, a limited expression of 8C/morula-specific DUX4 gene targets were upregulated (many which were already overexpressed in unmodified TIRN-SC) (**Fig. S2d-f**). To delineate the developmental identity of TIRN and TIRN + iDUX4 cells, we further employed mRNA expression (Petropoulos et al., 2016) and translatome (Zou et al., 2022) human embryo datasets to define gene sets specifying 2C-4C (pre-ZGA), 8C (E3), pre-lineage morula (E4-E5), and naïve ICM (E5-E6) stages in primed +/- iDUX4 and TIRN +/- iDUX4 cells (**Fig. 2c-e**). Although TIRN-SC robustly expressed cohorts of hundreds of transcriptional regulators (**Fig. S2f**) that specified 2C-4C, 8C, and morula-stage human embryos (Zou et al., 2022) (*e.g*., including *HOX-paired-like domain (HOX-PRD), HOX-antennapedia (HOX-ANTP), FOX, GATA, TBOX* pioneer family genes; **Fig. 2e**, and the 4C-specific ZGA regulator *TPRXL* (Zou et al., 2022)), these expressions were amplified in TIRN-SC upon iDUX4 (**Figs. 2c-e, S2d-f, Table S3**). However, iDUX4 activated expression of an *additional but limited* set of 8C-specific totipotency-associated gene targets (*i.e*., *LEUTX, ZSCAN4, TPRX1, ARGFX*) in TIRN-SC. Overall, transient iDUX4 significantly reinforced a ZGA-to-morula-specific program of transcripts (**Fig. 2c-e**) and proteins (**Fig. 2f,g**) that were already expressed in TIRN-SC.

### TIRN-SC were homogenous blastomere-like populations co-expressing ZGA-specific, 8C-specific, and naïve ICM pre-lineage-specific genes in single cells, in a hybrid, simultaneous manner

The bulk RNA-seq and proteomics analyses of TIRN-SC (+/- iDUX4) revealed acquisition of pre-lineage embryonic transcriptional programs. However, rare subsets of cells expressing 8C-specific genes were reported in heterogenous naïve epiblast-like hESC populations (Moya-Jodar et al., 2023). To determine if TIRN +/- iDUX4 cells were mixed populations or clonal blastomere-like cells, single cell (sc)RNA-seq was performed to investigate transcriptional heterogeneity at a single cell level. Overall, single, unmodified TIRN-SC shared high transcriptional identity with single TIRN+iDUX4 cells (**Fig. 3a**), and single primed hPSC and TIRN +/- iDUX4 cells expressed comparable and uniformly homogenous levels of core pluripotency factors (*e.g*., *NANOG, POU5F1 (OCT4), SOX2, KLF4, CMYC;* **Fig. S3**). However, single TIRN and TIRN+iDUX4 stem cell populations homogenously co-expressed not only higher levels of naïve ICM-specific genes (*e.g*., *DNMT3L, SP5, ERVH48-1, GDF3,* and *IFITM1;* **Fig. S3**) (Park et al., 2020; Zimmerlin et al., 2016), but also hundreds of diverse and antagonistic lineage-specifying genes (*e.g*., *HOX-PRD, T-BOX, FOX, GATA* families) that are co-expressed simultaneously only during 4C-to-morula human embryonic stages (Petropoulos et al., 2016; Zou et al., 2022). Transcript levels of 4C-specific (*i.e*., *TPRXL, HOXA1, EBF2*), 8C-specific (*i.e., GATA6, FOXB1*), trophectoderm lineage-specific (*i.e*., *CDX2*, *TFAP2C*), and endoderm lineage-specific (*i.e., GATA4*, *SOX17*) pioneer factors were co-expressed in the same single cell populations; with expressions that were further augmented following iDUX4 activation (**Figs. 3c-d; S3**).

**Figure 3.**
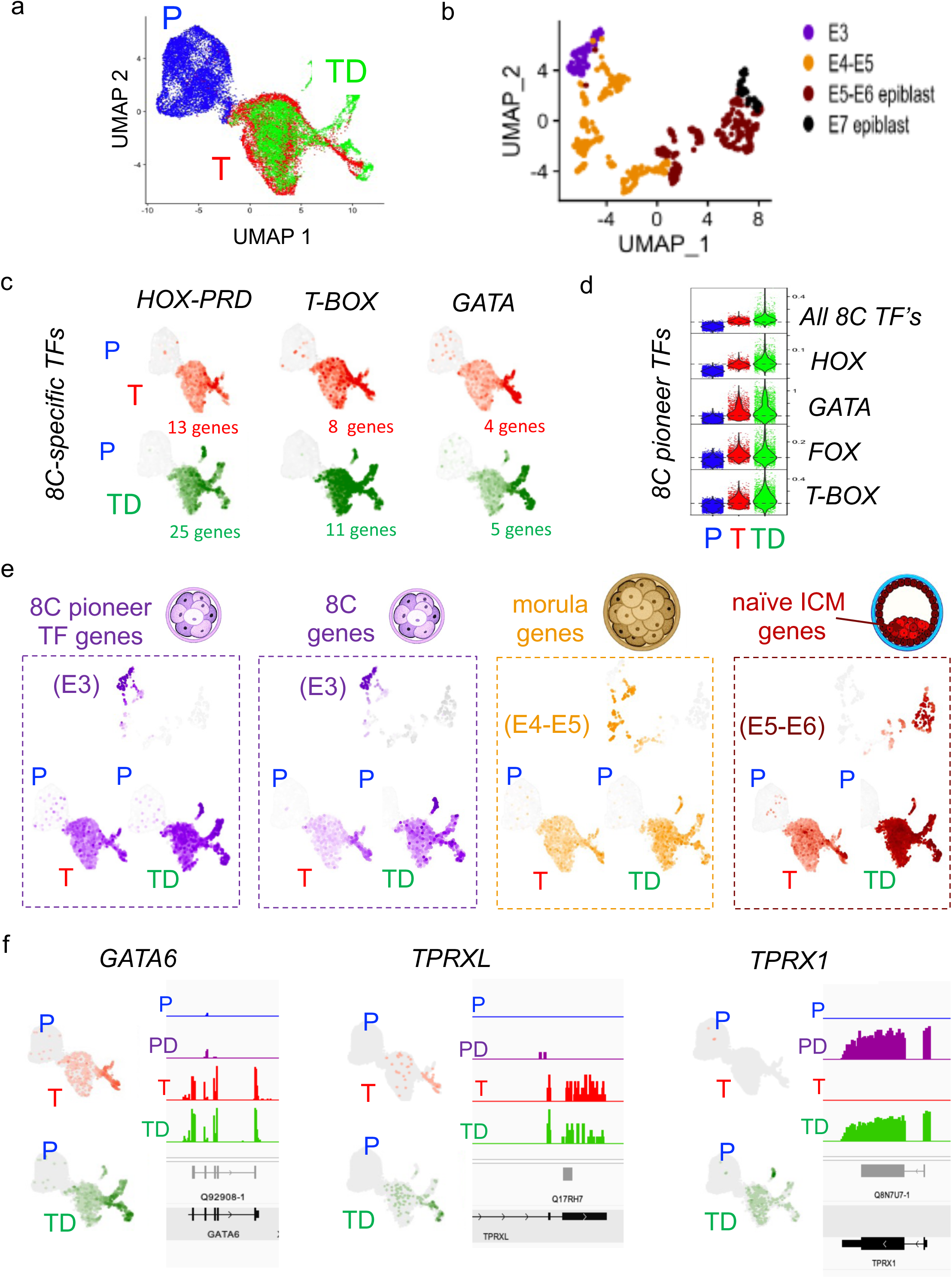
ScRNA-Seq analysis of TIRN-SC +/- iDUX4 activation. (**a**) UMAP plot of scRNA-Seq data for primed (P), TIRN (T) and TIRN+ iDUX4 (TD) cells showed major overlap between T and TD conditions. (**b**) UMAP plot of realigned (GRCh38) human embryo data (Petropoulos et al., 2016) with annotated E3-E7 stages, as shown. (**c**) UMAP plots showing average expression levels for HOX-PRD, T-BOX and GATA gene families that are overexpressed in TIRN (red) or TIRN+iDUX4 (green) cells. (**d**) Violin plots of single cell average expression levels for differentially expressed E3/8C specific pioneer HOX-PRD, GATA, FOX and T-BOX transcription factors families. (**e**) UMAP plots of indicated human embryo stages, P vs T and P vs TD for average expression levels of most differentially T/TD expressed (**Table S4**) 8C-specific transcription factor genes; (n=113), E3 (8C-specific; purple; (n=128 genes), E4-E5 (pre-lineage morula-specific genes; orange; n=134) and E5-E6 (naïve ICM-specific genes; maroon; n= 255 genes). (**f**) UMAP plot showing log-normalized corrected counts for selected DUX4 targets *GATA6*, *TPRXL,* and *TPRX1* in P vs T vs TD stem cells. Concordant bulk RNA-seq expression profiles show read signals at open reading frames for each gene in each condition.

To further map the developmental stage of single TIRN +/- iDUX4 cell gene expression, gene set modules (113-255 genes each) specifying 8C (E3), pre-lineage morula (E4-early E5), and naïve ICM (late E5-E6) embryonic stages(Petropoulos et al., 2016) were defined (**Fig. 3b,e, Table S4**). This strategy confirmed that single TIRN-SC homogenously co-expressed stage-specific pioneer factor genes that spanned human E3-early E5 pre-lineage human blastomere stages (**Fig. 3e**). The PRD-like homeobox *TPRX* genes (*i.e*., *TPRXL, TPRX1*) are DUX4 targets that are transiently translated before, during, and immediately following ZGA activation (4C-8C stages). Individual ectopic expressions of TPRXL and TPRX1 can activate ZGA-specific genes in hESC, and their knockdown stalls progression of ZGA in human embryos (Zou et al., 2022). Interestingly, a subset of TIRN-SC expressed the 4C-8C gene *TPRXL* (independent of iDUX4 activation) (**Figs. 3f; S3**). Additionally, although iDUX4 activation significantly augmented the global co-expressions of a large repertoire of 4C-8C-morula-specific pioneer factors in TIRN-SC (**Fig. 3e**), a subset of 8C-specific DUX4 targets (*e.g*., *TPRX1, ZSCAN4, LEUTX*) were homogenously expressed only in single TIRN + iDUX4 cells (**Figs. 3f; S3e,f; Table S4**). Collectively, these data suggested that iDUX4 had reinforced a more comprehensive totipotent 4C-8C molecular phenotype that was already expressed in unmodified single TIRN-SC.

### TIRN-SC functionally contributed embryonic and extra-embryonic lineages into developing human-murine chimeric feti, and generated human trophoblast stem cells (hTSC) with efficient *in vivo* placental chimerism

TIRN-SC possess higher *in vitro* directed differentiation potential (Park et al., 2020; Park et al., 2018; Zimmerlin et al., 2016), improved teratoma differentiation, and more efficient *in vivo* engraftment of differentiated progenitors (Park et al., 2020) than their isogenic, primed hPSC counterparts. We next investigated if TIRN-SC also possessed functional blastomere-like capacity to generate both embryonic and extraembryonic lineages in developing murine embryos. To track the *in vivo* developmental potential of TIRN-SC in human-murine interspecies chimeras, we transduced primed hPSC with stable lentivectors expressing cDNA for GFP/puromycin-resistance (puroR) and tdTomato/puroR. GFP/puroR-expressing primed hPSC were TIRN-reverted, and GFP^+^ TIRN-SC (>97% GFP^+^TRA-1-81^+^) were injected into E3.5 murine blastocysts and transferred into pseudopregnant foster females to generate human-mouse conceptuses (**Fig. S4a**). TIRN-SC-injected murine feti were harvested at E7.5, E9.5, E12.5, and E14.5 murine gestational time points. GFP^+^ mESC were also injected as controls in some experiments. Chimeric embryo cells were evaluated for human cell integration via multiple approaches: GFP expression by flow cytometry, presence of human mitochondrial genomic DNA sequence (Theunissen et al., 2016), expression of human GAPDH transcripts by sensitive qRT-PCR assay, expression of human-specific nuclear antigen (HNA) by immunofluorescence in fixed embryo sections, and *in vitro* expansion of puromycin-resistant human cells from whole embryo cultures.

Although 5-20% GFP^+^ cells could be detected within whole early post-primitive streak E7.5 murine embryo cells (which was comparable to murine GFP-mESC-injected controls), flow cytometry analysis revealed that TIRN-SC contributed to up to 30% of E7.5 human-mouse chimeras, and unlike mESC-injected control feti, GFP expression did not appear to be restricted to the epiblast (**Fig. S4b**). GFP^+^ expression dramatically diminished to below background at >E9.5 stages. Although the majority of GFP^+^TIRN-SC-injected murine blastocysts developed with normal fetal morphologies, 16%-18% of transferred embryos recovered at E9.5 and E14.5 stages were morphologically abnormal or growth-retarded. To better quantitate the extent of human chimerism at E9.5 and E14.5 stages, we analyzed a series of embryos using a sensitive human genomic mitochondrial DNA PCR assay (Theunissen et al., 2016). 22% of E9.5 embryos (*n*=50) and 27% E14.5 (*n*=18) embryos possessed human genomic DNA sequences at levels from 0.001% up to 1% (*i.e*., >1 human cell per 100-100,000 murine cells) (**Fig. 4a**). To confirm that genomic DNA results were recovered from live cells, RNA was also extracted from E9.5 chimeric embryos with positive DNA results and validated for expression of human-specific GAPDH by qRT-PCR (**Fig. 4b**). This analysis confirmed that 86% of E9.5 embryos (*n*=7) that were positive for human genomic DNA sequences also expressed human GAPDH transcripts at levels ranging from 1 human cell per ∼600-10,000 murine cells.

**Figure 4.**
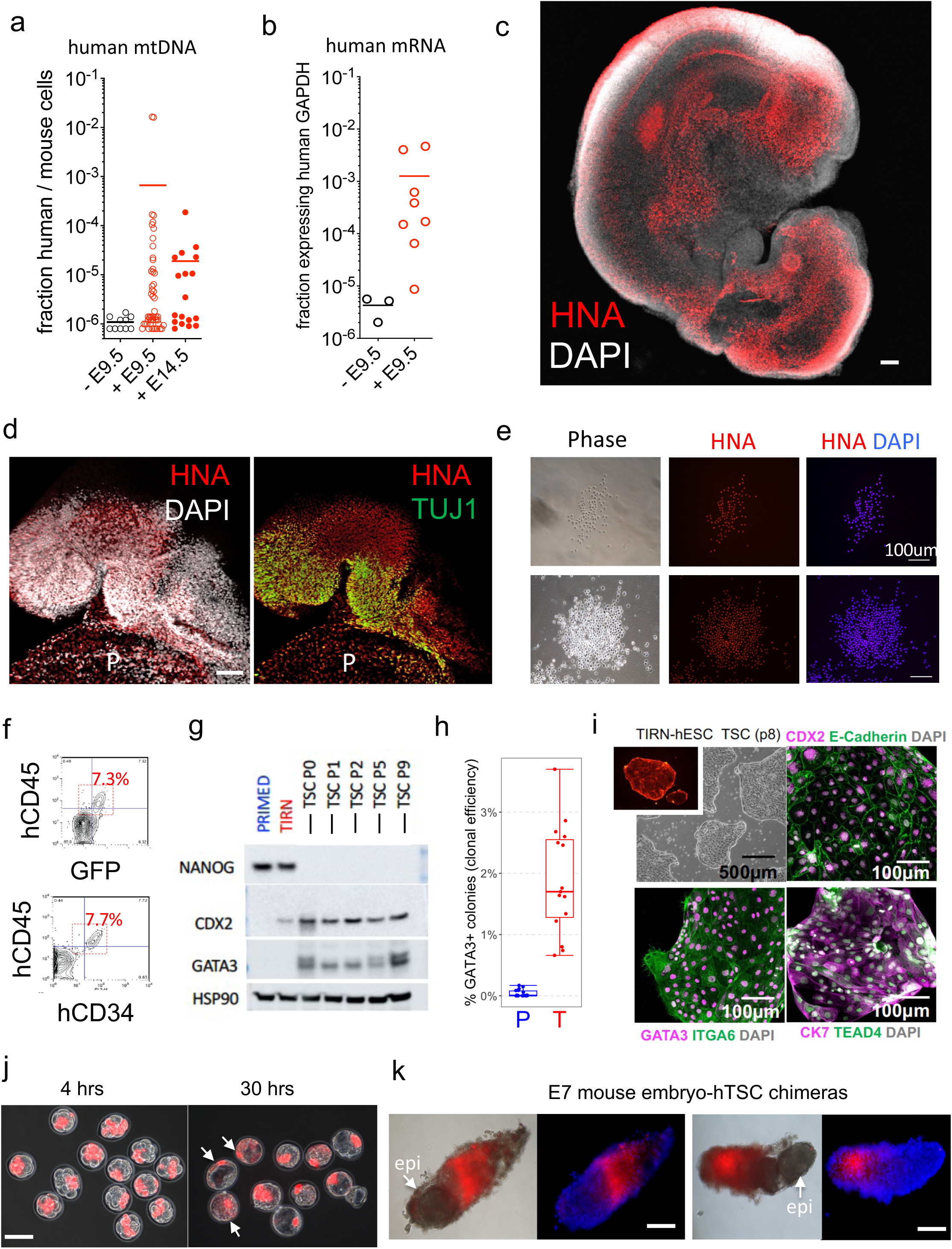
TIRN-SC contribution of human embryonic and extraembryonic lineages in human-murine chimeras, and derivation of human TSCs with *in vivo* trophectoderm potential. (**a**) Quantitation of human cell contribution from TIRN-SC-injected vs non-injected murine blastocysts in E9.5 and E14.5 human-murine fetal chimeras using human mitochondrial DNA-specific qPCR (Theunissen et al., 2016). (**b**) qRT-PCR quantification of human-specific GAPDH RNA transcript expression in human DNA positive E9.5 human-murine chimeric embryos and non-injected controls. (**c**) Whole mount immunofluorescent imaging of a representative E9.5 human-murine chimeric fetus showing detection of human (human nuclear antigen; HNA^+^)-expressing cells. Scale bar; 100 μm. (**d**) Immunofluorescent staining of HNA^+^β-Tubulin III (TUJ1)^+^ human neural cells in forebrain of E9.5 human-murine chimeric embryo. HNA+ cells were also detected sporadically in murine placenta (P). Scale bar; 100 μm. (**e**) Representative photomicrographs of HNA^+^ puromycin-resistant hematopoietic cells expanded and isolated from fetal livers of E12.5-E14.5 human-murine chimeric feti. (**f**) FACS analysis of GFP, human CD34 and human CD45 expression in puromycin-resistant *ex vivo* hematopoietic explants from human-murine chimeras. (**g**) Western blot analysis of NANOG, CDX2 and GATA3 protein expression in TIRN-SC and TIRN-derived TSC. Primed hPSC are shown as control. TIRN-hPSC co-express NANOG and low levels CDX2. TIRN-TSC rapidly adopt a NANOG^-^CDX2^+^GATA3^+^ phenotype. (**h**) Clonal efficiency of TIRN-TSC derivation during the first passage in TSC conditions. Compared to primed hPSC, TIRN stem cells efficiently produced GATA3^+^ TSC colonies (1-3%) upon switching to TSC conditions. (**i)** Immunofluorescent phenotyping of established TIRN-hESC-derived TSC. Representative colonies are shown after 8 passages and show uniform expression of CDX2, E-Cadherin, GATA3, ITGA6, CK7 and TEAD4. (**j**) Merged phase and fluorescent images of murine 8C embryos injected with tdTomato+ TIRN-derived TSC (arrows) following 4 and 30 hours in KSOM culture. Scale bars; 100 μm. (**k**) Phase and fluorescent photomicrographs of representative E7.5 mouse-human chimeras that resulted from injecting 5-10 td-Tomato+ human TSC in murine 8C embryos. Human TSC colonized exclusively the ectoplacental cone without evident engraftment in the murine epiblast (epi).

We next investigated if integration of human TIRN-SC had differentiated to specialized lineages within murine feti. Confocal microscopy of whole or cryo-sectioned E9.5 embryos revealed a fraction (38%; *n*= 21 embryos tested) with variably robust human-specific cell expression of HNA in somites, forebrain, spinal cord, eye, limb buds, heart bud, and extra-embryonic tissues (**Fig. 4c,d, Movie S1**); including neural-specific β-tubulin III^+^ TUJ1^+^HNA^+^ human cells integrated within murine prosencephalon, brain stem, and spinal cord, (**Fig. 4d**). Puromycin selection and expansion of chimeric E12.5 and E14.5 fetal liver (FL) or head explant tissues in neural or hematopoietic growth media allowed the selection and expansion of rare puroR HNA^+^ human cells in ∼50% of E12.5 chimeric embryos (*n*=6) (**Fig. S4c-e**). 15-30% of E12.5-E14.5 FL or neural cultures expanded low levels (2-8%) of puroR human hematopoietic (GFP^+^CD34^+^CD45^+^HLA-I^+^) or Neurofilament^+^ HNA^+^ cells (**Fig. S4c-f**). HNA^+^ human hematopoietic colonies were also isolated from puroR FL cells that co-expressed human CD34 and CD45 (**Fig. 4e,f**).

To evaluate the extra-embryonic potential of TIRN-SC, we next generated human trophoblast stem cells (hTSC) using published protocols established for primary placental tissue (Okae et al., 2018). TIRN cells generated self-renewing CDX2^+^GATA3^+^NANOG^-^ hTSC lines (**Figs. 4g-i, S4g**) at extremely high efficiencies (1-4%; **Fig. 4h**). Established hTSC could be passaged as self-renewing cell lines expressing TEAD4, E-Cadherin, ITGA6, and CK7 (**Fig. 4i**) and could specialize *in vitro* into SDC1^+^ syncytiotrophoblasts and HLA-G^+^ extravillous trophoblasts (**Fig. S4h**). To confirm the *in vivo* potential of TIRN-derived hTSC, single tdTomato^+^ TIRN-hTSC were injected into 4C-to-16C mouse embryos and transferred into pseudopregnant females. hTSC were rapidly integrated only into the trophectoderm of cultured blastocysts (**Fig. 4j**) and contributed specifically and efficiently only to the ectoplacental cone of E7 embryos *in vivo* (**Fig. 4k**). Collectively, these studies confirmed that TIRN-SC possessed both embryonic and extra-embryonic lineage differentiation potential *in vivo*.

### Single TIRN-SC with and without a murine E-Cadherin (mECad) transgene segregated *directly* to either embryonic or extra-embryonic compartments following injection into 8C-16C murine embryos

The simultaneous co-expression of naïve epiblast (*e.g*. *DNMT3L, NANOG, SOX2, POU5F1*), primitive endoderm (*e.g., FOXA2, GATA4, SOX17, GATA6*), and trophectoderm (*e.g., CDX2, GATA3, EOMES*) lineage genes (**Fig. S5a**) and proteins (**Fig. 5a,b**) in single TIRN-SC was similar to the concurrent expression of naïve epiblast, trophectoderm (TE), and primitive endoderm (PE) pioneer factors reported in 8C-to-32C (E3-E4) cleavage-stage human embryo cells (Petropoulos et al., 2016; Zou et al., 2022) and totipotent NANOG^+^GATA6^+^ and NANOG^+^CDX2^+^ 8C-to-32C murine blastomeres (Dietrich and Hiiragi, 2007; Maemura et al., 2021; Plusa et al., 2008). To test if TIRN-SC can functionally contribute to both embryonic (epiblast and PE) and extraembryonic (TE) lineages in a totipotent-like, single-cell manner, 5-6 single Hoechst-labeled TIRN-SC were injected into 8C-16C murine embryos and allowed to develop into blastocysts *in vitro* (**Fig. 5c-d**). To account for the possibility that interspecific differences in adhesion surface molecule interactions may be a barrier for early lineage segregation (Bao et al., 2022) that may inhibit efficiency of single cell human-murine engraftment, human TIRN-SC expressing a murine E-cadherin (mECad) transgene were prepared and injected in parallel (**Fig. S5b,c**). This approach permitted the direct quantitation of single human TIRN-SC +/- mECad into either murine trophectoderm or ICM via fluorescent microscopy. These confocal microscopic studies revealed that 5-6 Hoechst dye-labeled TIRN-SC were capable of colonizing and incorporating equipotently into *either* murine ICM or murine trophectoderm layers at 5-6 independently-integrated murine blastocyst sites at high single cell frequencies (**Fig. S5b,c**). Direct lineage segregation of injected TIRN-SC was further visualized directly by additional confocal microscopy that demonstrated distinct HNA^+^CDX2^+^GATA6^-^ TE-specific and HNA^+^GATA6^+^CDX2^-^ PE-specific of segregating clusters of cells in hatching human-murine chimeric blastocysts (**Fig. 5e, Movie S2**). However, we did not find significant differences in efficiency of murine ICM/trophectoderm blastocyst integration between TIRN-SC and TIRN-SC + transgenic mECad cells.

**Figure 5.**
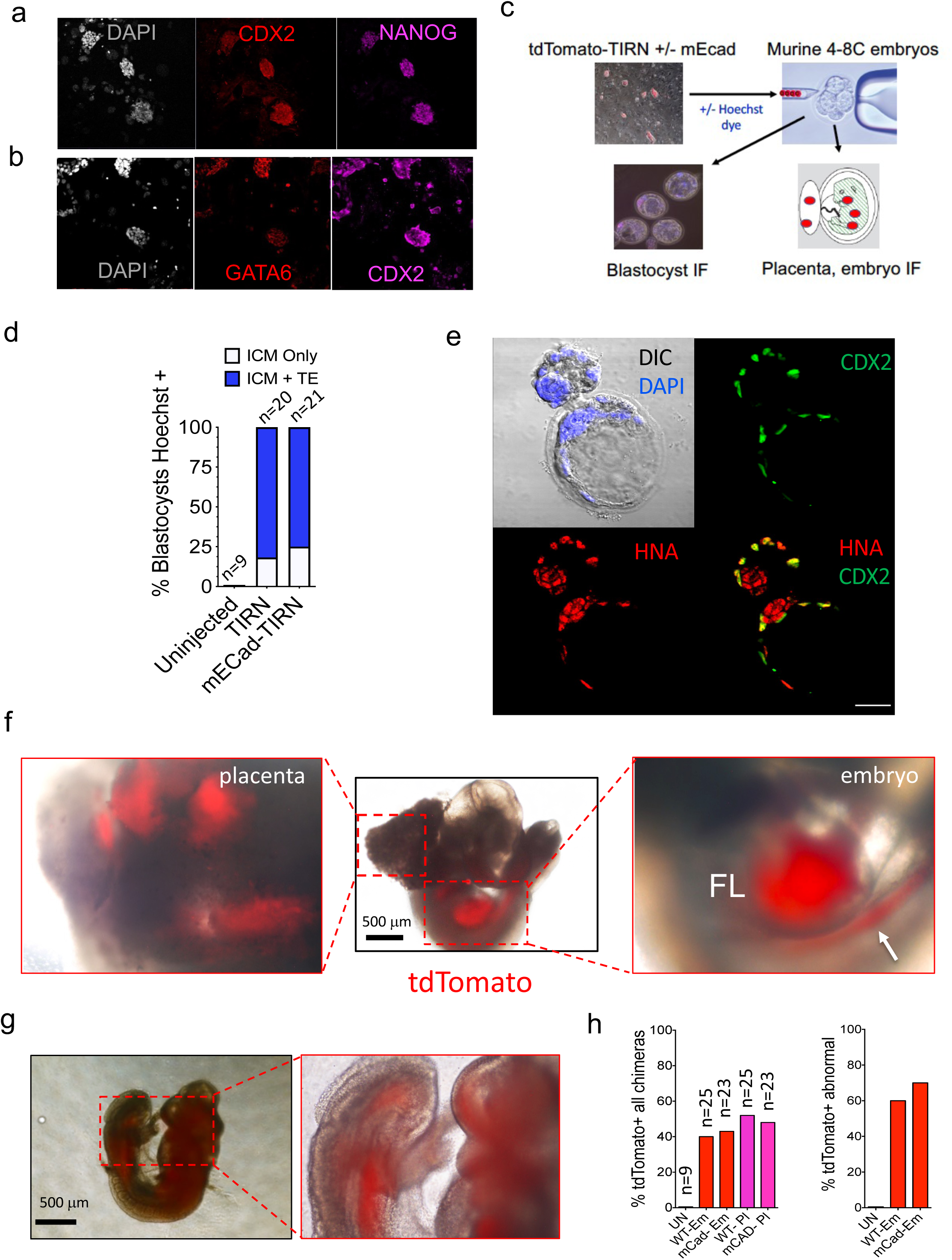
Direct single-cell contribution of TIRN-SC to both embryonic and extraembryonic lineages in human-murine blastocyst and fetal chimeras. Immunofluorescent cyto-staining of TIRN-SC showing simultaneous protein co-expression of (**a**) NANOG and CDX2 and (**b**) GATA6 and CDX2. (**c**) Experimental schema for testing direct lineage contribution of pre-lineage TIRN-SC with and without a murine E-Cad transgene. (**d**) Percentage of blastocysts that developed 24 hours following injection of 8C-16C murine embryos with 5-7 Hoechst-labeled TIRN +/- mECad cells with either ICM-only or both ICM and trophectoderm (ICM+TE) engraftments in chimeric blastocysts. (**e**) Confocal image of a hatching human-murine chimeric blastocyst ∼30 hours following injection of TIRN-SC into 4C-8C murine embryos showing merged (yellow) immunofluorescence of human HNA^+^ cells (red) in the ICM and hatching epiblast and CDX2^+^ in murine trophectoderm (green). (**f**) Fluorescent detection of TIRN-SC-derived tdTomato^+^ cells (red) in the extraembryonic placenta (left) and embryonic hematopoietic organs (fetal liver, FL; AGM region, arrow)) of a representative E11.5 human-murine chimeric embryo. (**g**) Robust fluorescent detection of tdTomato^+^ cells throughout a human-murine chimeric fetus shows broad contribution to rostral nervous system, thoracic and caudal structures. (**h**) quantitation of TIRN-SC-derived tdTomato+ cells in chimeric embryos (Em) and placentae (P).

Finally, to determine if single TIRN-SC were competent in contributing *equipotently* to both embryonic and extra-embryonic tissues in developing post-implantation chimeric conceptuses *in vivo*, we injected 5-6 single tdTomato^+^ TIRN-SC +/- mEcad cells into murine 8C-16C embryos. Chimeric blastocysts were transferred into females for implantation and development, and human cell integration in E11.5 chimeric feti was scored via tdTomato expression in either embryonic or placental tissues. These experiments detected robust tdTomato chimerism within both embryos and placentae, including robust human cell colonization of fetal liver and (aorta-gonad-mesonephros) AGM organs, but with little difference in chimera efficiency between control TIRN-SC and TIRN-SC + mECad cells (**Figs. 5g-I; S5c,d**). The human origin of tdTomato^+^ cells in murine placentae was further validated by immunostain detection of HLA-G and human placenta lactogen (hPL) antigens (**Fig. S5d**). These results demonstrating *direct* segregation of both embryonic and extra-embryonic lineages *in vivo* from single TIRN cells was suggestive (but not confirmatory in this surrogate interspecies assay) of totipotent-like functionality of TIRN-SC. It was also notable that although multilineage integration of human TIRN-SC +/- mECad cells in murine embryos was relatively efficient, the majority (∼60-70%) of tdTomato^+^ chimeric feti did not develop normally (**Figs. 5i; S5e**).

### TIRN-SC were diminished in *both* TNKS and PARP1 catalytic activities, displayed perturbed TNKS/PARP1 protein levels, and expressed hundreds of PARP1-regulated developmental genes

We next sought to understand the role XAV939 might be playing in reprogramming primed hPSC to a blastomere-like state. Interestingly, TIRN-SC had *increased* TNKS1/2 and *decreased* PARP1 protein levels relative to primed hPSC (**Table S2**), thus potentially resetting a TNKS/PARP1 protein ratio equilibrium. Previous studies showed that XAV939 treatment of murine 1C embryos arrested TNKS-dependent development at the ZGA/totipotent 2C stage (Gambini et al., 2020). Furthermore, PARP1-deficient mESC epigenetically de-repressed expression of a large cohort of multi-lineage-specifying pioneer factor genes (*e.g., Gata4*, *Gata6, Cdx2, Hand1, Mixl1*) (Liu and Kraus, 2017). A meta-analysis of published RNA-seq data from PARP^-/-^ mESC (Liu and Kraus, 2017) revealed that XAV939-inhibited TIRN-SC expressed >650 of the same developmental lineage-specifying genes over-expressed in PARP-deficient mESC (**Table S3**). Thus, we postulated that continuous culture at micromolar concentrations (*i.e*.., 4 μM) of XAV939 in the TIRN system non-specifically affected *both* TNKS1/2 and PARP1 protein stabilities; potentially resulting in (premature) transcriptional activation of PARP1-regulated lineage-specifying genes, in a manner similar to PARP1-deficient mESC (Liu and Kraus, 2017).

To confirm a role for XAV939-mediated PARP protein level perturbations, we evaluated the expressions, activities, and targets of *both* PARP1 and TNKS1/2 in TIRN-SC. As expected in XAV939-inhibited cells (Callow et al., 2011; Huang et al., 2009; Zimmerlin and Zambidis, 2020), TIRN-SC stabilized protein expressions of known TNKS substrates (*e.g*., TNKS1/2, AXIN1, and ANGIOMOTIN), and modulated subcellular distributions of the active, non-phosphorylated isoform of β-catenin as reported (Kim et al., 2013) (**Fig. 6a,b**); consistent with known XAV939-TNKS-mediated PARdU effects on these targets. To test the impact of XAV939 on PARP1-mediated activities, undifferentiated and meso-endoderm-differentiated cell lysates from primed vs TIRN-SC were bound to ADP-ribose-binding Af1521 macrodomain resin beads. Anti-poly-ADP ribosylation (PAR) and anti-mono-ADP-ribosylation (MAR), and PARP1 Western blotting were performed on enriched lysates. These studies revealed not only that TNKS1/2 and PARP1 were predominately non-ADP-ribosylated, but also demonstrated dramatic reductions of *all* PAR and MAR protein activities in undifferentiated TIRN-SC that persisted for up to 96 hours following initiating differentiation (**Fig. 6c,d**). Furthermore, unlike primed hPSC, TIRN-SC lysates exhibited reduced total and ADP-ribosylated-PARP1 protein levels throughout differentiation; in a manner similar, albeit not as completely, as PARP1-deficient mESC (Liu and Kraus, 2017).

**Figure 6.**
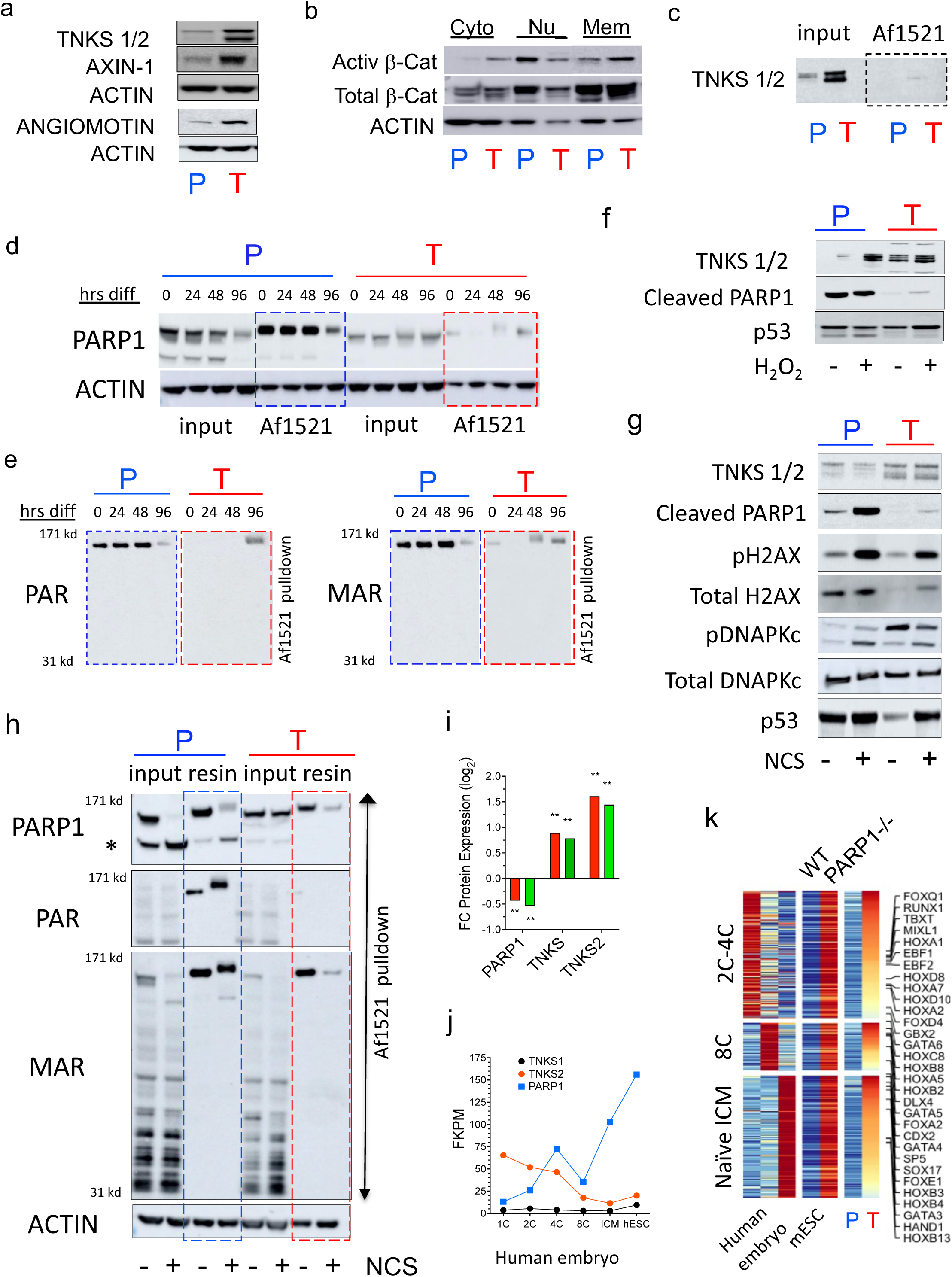
Dual inhibition of both TNKS and PARP1 activities in TIRN-SC. (a) Western blots of PARdU-regulated TNKS protein substrates: TNKS1/2, AXIN-1, and ANGIOMOTIN in primed (P) vs isogenic TIRN (T) cells. (**b**) Western blots of total and activated (non-phosphorylated) β-catenin in (P) vs isogenic (T) cells within cytoplasmic (cyto), nuclear (nu) and membrane (mem) fractions. (**c**) Western blots of input control and Af1521 resin pull-down lysates for TNKS1/2 protein expression in (P) and (T) cells. (**d-e**) Western blots of lysates from input control and Af1521 macrodomain resin pull-downs in (P) vs (T) undifferentiated (0 hours) cells and at indicated timepoints following meso-endodermal differentiation (24, 48, 96 hours). Blots show total input, and ADP-ribosylated (Af1521 pull-down; dashed line boxes) and probing with anti-PARP1, anti-PAR, and anti-MAR antibodies (Poly- and mono-ADP-ribosylations; PAR, MAR, respectively). PAR, MAR are dramatically suppressed in TIRN-SC (red dashed line box) compared to primed controls (blue dashed line box). (**f**) Western blot analysis of DNA damage/oxidative stress responders: TNKS1/2, cleaved PARP1, and p53 proteins following oxidative stress conditions (H_2_O_2_ treatment) in (P) vs isogenic (T) cells. (**g**) Western blot analysis of TNKS1/2, cleaved PARP1, p53 and DNA damage and repair-associated proteins in (P) vs isogenic (T) cells; with or without treatment with the radiomimetic DNA-damaging chemical NCS. (**h**) Western blots of Af1521 resin pulldowns of all ADP-ribosylated proteins in lysates of (P) vs isogenic (T) cells; with or without treatment with NCS. Input controls are shown. PAR, MAR are dramatically suppressed in TIRN-SC (red dashed line box) compared to primed controls (blue dashed line box). Blots revealed both ADP-ribosylated and non-ADP-ribosylated full length forms of PARP1, and its cleaved form (*). Notably, TIRN-SC were reduced in detectable cleaved PARP1 (* lower band). (**i**). Relative TIRN/primed fold changes (FC) in protein expressions of indicated PARP proteins in primed (P; blue) vs TIRN (T; red) stem cells from proteomics data (** p<0.001; n=3 replicates). **(j**) Protein expression of PARPs during human early embryogenesis (translatome datasets (Zou et al., 2022)) relative to hESC lines. **(k)** Heatmap of expression meta-analysis of PARP1^-/-^ vs WT mESC (Liu and Kraus, 2017) and (T) vs (P) from RNA-seq data; showing the intersection of human and orthologous mouse genes overexpressed (FC>1.5) in both TIRN-SC and PARP ^-/-^ mESC, and correlation to human embryo translatome datasets (Zou et al., 2022) (**Table S3**).

To validate that dual PARP1/TNKS perturbations impacted not only the protein levels, but also the functionality of both PARPs, we performed Western blot analysis of DNA damage response (DDR) proteins in TIRN vs primed cells following oxidative (H_2_O_2_) and radiomimetic DNA-damaging (neocarzinostatin; NCS) stress conditions (when TNKS and PARP1-mediated PAR activities are normally upregulated). These studies revealed that under these stress conditions (**Fig. 6f-h**), XAV939-inhibited TIRN cells responded with reinforced TNKS protein expression, while restraining upregulation of DDR-activated cleaved PARP1 levels. Moreover, PAR- and MAR-modified protein levels were globally decreased in TIRN-SC in DDR conditions, including PARP1 (self)-PARylation (Ko and Ren, 2012) (**Fig. 6h**). Despite suppression of *both* PARP1- and TNKS-mediated PAR catalysis, XAV939-inhibited TIRN-SC were still able to activate protein expression of genome-stabilizing DDR machinery (*e.g*., phosphorylated histone H2AX, p53, and the non-homologous end joining (NHEJ) protein DNAPKc). Collectively, these results suggested that blastomere-like reprogramming of TIRN-SC may be driven by a XAV939-perturbed TNKS/PARP1 protein ratio expression disequilibrium of *increased TNKS* and *decreased PARP1* protein levels relative to primed hPSC; which curiously also resembled the patterns of protein expression levels of these PARPs and their targets during 2C-8C stage embryogenesis (**Fig. 6i-k)**.

### TIRN-SC possessed DUX4-accessible totipotent-like enhancer regions co-occupied by NANOG-SOX2-OCT4 and PARP1 (NSOP)

PARP1 is a nucleosome-binding protein that cooperates with core pluripotency factors to regulate lineage-specific developmental programs (Liu and Kraus, 2017). Since zebrafish studies have assigned a master role for core pluripotency factors in the initiation of ZGA (Gao et al., 2022; Lee et al., 2013; Riesle et al., 2023), we postulated that the TIRN blastomere-like state may be driven by PARP1-mediated epigenetic reprogramming of NSO factors. To investigate the epigenetic consequences of a perturbed PARP expression equilibrium on NSO-regulated gene expression, we performed TIRN vs primed cell ChIP-Seq on PARP1, NANOG, SOX2, and POU5F1 (OCT4) factors, as well as histone regulatory marks (*i.e*., H3K4me3, H3K27me3, H3K27ac). These studies revealed significant genome-wide reorganizations of PARP1, NANOG, SOX2, and OCT4 chromatin binding in TIRN vs primed cells (**Fig. S6a-e**). Consistent with relative cellular PARP1 depletion, genome-wide PARP1 enrichment in TIRN-SC was globally decreased relative to primed hPSC (**Fig. S6a,f**), whilst the total number of NANOG and SOX2 binding sites were significantly increased genome-wide (**Fig. S6b,c**). As reported in mESC (Liu and Kraus, 2017), PARP1 co-bound with NANOG, SOX2, and OCT4 individually as a quartet (NSOP) in both primed and TIRN cells (**Fig. 7a S6f**). An integration of the 4692 gained NSOP co-binding sites in TIRN-SC with published DUX4 ChIP-Seq data in hPSC (Hendrickson et al., 2017) revealed that, in contrast to primed hPSC, reprogrammed TIRN-SC NSOP sites were relocated and centered at chromatin regions accessible to DUX4 co-binding (**Fig. 7**).

**Figure 7.**
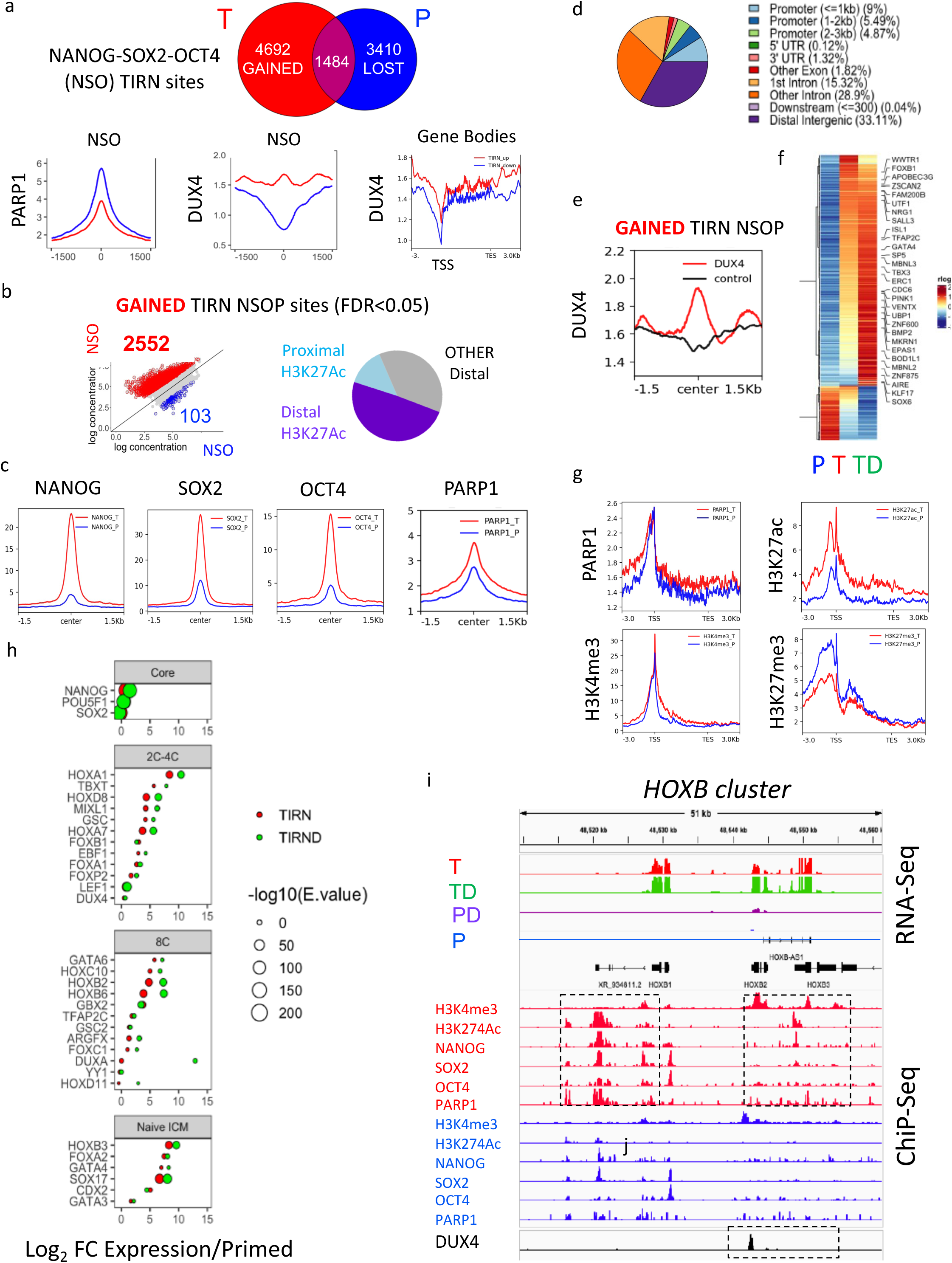
ChIP-Seq analysis of DUX4-accessible totipotent-like chromatin enhancer regions that co-bind NSO and PARP1. (**a)** (Top) Euler diagram showing number of overlapping shared (purple), gained (red), and lost (blue) NANOG-SOX2-OCT4 (NSO) co-binding sites in TIRN (red) vs primed (blue) cells. (Bottom) Distribution histograms of average signal for PARP1 and DUX4 at co-bound TIRN NSO gained (red line) vs primed NSO lost (blue line) sites (recentered and normalized with DiffBind software, DUX4 signal obtained from hg38 realigned transgenic DUX4 datasets (Hendrickson et al., 2017)); averaged DUX4 signal is also shown at NSO sites within gene bodies (TSS: transcriptional start sites) of genes differentially overexpressed in TIRN-SC (red), or primed hPSC (blue). (**b**) Scatter plot shows averaged log-transformed concentrations (n=2 replicates per factor) at NSO co-bound sites (2552 differentially bound sites with highest confidence (FC>2, FDR<0.05) are colored for TIRN gained (red) and primed lost (blue) sites). Pie chart shows percentage distribution of these 2552 sites co-bound by H3K27ac at <3kb of TSS (proximal enhancer), or >3kb of TSS (distal enhancer). (**c**) Distribution histograms show averaged, normalized NANOG, SOX2, OCT4, and PARP1 read signals (red), centered on non-overlapping NSO TIRN-gained peaks (FC>2, FDR<0.05) vs same locations in primed (blue) samples. These significant 2552 TIRN-gained NSO sites possessed enriched PARP1 deposition, predominated at proximal and distal H3K27ac-enriched enhancer regions, and were located at (**d**) intronic and distal intergenic chromatin. (**e**) Profile plots of averaged DUX4 ChIP-Seq binding scores centered at the 2552 TIRN-gained NSO sites (red) vs control DUX4-ChIP signal (black) (Hendrickson et al., 2017). Data is centered on NSO binding sites identified in adjoining plots. (**f**) Heatmaps of RNA-seq data for differentially expressed genes bound within 10 kb of TSS at TIRN-gained NSO-DUX4 regions (FC>2, FDR<0.05). Select embryonic transcription factors are indicated. (**g**) Line plots show average ChIP-Seq signal for PARP1, H3K4me3, H3K27me3 and H3K27ac at gene bodies with NSO TIRN-gained sites (FC>2, FDR<0.05) in TIRN-SC and primed samples **(h)** Dot plot of top enriched motifs (CentriMo) in NSO gained sites (FC>2, FDR<0.05) ranked by expression level showing -log_10_ E-value (adjusted p-value using the binomial test multiplied by the number of identified motifs) with correlative relative expression (log_2_ fold change) for genes expressed in TIRN (red) and TIRN+iDUX4 (green) cells vs primed controls. (**i**) Integrative Genomics Viewer snapshots of representative TIRN NSOP-associated *HOXB* cluster RNA-seq and ChIP-Seq tracks showing RNA expression signal in (T), (TD), (PD) and (P) samples and the averaged input-subtracted RPGC-normalized binding of H3K4me3, H3K27ac, NANOG, SOX2, OCT4 and PARP1 in TIRN (red) and primed (blue) samples. Averaged, and genome-aligned DUX4 signal from hg38 realigned DUX4-ChiP data (Hendrickson et al., 2017) is also shown (black). Boxed regions highlight novel TIRN NSOP-co-binding putative enhancer regions (enriched for H3K27ac and H3K4me3).

Interestingly, the most differentially bound (FDR<0.05; FC>2) NSOP regions in TIRN-SC defined a cohort of 2552 DUX4-accessible sites enriched for H3K27ac-occupied proximal (<3 kb from TSS) and distal (>3 kb from TSS) enhancers (**Fig. 7b-e**). Unlike the relative genome-wide reduction of PARP1 binding in TIRN vs primed cells, these NSOP-DUX4 TIRN regions were *paradoxically enriched* with PARP1 binding. To explore how these DUX4-NSOP regions might control gene expression outcomes, we examined transcript expressions from gene promoters and bodies within 10 kb of these NSOP enhancer sites with our RNA-seq data from TIRN +/- iDUX4 cell lines (**Figs 7f; S6g**). Promoters and gene bodies associated with these NSOP regions were transcriptionally active (with increased PARP1, H3K4me3, H3K27ac, and decreased H3K27me3 enrichments) (**Fig. 7g, Table S6**). These expressed genes included many of the 2C-4C, 8C, and naïve ICM lineage-specifying factors described above, and their expressions were further augmented following iDUX4 activation.

We next performed *de novo* sequence motif analysis of these DUX4-accessible NSOP TIRN-SC regions. This analysis revealed significant motif enrichment for *combinatorial co-binding* of hundreds of multilineage, developmentally critical pioneer transcription factors predicted to regulate ZGA, epiblast, trophectoderm, primitive and definitive endoderm, ectoderm, and mesoderm lineage specifications (*e.g*., *HOX, FOX, GATA, SOX, TBX, CDX, DUX* families; **Fig. 7h**) (Goke et al., 2011; Liu et al., 2019; Riesle et al., 2023). The predicted motifs of factors in these DUX4-accessible TIRN NSOP regions mirrored the lineage-specifying 4C-8C-naïve ICM pioneer factors that were already co-expressed in hybrid fashion in single TIRN-SC (**Fig. 1C, Table S1**), including EBF2, GATA6, and HOXB cluster genes (**Fig. 7i, Table S6**). Collectively, these results exposed a genome-wide remodeling of NSOP enhancer sites in TIRN-SC that resided in DUX4-accessible regions, and that putatively regulated a pioneer factor-driven, multilineage (embryonic and extraembryonic), totipotent-like transcriptional program.

### TNKS/PARP1 substrate perturbations drove a global reprogramming of the ubiquitinome in TIRN-SC

The preceding studies revealed an unexpected epigenetic reprogramming of core lineage-regulating factors in PAR-deficient TIRN-SC. In trying to understand how dual TNKS/PARP1 perturbations were able to reprogram the epigenome, we postulated a likely mechanism was that global shutdown of cellular ADP-ribosylation and decreased PARP1 levels mediated a profound impact on the PARdU-dependent, ubiquitin-modified epigenetic machinery (*e.g.*, PAR-dependent histones, ubiquitin ligases, and transcriptional regulators). Since post-translational ubiquitination of histones and core pluripotency factors alters DNA binding (Cui et al., 2018; Fang et al., 2014a; Kim et al., 2014; Pei, 2017; Wang et al., 2016), we postulated that PARP-regulated ubiquitination was the driving force in NSOP-associated epigenetic reprogramming to a blastomere-like state.

To uncover a role for a reprogrammed ADP-ribosylome that in turn drove a reprogrammed ubiquitinome, we paired whole proteome with ubiquitinome studies in isogenic primed and PAR-depleted TIRN +/- iDUX4 cells and employed a database of known PARP1 and TNKS1/2 substrates (Ayyappan et al., 2021) to help interpret results. TIRN and TIRN+iDUX4 cell proteomics revealed similar patterns of decreased PARP1, increased TNKS, and 8C/morula-specific proteins. Proteomic GSEA revealed abundant developmental pathways that were driven by differential expression of TNKS and PARP1 protein substrates (**Figs. 8a,b; S8**). Moreover, differential TIRN +/- DUX4 proteomics were primarily driven by protein levels of known PARP1/TNKS targets that displayed discordance with mRNA expression, suggesting a major role for UPS-driven post-transcriptional gene regulation (**Fig. 8b**). Remarkably, an analysis of the whole ubiquitinome of primed vs TIRN +/- iDUX4 cells revealed a hyper-ubiquitinated proteome in TIRN-SC relative to primed hPSC (**Fig. 8c,d**). This hyper-ubiquitinated proteomic signature was almost *entirely* driven by differential ubiquitination of known PARP1/TNKS substrate targets (**Fig. S9a,b**), suggesting a mechanism involving the reprogramming of the ubiquitination/deubiquitination machinery. Indeed, an analysis of differential protein expression of PAR-dependent ubiquitin modifying enzymes revealed a global reprogramming of TNKS/PARP-targeted E3 ligases and DUBs in TIRN-SC (**Fig. S9c, d**). An analysis of TIRN vs primed cells revealed significant differential expressions of other ubiquitin-modifying enzymes (*e.g*., NEDD4, RING1, DTX3L, USP9X) that regulate and reshape chromatin, or that modify the transcription factors PARP1, SOX2, OCT4, and NANOG (*e.g*., WWP2, TRIM32) (**Fig. 8e**),

**Figure 8.**
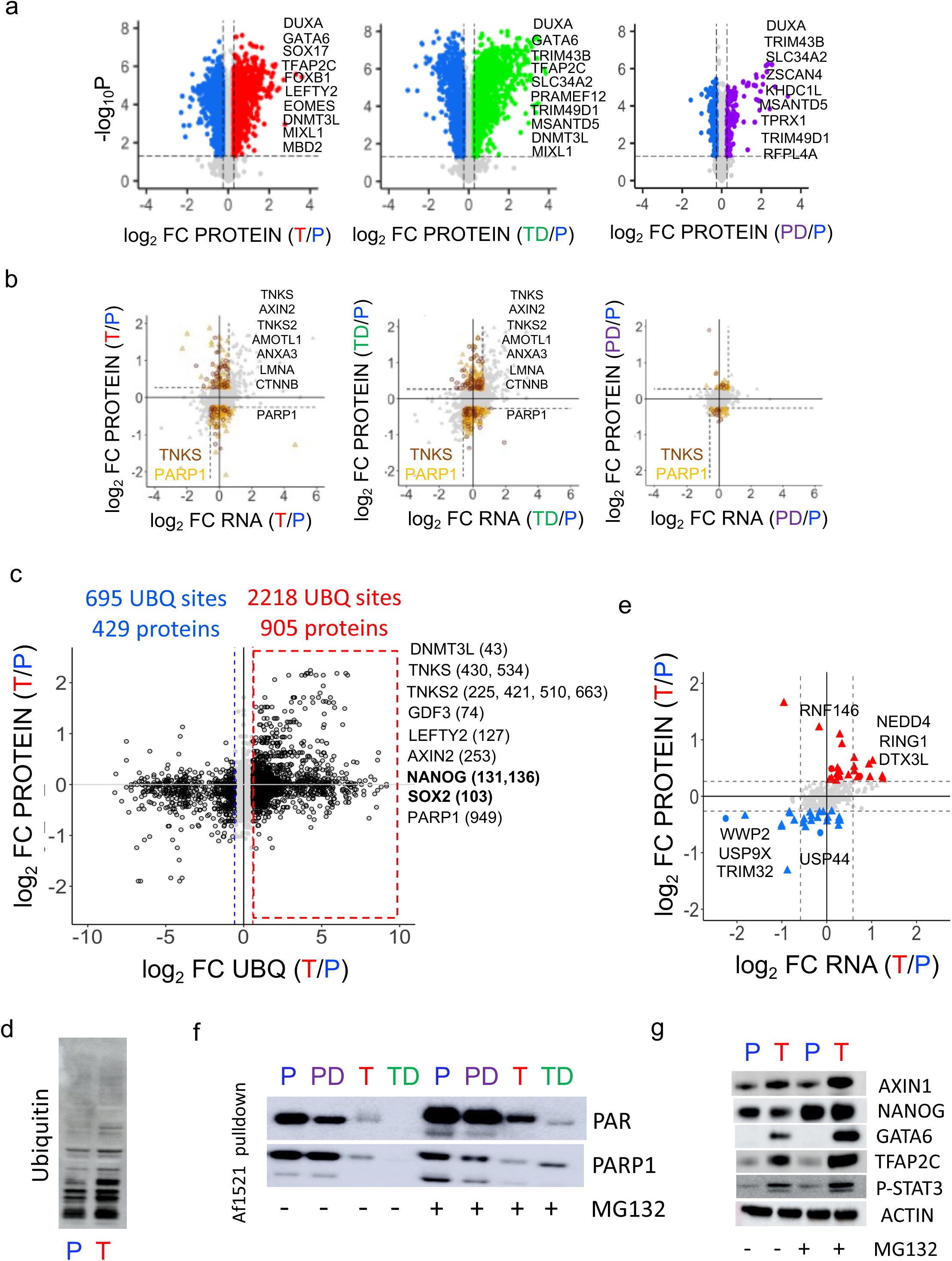
PARP1/TNKS protein substrate expressions and reprogramming of the ubiquitinome in TIRN-SC. (**a**) Volcano plots show global proteome expressions of differentially expressed proteins (fold change > 1.2, p < 0.05, dashed lines) in TIRN (red), TIRN + iDUX4 (green) and primed+iDUX4 (purple) vs primed (blue) hPSC. Selected upregulated cleavage stage protein factors are listed. (**b**) Cross plots of log2 fold change (FC) of RNA-seq (x-axis; RNA) vs whole proteome (y-axis; PROTEIN) data showing non-correlated (dashed lines) RNA/protein expressions of known TNKS and PARP substrates (protein FC>1.2, RNA FC < -1.5 or protein FC < -1.2, RNA FC > 1.5) in TIRN, TIRN+iDUX4 and primed+iDUX4 cells relative to primed controls. Representative proteins are indicated. Both TIRN-SC conditions differentially express proteins with non-correlative RNA expression, suggesting post-translational stabilization or degradation mechanisms. (**c**) Crossplot of ubiquitinome sites (x-axis UBQ) vs proteome (y-axis PROTEIN) relative expressions in TIRN (T) vs primed (P) cells show the most differentially ubiquitinated protein sites (dashed lines, FC>1.5). Hyper-ubiquitinated proteins and lysine sites of representative proteins in TIRN-SC are indicated. (**d**) Western blot analysis of total ubiquitinated proteins in primed (P) and TIRN (T) lysates. (**e**) Cross plots of log2 fold change (FC) of RNA-seq (x-axis; RNA) vs whole proteome (y-axis; PROTEIN) data showing expression of ubiquitin machinery proteins. Significantly expressed proteins (FC>1.2, p<0.05) are indicated in red (TIRN) and blue (primed). (**f**) Western blot of Af1521 pull-down enriched PAR and ADP-ribosylated PARP1 in primed (P) primed + iDUX4 (PD), TIRN (T) and TIRN + IDUX4 (TD) cells with (dashed box), or without MG132-mediated proteasomal inhibition. PAR levels in TIRN-SC augment with MG132 revealed a role of the UPS in their reduced PARylation. (**g**) Western blot of representative proteins in primed (P) and TIRN-SC (T) with or without MG132 proteasomal inhibition.

Furthermore, we detected differential polyubiquitination of key developmental proteins (e.g., DNMT3L, TNKS, LEFTY2, GDF3, AXIN2, PARP1, SOX2, and NANOG) in TIRN +/- iDUX4 cells; with potential impacts on protein conformation or UPS-regulated degradation of these factors (especially when protein expression was discordant with mRNA expression). To validate that the mass discordance of mRNA expression with protein expression levels of differentially ubiquitinated TNKS/PARP1 protein substrates (**Fig. 8b**) was due to UPS regulation, we validated selected differentially ubiquitinated protein targets with MG132 proteosome inhibitor studies in TIRN-SC. These MG132 experiments confirmed proteosome-mediated regulation of expressions of key stem cell proteins (e.g., PARP1, AXIN2, NANOG, GATA6, TFAP2C, and pSTAT3, **Fig. 8f,g**). Interestingly, NANOG and SOX2 were differentially over-ubiquitinated at or near their DNA-binding domains; potentially altering their DNA binding (**Movie S3**). Finally, we found protein expression level shifts in TIRN-SC of differentially ubiquitinated histones, histone modifiers, ubiquitin ligases, and deubiquitinases (**Fig. S9d-f**). Collectively, these data revealed a broad post-translational proteomic reprogramming of the epigenetic landscape of TIRN-SC that was likely driven by a XAV939-mediated reprogramming of PARdU-dependent ubiquitinated transcriptional machinery.

## DISCUSSION

Understanding the molecular events that occur at the first cell divisions of human embryogenesis greatly impacts our knowledge of human ontogeny and reproduction. TIRN-SC clonally co-expressed large cohorts of homeobox transcription factors that were previously only predicted at the transcript level for the earliest human embryonic stages (Kuliev et al., 1996; Verlinsky et al., 1995). Because development kinetics between human and mouse or zebrafish embryos are widely divergent (Thomas et al., 2021), access to human reference data has become essential to accurately evaluate the developmental congruity of putative *in vitro* analogs. The recent availability of 2C-8C-morula human embryo single cell RNA-seq (Petropoulos et al., 2016) and translatome Ribo-seq (Zou et al., 2022) datasets provided new tools to validate the reprogrammed human blastomere proteo-transcriptomic signatures we discovered in TIRN-SC. Several decades ago, human embryonic stem cells (hESC) (Thomson et al., 1998) and human induced pluripotent stem cells (hiPSC) (Takahashi et al., 2007) were introduced as *in vitro* models of epiblast-stage embryos for studying human development, drug discovery and regenerative medicine. TIRN-SC lines can now provide a cell culture model for elucidating the proteogenomic remodeling that orchestrates human ZGA and totipotent stem cell biology.

In human embryos, ZGA is ignited by master pioneer transcription factors that are specifically and momentarily expressed (*e.g*., DUX4 (Hendrickson et al., 2017; Vuoristo et al., 2022) and TPRXL/1/2 (Zou et al., 2022)). We demonstrated that TIRN-SC activated hundreds of 4C-8C DUX4 targets (including TPRXL) and putative 2C-4C maternal pre-ZGA reprogramming factors with a blastomere-like program that was further reinforced by transgenic iDUX4 overexpression. We also identified expression of a limited repertoire of 8C-specific DUX4 targets (e.g., *TPRX1, LEUTX*) that required exogeneous, high level iDUX4 expression for their activation. Since *TPRXL* was sufficient to activate ZGA-specific genes in hESC (Zou et al., 2022), it is currently unclear if the additional DUX4 targets activated by iDUX4 are redundant or self-regulating for priming of the totipotent phenotype *in vivo*. Rare 8C-like cells were recently described within heterogeneous naïve hESC (Mazid et al., 2022; Taubenschmid-Stowers et al., 2022) and analogous 8C-like populations can be induced by DUX4 overexpression or chemical manipulation (Yu et al., 2022); albeit functional differentiation of these cells was not demonstrated as we have demonstrated herein for TIRN-SC. Additionally, unlike TIRN-SC, these 8C-like cells were not homogenous in their expression of blastomere-stage genes and expressed a more limited repertoire of ZGA-associated DUX4 targets (*e.g*., *LEUTX* and *TPRX1*), that closer resembled our primed hPSC+iDUX4 controls than TIRN+iDUX4 cells. Furthermore, TIRN-SC over-expressed 2C-4C candidate maternal reprogramming factors (e.g. HOX cluster genes), and other 8-cell/morula-specific genes implicated in activation of human ZGA (*e.g*., ZNF675 and LSM1; **Table S1**); which may overlap with ZGA-associated maternal OCT4 activation (rather than DUX4) in 8C human embryos (Yuan et al., 2023). Overall, our TIRN +/- iDUX4 cell model is consistent with the previously suggested notion that DUX4 primes and enhances, but does not initiate or is required, for the activation of ZGA (De Iaco et al., 2020; Vuoristo et al., 2022).

Importantly, TIRN-SC co-expressed a hybrid transcriptional program of pioneer factors that uniquely mimicked expression profiles in human 8C and cleavage-stage cells, and that we propose is regulated by PARP1 via an epigenetic DUX4-NANOG-SOX2-OCT4 axis. Co-expression of lineage-specifying factors along with core pluripotency factors is a feature not only in 8C-morula stage human embryos (Petropoulos et al., 2016; Zou et al., 2022) but also translates across pre-lineage cells of disparate species (Farrell et al., 2018; Lee et al., 2013; Redo-Riveiro et al., 2024; Thompson et al., 2022). In zebrafish and mice, diverse cell types arise from a pool of equipotent common stem-progenitors that similarly express combinatorial transcription factor modules of multiple fates, but that can subsequently trans-specify from one fate to another along epigenetically plastic lineage bifurcations. For example, although primitive endoderm (PE)-specifying GATA6 expression is lost in mESC and hESC (Warrier et al., 2018), forced NANOG and GATA6 expression demonstrated a capacity to co-bind shared enhancer regions to promote ICM plasticity. Similarly, cooperative epiblast specific SOX2 and PE GATA6 enhancer region co-binding was detected in a subset of nave hESC and cleavage stage blastomeres (Redo-Riveiro et al., 2024).

In human and mouse ESC, NANOG-SOX2-OCT4 bind promoters and enhancers of circuitry that both activate and repress expression of target genes (including homeodomain proteins) with downstream self-renewal or differentiation outcomes (Ai et al., 2022; Boyer et al., 2005; Buecker et al., 2014; Heurtier et al., 2019; Kim et al., 2008). This study discovered novel TIRN-specific NANOG-SOX2-OCT4 and PARP1 co-binding proximal and distal enhancer sites at DUX4-accessible, H3K27ac-bound chromatin that appear to regulate the expressions of hundreds of lineage-specifying factors. While TIRN-SC were characterized by global, genome-wide loss of PARP1 binding (correlating to lower PARP1 protein expression), this cohort of 2552 novel gained NSO regulatory sites in TIRN-SC chromatin were paradoxically *increased in enrichment* of PARP1 co-binding, and were associated with 2C-4C, 8C, and pre-lineage naïve ICM genes that were transcriptionally activated in TIRN-SC by increased H3K27ac and H3K4me3, and decreased H3K27me3 co-deposition at their gene bodies. Remarkably, these NSOP enhancer sites contained co-binding motifs for hundreds of the same 2C-4C, 8C, and pre-lineage naïve ICM lineage-specific pioneer factors that were already expressed in TIRN-SC; potentially defining a broad feedback loop that putatively provides an epigenetic and transcriptional blueprint for post-totipotent embryonic and extraembryonic lineage development. The assignment of zebrafish paralogs of NANOG-SOX2-OCT4 at the top of a hierarchy of maternal reprogramming factors that ignite ZGA (Gao et al., 2022; Lee et al., 2013; Riesle et al., 2023) collectively highlights that: 1) core pluripotency factors may be critical in the transcriptional regulation of human cleavage-stage embryonic cells, and 2) the deployment of core pluripotency factors at enhancer regions may involve a cooperative partnership with hundreds of extra-embryonic and embryonic lineage-specifying pioneers factors that ultimately execute the outcomes of human totipotency.

Our studies also revealed, to our knowledge, the first example that a global perturbation of PARP protein expressions and catalytic activities (along with LIF-2i chemical inhibition) can mediate proteogenomic reprogramming *in reverse* from one developmental state (pluripotent) to another (blastomere). TIRN-SC displayed: 1) global loss of all ADP-ribosylation activities, 2) *increased* protein levels of TNKS1/2, and 3) *decreased* protein levels of PARP1; in a manner that mimicked the protein expression ratios of these PARPs during 2C-8C human development. In contrast, primed, undifferentiated hPSC expressed robust PAR/MAR activities, including PARylated levels of PARP1 protein both before and after differentiation. Since TIRN-SC and PARP^-/-^ mESC both upregulated over 650 identical multilineage and cleavage-stage lineage pioneer factors, we propose a central role for PARP1 in activating the observed simultaneous expression of disparate, and antagonistic multi-lineage (mesoderm/endoderm/ectoderm) pioneer factors, and we postulate that the relative loss of PARP1 in XAV939-inihibited TIRN-SC likely activated their blastomere-like pre-lineage state. PARP1 was identified as a member of the OCT4-SOX2 interactome which was highly expressed in a subset of 32-64 cell preimplantation mouse embryos that resembled the mESC 2i naïve state (Gerovska and Arauzo-Bravo, 2019). Furthermore, PARP1’s regulation of SOX2 binding (Liu and Kraus, 2017), and its capacity to occupy and protect the promoters of Nanog, Pou5f1, Sox2, and Dppa3 from becoming epigenetically repressed (Roper et al., 2014), supports our proposal that PARP1 likely plays a master role in ZGA-associated epigenetic reprogramming *in vivo*; likely via a PARP1-DUX4 axis. However, the mechanism of a dual PARP1-TNKS-mediated reprogramming currently remains enigmatic in our studies, but *likely synergistic*, since TNKS has also been independently implicated in regulating ZGA activation (Gambini et al., 2020).

Indeed, we propose a critical and general role for *synergistic* TNKS/PARP1-mediated cellular ADP-ribosylation, PARdU, and ubiquitination for regulating the totipotent stem cell state. ADP-ribosylation is a central post-translational modification (PTM) that directly influences other critical PTMs (*e.g*., ubiquitination, methylation, acetylation, phosphorylation, and SUMOylation) that in turn collectively regulate stem cell transcription, chromatin structure, and cell signaling. For instance, PARP1 catalytic activity controls H3K4me3 methylation and transcription through PARylating KDM5B (Krishnakumar and Kraus, 2010). Ubiquitination also regulates chromatin/histone architecture (Mattiroli and Penengo, 2021; Oss-Ronen et al., 2022; Vaughan et al., 2021), transcription factor binding and stability, and the ubiquitin machinery itself (Rona et al., 2018; Vittal et al., 2015). Furthermore, TNKS-mediated PARdU is tightly intertwined with ubiquitination of AXIN1-mediated WNT signaling via regulation of PAR-dependent ubiquitin ligase RNF146, that requires PAR for activation (Callow et al., 2011; Cho-Park and Steller, 2013; Rona et al., 2018; Vittal et al., 2015; Vivelo et al., 2019). Other PAR-regulated ligases and deubiquitinating enzymes conversely regulate the stability and degradation of epigenome-regulating pluripotency factors, including PARP1, SOX2 (Cui et al., 2018), NANOG (Liu et al., 2016), and OCT4. We demonstrated that PAR-deficient TIRN-SC vastly reprogrammed their ubiquitinome and that this differential ubiquitinated repertoire correlated directly with shifts in protein levels of ubiquitin writers and erasers, histone modifiers, histones, and transcriptional regulators that are known TNKS and PARP1 substrates. Accordingly, histone ubiquitination controls transcriptional activity (Oss-Ronen et al., 2022) and genome stability (Mattiroli and Penengo, 2021). Because NANOG-SOX2-OCT4 regulate chromatin accessibility and gene expression during ZGA in the zebrafish (Gao et al., 2022; Lee et al., 2013; Miao et al., 2022) and possibly human cells (Gao et al., 2018; Yuan et al., 2023), and because PARP1 may regulate the binding and transcription of NSO via effects on ubiquitination (Liu and Kraus, 2017; Roper et al., 2014), we investigated and indeed found differentially ubiquitinated sites in both PARP1 and at the DNA binding domains of NANOG and SOX2, and the implications of these findings is yet to be discerned. Our studies herein should liberate future investigations on the mechanisms behind PARP-regulated proteogenomic reprogramming of developmental states.

Finally, we demonstrated that TIRN-SC possess functional extra-embryonic and embryonic differentiation potential *in vivo* via interspecific chimera assays. TIRN-SC generated TSC with high fetal placental contribution, and following injection into 8C stage mouse embryos, single TIRN-SC populated both the ICM and the trophectoderm of the mouse blastocyst. The level of contribution of stem cells to embryonic chimeras is dependent on the developmental potency and stage of both the donor cells and the recipient embryo. Complete totipotency extends only to single blastomeres of 4-cell stage mouse embryos (Maemura et al., 2021), and unbalanced chimeric contribution can be achieved by subsequent cleavage cells (8-cell to 32-cell pre-ICM stages). Due to ethical constraints surrounding the generation of chimeras with human embryos, functional validation of these various human naïve pluripotent states has been performed only in interspecies chimera assays. Compared to the relatively robust interspecific chimerism demonstrated between rat ESC and mouse embryos (especially within the context of blastocyst complementation strategies) (Kobayashi et al., 2010; Wu et al., 2017; Yamaguchi et al., 2017), thus far, various hPSC have contributed only modest amounts of differentiated human cells into human-mouse (Gafni et al., 2013; Theunissen et al., 2016; Theunissen et al., 2014), human-pig, human-sheep (Wu et al., 2017) and human-monkey chimeras (Fang et al., 2014b).

### Limitations of the Study

Overall, our functional studies validate a putative but not definitive totipotent-like capacity of TIRN-SC. The most stringent demonstration of totipotency would require that a single TIRN cell could give rise to an entire human conceptus, which is an ethically unacceptable test. Alternative functional validation of TIRN-SC totipotency by injection of single cells into human 8C embryos with tracking of development to embryonic and extraembryonic lineages in chimeric human feti is also not ethically conceivable. However, herein, we successfully tracked individual Hoechst^+^ or tdTomato-labeled human TIRN cells following injection into 8-16-cell murine embryos and found that ∼5-7 TIRN-SC could proportionally and equipotently contribute to ∼5-7 independent murine blastocyst ICM or trophectoderm implantations, and ultimately gave rise to *both* embryonic and placental lineages *in vivo*. However, chimeric human-murine feti with the highest human TIRN-SC contributions also developed abnormally or rapidly degenerated in a mouse uterus by E14.5. Several xenobarriers may account for this observation; and that ultimately impede the measurement of single-cell human totipotent cell contributions of TIRN-SC in species-disparate 2C-8C animal embryos. These barriers include disparate developmental kinetics between human TIRN-SC and recipient murine embryos (Thomas et al., 2021), as well as non-optimal cell-cell communications between human blastomere-like cells developing within species-incompatible embryonic niches. Although murine E-cadherin-modified TIRN-SC appeared to better target human cell integration into murine neural, FL, and AGM organs, efficient interspecific crosstalk will ultimately require not only sophisticated genetic strategies of interspecific embryo complementation, but also potentially cross-species adhesion molecule and cell signaling engineering.

## ACKNOWLEDGEMENTS

We acknowledge support of the JHUSM Proteomics core and shared instrumentation grant S10OD021844, and the Wilmer Core Grant for Vision Research (EY001765). We are grateful to Chip Hawkins and the Johns Hopkins Transgenic Mouse Core Facility for technical support with murine embryo injections and uterine transfers. This work was supported by grants from the NIH/NEI (R01EY032113; ETZ), The Maryland Stem Cell Research Fund (2023-MSCRFV-5995; 2024-MSCRFV-6248; ETZ), and The Lisa Dean Moseley Foundation (ETZ). We are grateful for technical contributions from Dr. Willem Buys, Moritz Becker, Andre Vu, Yunlong He, and Xiaoyuan Chen.

## AUTHOR CONTRIBUTIONS

ETZ and LZ co-wrote the manuscript. LZ, AAM, TSP, REM, JT, SY, IU, IV, CK, WB, AKLL, and ETZ designed, performed, and/or analyzed all experiments. LZ wrote code for and analyzed all bioinformatics data. All authors interpreted results and edited the manuscript. ETZ conceived, supervised, procured funding for, and directed the project. ETZ and LZ approved the final manuscript.

## COMPETING INTERESTS

The authors declare no competing interests.

## SUPPLEMENTAL INFORMATION

Accompanies this manuscript at the following weblink:

## DATA AVAILABILITY

The NIH Gene Expression Omnibus has issued the accession number, for RNA-Seq data, Chip-Seq data in this manuscript. The GEO-supplied link for access will be released on publication.

## * STAR METHODS

## RESOURCE AVAILABILITY

**1. Lead Contact.** Further information and requests for resources and reagents should be directed to and will be fulfilled by the lead contact, Elias T. Zambidis, MD, PhD.
**2. Materials Availability.** All unique/stable reagents and cell lines generated in this study are available from the Lead Contact without restriction.
**3. Data and Code Availability.** Data: Single cell and bulk RNA sequencing, and ChIP-Seq data in this manuscript are available through Gene Expression Omnibus at GEO with accession numbers: TBA. NIH and ArrayExpress GEO accession numbers for previously published datasets are listed and catalogued in Table S7 (*e.g*., Zou et al, Petropoulos, Liu, Kraus, Zimmerlin et al data), and include: GSE141639 (bulk RNA-seq of primed and TIRN-SC; Park et al, 2020), E-MTAB-3929 (scRNA-Seq of human embryos; Petropoulos et al, 2016), GSE197265 (Ribo-seq and RNA-seq of human embryos; Zou et al, 2022), GSE74111 (RNA-seq of Parp1KO mESC (Liu and Kraus, 2017, GSE95515 (DUX4 ChIP-Seq in hiPSC; Hendrickson et al, 2017). Microscopy and behavior data reported in this paper will be shared by Elias T. Zambidis upon request. Code: R and Python code used to analyze single cell datasets will be available on GitHub upon publication. Any additional information required to reanalyze the data reported in this paper is available from Elias T. Zambidis upon request.
**4. Key Resources Table** (Table S7).
**5. Methods Details**

### Bioethics

All hPSC experiments conformed to the guidelines published by the National Academy of Sciences, and the International Society of Stem Cell Research (ISSCR). The origins of all hiPSC and hESC used in this study are summarized in **Table S7**. hESC lines were purchased from the Wisconsin International Stem Cell Bank (WISCB) and the Rockefeller University (RU). hiPSC lines were previously described (Zimmerlin et al, 2016, Park et al, 2018, 2020). All experiments were conducted under purview of the Johns Hopkins University (JHU) Institutional Stem Cell Research Oversight (ISCRO) and conform to Institutional standards regarding informed consent and provenance evaluation. The use of WISCB- and RU-donated hESC lines in these studies was approved by the Johns Hopkins University (JHU) Institutional Stem Cell Research Oversight (ISCRO) Committee and conformed strictly to JHU Institutional standards regarding written informed consent and provenance evaluation. All human-murine embryo chimera studies received approval by the JHU ISCRO committee prior to initiation of experiments. The use of Rockefeller University hESC cell lines RUES1 and RUES2 used for injection into murine embryos was approved by the JHU ISCRO to be consistent with the provenance of the original hESC derivation protocol where donors consented to the mixing of xenogeneic embryos with hESC. Additionally, all animal use and surgical procedures were performed in accordance with protocols approved by the Johns Hopkins School of Medicine Institute of Animal Care and Use Committee (IACUC). All experiments involving animal procedures (*e.g.,* subcutaneous/intra-muscular injections of NOD.Cg-Prkdcscid Il2rgtm1Wjl/SzJ (NOG SCID) mice for teratoma analysis and blastocyst transfer into ICR pseudopregnant mice) were reviewed and approved by the Johns Hopkins University Animal Care and Use committee. These reviews included considerations for ethical sacrifice, humane housing, and appropriate measures to minimize animal discomfort. All mice (C57BL/6J, ICR and NOG SCID) used in this study were purchased from The Jackson Laboratory.

### Conventional, primed (E8) and TIRN (LIF-3i) cultures of hESC and hiPSC

All hESC and hiPSC lines used in these studies were maintained and expanded either in conventional Essential 8 (E8) medium or chemically reverted with the LIF5i -> LIF-3i TIRN method (**Fig. S1a**), as previously described (Zimmerlin et al, 2016, Park et al, 2018, 2020). Conventional, primed cultures were propagated using an in-house E8 medium formulation consisting of DMEM/F-12 supplemented with 2.5 mM L-Glutamine, 15 mM HEPES and 14 mM sodium bicarbonate (ThermoFisher Scientific, cat# 11330), 50-100 ng/mL recombinant human FGF-basic (Peprotech), 2 ng/mL recombinant human TGF-β1 (Peprotech), 64 μg/mL L-ascorbic acid-2-phosphate magnesium (Sigma), 14ng/mL sodium selenite (Sigma), 10.7 μg/mL recombinant human transferrin (Sigma), and 20 μg/mL recombinant human insulin (Peprotech). Primed hPSC were cultured in E8 on Vitronectin XF (STEMCELL Technologies) matrix-coated tissue culture-treated 6-well plates (Corning) and passaged every 5-6 days by non-enzymatic reagents (*i.e*., Versene solution (ThermoFisher Scientific) or Phosphate-Buffer-Saline (PBS)-based enzyme-free cell dissociation buffer (ThermoFisher Scientific, #13151). TIRN-reverted stem cell cultures were stably propagated following LIF-5i adaptation, as described (Zimmerlin et al, 2016, Park et al, 2018), using ‘LIF-3i’ medium, which consists of DMEM/F-12 supplemented with 20% KnockOut Serum Replacement (KOSR, ThermoFisher Scientific), 0.1 mM MEM non-essential amino acids (MEM NEAA, ThermoFisher Scientific), 1mM L-Glutamine (ThermoFisher Scientific), 0.1 mM β-mercaptoethanol (Sigma), 20ng/mL recombinant human LIF (Peprotech), 3 µM CHIR99021 (Peprotech), 1µM PD0325901 (Sigma or Peprotech), and 4 µM XAV939 (Peprotech). As mentioned, TIRN-SC were first chemically-reverted from isogenic, primed E8 medium-expanded primed hPSC cultures using a single adaptation step in ‘LIF-5i’ medium (*i.e*., LIF-3i medium supplemented with 10µM Forskolin (Peprotech), 2µM purmorphamine (Peprotech) and 10ng/mL recombinant human FGF2 (Peprotech)), as described (Zimmerlin et al, 2016, Park et al, 2018, 2020). Briefly, 4-to-5-day-old primed hPSC E8 cultures were switched to LIF-5i +FGF2 medium for 12-24 hours, enzymatically dissociated (Accutase, ThermoFisher Scientific), and transferred onto irradiated mouse embryonic fibroblast (MEF) feeders in LIF-5i medium for only 3 to 5 days. All subsequent passages were grown in LIF-3i (TIRN) medium alone on MEF feeders and passaged as single cells using Accutase. Isogenic E8 (primed) vs LIF-3i (TIRN) cultures were maintained in parallel passage number for simultaneous phenotypic characterization, as previously described (Zimmerlin et al, 2016, Park et al, 2018, 2020). Unless otherwise indicated, parallel isogenic hPSC samples were prepared for all downstream analytical analyses following at least 5-7 passages in TIRN conditions, by collecting cells using a 1:1 Versene/Accutase mixture (5min at 37°C) and passing through a 100 μm cell strainer. All hPSC samples used in analytical RNA/genomic/protein studies were first quality-verified for >90-95% cellular viability (Trypan blue) before sample collections and were also verified by flow cytometry for undifferentiated cell identity (*e.g*., >95% TRA-1-81^+^SSEA4^+^ expression).

### Transgenic stem cell lines

The pCW57.1-DUX4-CA lentivector plasmid (Jagganathan et al, 2016) was purchased from Addgene (#99281); packaged and purified lentiviral particles were produced by SignaGen Laboratories. pCW57.1-DUX4-CA lentiviral particles were transduced into primed RUES2. The CMV-Luciferase-EF1α-copGFP-T2A-Puro BLIV 2.0 Lentivector (SBI, BLIV513VA-1) was transduced into primed RUES1. A PiggyBac transposable vector expressing murine E-cadherin with mCherry fusion (Addgene 71366) was created by using the Gateway recombination system and pAC150-PBLHL-4xHS-EF1a-DEST (Addgene 48234) destination vector and was integrated into primed RUES2. The LV-EF1a-tdTOMATO-IRES-NEO (SignaGen Laboratory, SL100201) tdTomato expression vector was transduced into primed RUES2 and RUES2-mEcadherin transduced lines. Stable transfectants were selected by supplementing E8 medium with 0.5 μg/mL puromycin or 100 μg/mL hygromycin continuously. Stable clones were validated and selected based on GFP/tdTomato expression by fluorescent microscopic and flow cytometric analysis.

### Antibodies

Source and working dilutions of all antibodies used in these studies for Western blots, FACS, ChIP-Seq, and immunofluorescence experiments are listed in **Table S7.**

### Flow Cytometry and immunofluorescence of stem cell lines

#### Sample preparation and flow cytometry of cell surface marker expressions

Primed and TIRN-SC were collected with Versene/Accutase (1:1) to diminish MEF feeder carryover in the LIF-3i condition. Cells were washed once with PBS, filtered through a 70μm cell strainer to discriminate against doublets and counted with the Countess cell counter. For each staining, single cell suspensions (<1×10^6^ cells in 100 µL) were incubated for 20 min on ice with directly conjugated mouse monoclonal anti-human antibodies or the appropriate isotype controls using the antibody volumes listed in **Table S7**. Antibodies included mouse monoclonal anti-human CD24-PE (BD Biosciences), CD77-FITC (BD Biosciences, #551353), CD90-PE (BD Biosciences, # 555596), HLA-ABC-APC (BD Biosciences, # 555555), SSEA4-APC (R&D Systems, # FAB1435A), TRA-1-81-PE (BD Biosciences, # 560161), and TRA-1-81-Alexa Fluor 647 (BD Biosciences, #560793). Stained cells were washed with 3mL PBS and filtered through a 40µm cell strainer prior to acquisition using a FACSCalibur flow cytometer (BD Biosciences) and the BD CellQuest Pro analytical software (BD Biosciences) or a Beckman Coulter Cytoflex flow cytometer. The FlowJo analysis software (Tree Star v10) was used for offline data analysis and Figure preparation. Cell debris and aggregates were excluded from the analysis based on their light scatter profile.

#### Immunofluorescence staining on chamber slides and culture plates

Isogenic hPSC cultures and TIRN-SC-derived TSC were passaged at 30,000 cells per cm^2^ and expanded for 4-6 days in 8-well Lab-Tek II chamber slides (Nunc, Thermo-Fisher Scientific) in their respective media onto Vitronectin-XF matrix (primed E8), over a mouse feeder monolayer (TIRN LIF-3i/MEF) or 0.1% gelatin (TSC). For embryo explant cultures, see embryo culture explant section for culture plate and sample preparation. Cells were washed with PBS, fixed using 2% formaldehyde solution in PBS (Affymetrix) for 15 minutes at 4°C, washed with PBS and maintained in sterile PBS at 4°C until immunostaining. All samples were incubated for one hour in blocking solution (either PBS, 5% goat serum (Sigma) and 0.05% Tween 20 (Sigma) or TBS, 0.1% Triton X-100 (Sigma), 5% goat serum (Sigma)). All antibody dilutions were performed in their corresponding blocking solution to reduce non-specific binding. Cells were incubated overnight at 4°C in humid chambers with primary unconjugated antibodies at indicated dilutions (**Table S7**). The next day, the slides were washed 3 times for 5 minutes (1X Dako Wash buffer or TBS-T) and incubated for 1 hour at room temperature with the diluted Alexa488, Alexa555 or Alexa647 highly cross-adsorbed secondary antibody corresponding to the primary antibody host species. Cells were washed 3 times. For dual-antibody staining, samples were incubated with a second primary antibody that was either directly conjugated or raised from a distinct species (i.e., rabbit vs. mouse) for 2 hours at room temperature, wells were washed 3 times for 5 minutes in Dako wash buffer, and, if the second primary antibody was unconjugated, incubated for 1 hour at room temperature with a diluted highly cross-adsorbed secondary antibody corresponding to the second primary antibody host species. Cell nuclei were stained with 10µg/mL 4′,6-diamidino-2-phenylindole (DAPI, ThermoFisher Scientific) for 5-10 minutes at room temperature. Cells were washed 3 times with PBS, the wells were separated from the slide, and the slides were mounted with a coverslip using Prolong Gold or Glass Antifade Reagent (ThermoFisher Scientific). The mounting reagent was left to cure overnight at room temperature in the dark. Immunostains were imaged with a Nikon Eclipse TE 2000-U or using a Zeiss LSM 510 Meta Confocal Microscope (Zeiss) at the Wilmer Eye Institute, Johns Hopkins University’s imaging facility. Isotype controls for mouse (ThermoFisher Scientific) and rabbit (Dako) IgG replaced corresponding primary antibodies at matching concentrations as negative controls and were imaged using identical settings.

### Reverse-transcription Polymerase Chain Reaction (RT-PCR)

The sequences and references for all PCR primers and Taqman probe sets used in these studies are catalogued in **Table S7**.

#### Quantitative Real-Time Polymerase Chain Reaction (qRT-PCR)

Isogenic primed and TIRN-SC were simultaneously prepared and collected with Versene-Accutase, as described above. Additionally, primed and TIRN RUES02 cells expressing transgenic DUX4 were prepared after 24 hours supplementation with doxycycline. Total RNA was extracted from snap-frozen cell pellets using the RNeasy Mini Kit (Qiagen) according to the manufacturer’s protocol. DNase digestion (Qiagen) was performed in-column to eliminate genomic DNA. Total RNA concentration and purity were measured using a NanoDrop spectrophotometer (ThermoFisher Scientific). For each sample, 2μg of RNA was reverse transcribed with the SuperScript VILO cDNA Synthesis Kit (ThermoFisher Scientific) according to the manufacturer’s protocol and a MasterCycler EPGradient thermocycler (Eppendorf). Diluted cDNA samples (either 1:10 or 1:20 in nuclease-free water) were admixed either with TaqMan gene expression assays (ThermoFisher Scientific, see **Table S7** for primer sequences) and TaqMan Fast Advanced Master Mix (ThermoFisher Scientific) or with relevant primers (**Table S7**) and Power SYBR Green Master Mix (ThermoFisher Scientific) for quantitative real-time PCR amplification using the ViAA7 Real Time PCR System (ThermoFisher Scientific). Relative gene expression was calculated using the ΔΔCt method, using ACTB expression assay primed samples as controls.

For time-course molecular analyses, feeder-dependent hPSC cultures were MEF-depleted by pre-plating onto gelatin-coated plates for 1 hour at 37°C as previously described (Park et al, 2018). Samples were sequentially and simultaneously collected from 3 representative hPSC lines in primed conditions (10ng/mL FGF2), after adaptation for 24 hours in LIF-5i, at the 1^st^ passage in LIF-5i (p1) and at the 3^rd^ or 4^th^ passages in LIF-3i (p>3). Total RNA was isolated from snap-frozen samples using the RNeasy Mini Kit (Qiagen) following the manufacturer’s instructions and quantified using a Nanodrop spectrophotometer (ThermoFisher Scientific). Genomic DNA was eliminated by in-column DNase (Qiagen) digestion. Reverse transcription of RNA (1µg/sample) was accomplished using the SuperScript VILO cDNA Synthesis Kit (ThermoFisher Scientific) and a MasterCycler EPgradient (Eppendorf). For real-time PCR amplification, diluted (1:20) cDNA samples were admixed to the TaqMan Fast Advanced Master Mix (ThermoFisher Scientific) and the following Taqman gene expression assays (ThermoFisher Scientific): DNMT3L (Hs01081364_m1), GAPDH (Hs99999905_m1), KLF2 (Hs00360439_g1), NANOG (Hs02387400_g1), and NR5A2 (Hs00187067_m1). For analysis of HERV-H expression, diluted (1:50) cDNA samples were amplified using the Power SYBR Green Master Mix (ThermoFisher Scientific) and the following primers: GAPDH forward GGTCATCCATGACAACTTTGG and reverse ACAGTCTTCTGGGTGGCAGT, HERV-H pol forward CGCCCTTCTTCCCAATCCAA and reverse GCCAAGGAGGGAGTAGAGGT. Samples were amplified in technical triplicates and real-time fluorescence detection was achieved using a ViAA7 Real Time PCR System (ThermoFisher Scientific). Relative quantification of gene expression for each time point was normalized to GAPDH and isogenetically compared to starting primed samples using the ΔΔCt method.

#### Non-Quantitative RT-PCR

Collection and freezing of cell pellets, RNA extraction, DNase digestion, and total RNA quantification were performed, as described above. Briefly, 2μg of total RNA was reverse transcribed using the SuperScript VILO cDNA Synthesis Kit (ThermoFisher Scientific), according to the manufacturer’s protocol, and thermo-cycling was performed in a MasterCycler EPGradient (Eppendorf). For the PCR amplification of the DUX4 Transgene and GAPDH, 1μL of 1:10 diluted cDNA was mixed with Platinum Taq DNA Polymerase (Invitrogen), 10X PCR Buffer - MgCl2 (Invitrogen), 50 mM MgCl2 (Invitrogen), 10mM dNTPs, forward and reverse primers (see **Table S7** for primer sequences), and nuclease-free water. The MasterCycler EPGradient (Eppendorf) was used to amplify each mixture for 35 PCR cycles at an annealing temperature of 55℃. Amplified DNA was mixed with 6X loading dye (ThermoFisher Scientific), and 10μL of this product was loaded into each lane of a 1.5% agarose gel.

### Western blotting

#### Whole lysate Western blotting

Cells were collected from parallel primed (E8 medium) and TIRN (LIF-3i/MEF) using Versene/Accutase (1:1 mixture) solution. Cells were washed in PBS pH 7.4 (Gibco, 10010-023) and pelleted. Cell pellets were lysed using 1x RIPA buffer (ThermoFisher Scientific, 89900), 1mM EDTA, 1x Protease Inhibitor (ThermoFisher Scientific, 78430), and quantified using the Pierce bicinchoninic acid (BCA) assay kit (ThermoFisher Scientific, 23225). For each sample, 25μg of protein was loaded on a NuPage Bis-Tris, 4-12%, 1.5mm precast gel (ThermoFisher Scientific) for electrophoretic separation according to manufacturer’s protocols. The gel was transferred to a PVDF membrane using the iBlot2/iBlot3 systems (ThermoFisher Scientific). Samples were blocked in Tris-buffered saline (TBS), 5% non-fat dry milk (Research Products International), 0.1% Tween-20 (TBS-T) for 1 hour and incubated overnight at 4°C with primary antibody using gentle agitation. Membranes were rinsed 3 times in TBS-T, incubated for 1 hour at room temperature with either horseradish peroxidase (HRP) –linked goat anti-rabbit secondary antibody (Cell Signaling, 7074) or HRP –linked horse anti-mouse secondary antibody (Cell Signaling, 7076) for rabbit and mouse sourced antibodies, respectively, then rinsed 3 times. The membranes were developed using the Pierce ECL Western Blotting Substrate (ThermoFisher Scientific, 32209). Chemiluminescence detection was imaged using an Amersham Imager 600 (Amersham). Equal protein loading was verified for each membrane using a control antibody (e.g., anti-Actin).

#### Subcellular fraction Western blotting

Parallel isogenic hPSC were expanded in primed (E8) and TIRN (LIF-3i) culture systems and isolated using Enzyme-Free Cell Dissociation Buffer (ThermoFisher Scientific). After one wash in PBS, cell pellets were snap frozen in liquid nitrogen and stored at -80°C until analysis. Whole cell lysates were prepared using RIPA buffer (ThermoFisher Scientific, 89900), 1mM EDTA (Sigma), 1x Protease Inhibitor (ThermoFisher Scientific, 78430) for 15 minutes on ice. For histones, PRC1 and PRC2 proteins, lysis buffer was supplemented with 10U of benzonase nuclease (Sigma) and 4°C, 30min gentle continuous agitation. Protein fractions were isolated using the Subcellular Protein Fractionation Kit for Cultured Cells (ThermoFisher Scientific, 78840) according to the manufacturer’s protocol. The protein content was quantified using the Pierce BCA assay (ThermoFisher Scientific) method. For each assay, 25µg of total protein were loaded into a pre-cast NuPAGE 4-12% Bis-Tris gel (ThermoFisher Scientific, NP0336). Protein separation was performed by electrophoresis (100V, 2 hours) using the Novex Mini Cell (ThermoFisher Scientific) electrophoresis system. Proteins were transferred onto nitrocellulose membranes using iBlot2 NC Mini stacks and the iBlot2 blotting system (ThermoFisher Scientific). Alternatively, proteins >150kDa (i.e., DNMT1) were slowly (60V, 2 hours) transferred at 4°C, onto PVDF membranes using the Mini PROTEAN Tetra Cell system (Bio Rad). Membranes were first incubated for 1 hour at ambient temperature in blocking solution consisting of Tris Buffer Saline (TBS, Boston BioProducts), 0.1% Tween-20 (TBST), 5% nonfat dry milk to block unspecific antibody binding. Membranes were washed 3 times in TBST and then incubated overnight at 4°C with primary antibodies that were diluted in blocking buffer (mouse primary antibodies) or in TBST, 5% BSA (rabbit primary antibodies) at indicated dilutions (**Table S7**). Membranes were washed 3 times with TBST and incubated for 1 hour at room temperature with horseradish peroxidase (HRP)-conjugated secondary anti-mouse (Cell signaling, 7076) or anti-rabbit (Cell Signaling, 7074) antibodies (1:1000 in blocking buffer). After 3 washes in TBST, samples were exposed to Pierce ECL Substrate (Thermo Fisher, 32106) or Amersham ECL Substrate (GE, RPN2235) for chemiluminescent detection of proteins. For all Western blotting experiments, detection of actin and/or HSP90 was performed to control for equal loading of proteins. Membrane exposure and chemiluminescence detection were achieved using an Amersham Imager 600 (Amersham).

#### AF1521 PAR resin pulldown Western blotting

Cells were collected and pelleted as previously described. For AF1521 pulldown, cell pellets were lysed in a buffer containing 50mM Tris, pH 8, 200mM NaCl, 1mM EDTA, 1% Triton X-100, 10% glycerol, 1 mM DTT, 0.5% deoxycholate, and 1x protease inhibitor (ThermoFisher Scientific, 78430) and quantified using the Pierce bicinchoninic acid (BCA) assay kit (ThermoFisher Scientific, 23225). For each sample (1mg of protein), the ADP ribosylated fraction was separated the Af1521 Macrodomain Affinity resin set (Tulip Biolabs, 4301) following the manufacturer protocol. The Af1521 pulldown product (20μL per sample) was directly used for Western blot analysis as described above.

#### DNA damage response (DDR) and H_2_O_2_ oxidative stress assay Western blotting

To evaluate DNA damage, responses, isogenic primed and TIRN-SC were cultured in their respective medium supplemented with 100 ng/mL of the radiomimetic agent neocarzinostatin (NCS, Sigma N9162)) for 6 hours. Untreated hPSC were analyzed in parallel as controls. AF1521 pulldown and western blot analysis were performed as described above. To evaluate oxidative stress response, isogenic primed and TIRN-SC were cultured in their respective medium supplemented with 100μM hydrogen peroxide solution (Sigma, H1009) for 24 hours. Untreated hPSC were analyzed in parallel as controls. Western blot analysis was performed as described above.

#### Proteasome inhibitor MG-132 Western blotting

To evaluate proteasomal activity, isogenic primed and TIRN-SC were cultured in their respective medium supplemented with 62.5 nM proteasome inhibitor MG-132 (Selleckchem, S2619) for 24 hours, as previously described for hESC (Vilchez et al, 2012). Control hPSC were analyzed simultaneously as controls. Western blot analysis was performed as described above.

#### Mesodermal differentiation for PAR/PARP1 kinetics

Differentiation of primed vs TIRN-SC was performed using a modified protocol based on the STEMdiff APEL-Li medium system (Ng et al, 2008). APEL-2Li medium (StemCell Technologies, #5271) was supplemented with Activin A (25 ng/mL), VEGF (50 ng/mL), BMP4 (30 ng/mL), and CHIR99021 (1.5 μM) for the first 2 days, and then APEL-Li that was supplemented with VEGF (50 ng/mL) and SB431542 (10μM). Samples were collected using Accutase and immediately transferred in appropriate lysis buffer for Western blotting and Af1521 pull-down at 24hours, 48 hours and 96 hours.

### Trophectoderm stem cell differentiation

TSC were derived from TIRN-SC using a modified published protocol (Okae et al, 2017). TSC medium consisted of DMEM/F-12 supplemented with 2.5mM L-Glutamine, 15mM HEPES and 14mM sodium bicarbonate, 0.1mM β-mercaptoethanol, 0.2% embryonic stem-cell grade FBS (ThermoFisher Scientific), 0.3% bovine serum albumin (Sigma), 1% ITS-X supplement (ThermoFisher Scientific), L-ascorbic acid (1.5 μg/mL), EGF (50ng/mL), CHIR99021 (2 μM), A83-01 (0.5 μM), SB431542 (1 μM), valproic acid (0.8mM), Y27632 (2.5 μM), FGF10 (50 ng/mL), HGF (50ng/mL) and Noggin (20ng/mL). TSC medium was changed every 2-3 days and TSC were passaged about once a week using Trypsin-EDTA (Sigma). Samples were collected using Trypsin-EDTA and immediately transferred in lysis buffer (western blot) after 1 week (TSC passage 0; p0) and subsequent passages, or alternatively TSC were transferred for 1 passage onto 0.1% gelatin-coated 8-well Nunc Labtek II chamber slides (100,000 cells per slide) for immunostaining (fixation for 15 minutes using 2% formaldehyde in PBS).

### Injection of TIRN-SC into murine 8C-16C embryos and blastocyst embryos and processing of human-murine chimeric feti

#### Generation of human-murine fetal chimeras

Chimeric experiments employed either GFP+ RUES1, tdTomato+ RUES2 or tdTomato+mCherry+ RUES2 expressing constitutively murine E-Cadherin (see transfectant preparation section above). Prior to all interspecies chimera experiments, RUES1 and RUES2 cells were transitioned from E8 to LIF-3i for 4 to 7 passages. Puromycin was omitted the last passage before injection to ensure optimal viability of hPSC reporter clones. Expression of fluorescent proteins was verified and documented by photomicrographs using an epi-fluorescence microscope (Nikon Eclipse TE 2000-U). In addition, for some experiments, non-transduced RUES2 cells were prelabeled with the live stain Hoechst 33342 (1:2000 in TIRN medium, 10 minutes, Invitrogen H3570). TIRN-SC were gently dissociated using PBS-based enzyme-free cell dissociation buffer (ThermoFisher Scientific) and maintained in DMEM/F-12, 20% KOSR 0.1mM MEM non-essential amino acids, 1mM L-Glutamine, 0.1mM β-mercaptoethanol, 20ng/mL recombinant human LIF prior to injection. Uniform expression of fluorescent proteins was validated by flow cytometry analysis using a FACSCalibur flow cytometer (BD Biosciences). Embryo injections were performed by the Transgenic Core Laboratory at Johns Hopkins, Baltimore, MD. For most experiments, E2.5 (4 to 16 cells) or E3.5 C57BL/6J host embryos were collected by flushing uterine horns of super-ovulated females the day of injection. 5-7 stem cells were injected under the zona pellucida in the inner space of noncompacted 8-16-cell embryos or into the blastocoel cavity of blastocysts. Pseudopregnant ICR females were picked in estrus and mated with vasectomized males the day before (E0.5 pseudopregnant) embryo transfer. 5-7 stem cells were microinjected into host mouse embryos in a drop of KSOM under mineral oil. Injected embryos were maintained in KSOM prior to transfer into foster mothers. ∼10 embryos were transferred into the oviduct (E0.5 foster recipient) of pseudopregnant females. Fetuses were recovered at E9.5, E11.5, E12.5 or E14.5 from the uterine horns of the foster mothers. For each implantation site, fetuses were separated from the uterine wall using surgical forceps and iris scissors (Novo Surgical Inc.). Fetal and placenta/decidua tissues were gently dissected within a stem cell workstation and live fluorescence was documented for transgenic hPSC (Nikon Eclipse TE 2000-U).

#### Immunostaining of human-murine chimeric preimplantation embryos

Preimplantation murine embryos injected with human TIRN-SC were maintained in KSOM drops under oil and fixed using 1% formaldehyde in PBS for 10min at 4°C. For immunofluorescent staining, embryos were carefully manipulated using stripper tips (Origio) within a stem cell workstation cabinet under microscope. All staining procedures were similar to protocols detailed for Labtek chamber slides above, but with reduced volume. For each wash/incubation step, embryos were transferred between 50-100 μL drops under oil in a 30mm dish or directly onto 2-well Lab-Tek coverslips. After immunostaining, embryos were mounted with a coverslip using Prolong Glass Antifade Reagent (ThermoFisher Scientific) and imaged using confocal microscopy (LSM510 Meta, Carl Zeiss Inc., Thornwood, NY) at the Wilmer Eye Institute Imaging Core Facility, Baltimore, MD.

#### Whole-mount 3D-imaging of immunostained human-murine chimeric embryos

For whole-mount 3D imaging, isolated embryos were individually placed into 1.5mL Eppendorf tubes for fixation and staining. Embryos were rinsed in PBS, 5% goat serum, and fixed with freshly prepared PBS, 2% formaldehyde (1-2mL per embryo for 20 minutes in ice). Embryo preparation and clearing methods were slightly modified from a previously published protocol (Yokomito et al, 2012).

In brief, embryos were dehydrated on ice with 50% methanol/PBS for 10 minutes, followed by incubation in 100% methanol for 10 minutes. While embryos were in 100% methanol, they were transferred to chamber slides for imaging. The methanol was gently removed from the well without disturbing the embryo, and a 1:1 BABB (one part benzyl alcohol (Sigma, 402834) with two parts benzyl benzoate (Sigma, B6630)) / methanol mixture was added. This solution was gently discarded and the 1:1 BABB/methanol mixture was reapplied for two more washes before switching to 100% BABB solution. The BABB solution was also repeatedly replaced until the embryo was clear and ready for image analysis.

Human cells were detected directly with anti-human nuclear antigen (HNA) with or without anti-β-tubulin III antibody (clone TUJ1) co-staining to co-localize the neural system. Briefly, embryos were permeabilized with 0.1% Triton-X-100 in TBS solution (TBS-T), 5% goat serum for 1 hour on ice. All primary and secondary antibodies were diluted in blocking solution and incubated overnight at 4°C with at least 3 one-hour washes in TBST-T between each incubation with gentle agitation. After the final wash in TBS-T, embryos were incubated for 30 min with DAPI (1:500) and washed again in TBS-T.

#### Cryosections

For immunofluorescent staining of embryo/placenta cryosections, embryos or placentae were fixed in PBS, 2% formaldehyde for 15-35 min depending on the stage and size of embryos and washed in PBS. Embryos were incubated in 30% sucrose solution prior to cryopreservation in Tissue-Tek O.C.T compound solution (Fisher Scientific) after immersion in 2-Methylbutane (Sigma Aldrich) chilled with dried ice. Frozen embedded tissues were sectioned 8μm cryosections using cryostat at the Wilmer Eye Institute Imaging Core Facility. Tissue sections were stained using the protocol above with the antibodies mentioned in **Table S7**.

Embryos were imaged using confocal microscopy (LSM510 Meta, Carl Zeiss Inc., Thornwood, NY) at the Wilmer Eye Institute Imaging Core Facility, Baltimore, MD. (EY001765, Wilmer Core Grant for Vision Research, Microscopy Module).

#### Detection of human chimerism by genomic DNA PCR and RT-PCR

Feti and placentae were carefully dissected out from the uterine wall at E9.5, E12.5 or E14.5 and individually transferred into sterile Eppendorf tubes. Tissues were centrifuged at 500g, supernatant was discarded, and samples were snap-frozen in liquid nitrogen and stored at -80°C. DNA/RNA samples were isolated using the ZR-Duet DNA/RNA Miniprep kit (Zymo Research) according to manufacturer’s recommendation. Tissues were resuspended in DNA/RNA lysis buffer and homogenized using single-use RNase-free, DNase-free micro-pestle tips (Wilmad Lab Glass) and a Bel-Art micro-tube homogenizer. DNA and RNA samples were purified using Zymo-Spin columns, eluted in DNase/RNase-free water and quantified using a Nanodrop spectrophotometer (ThermoFisher Scientific).

For genomic DNA analysis, the detection of human DNA was performed using a sensitive mitochondrial DNA detection assay according to published methods^16^. For each sample and assay, 25ng DNA were amplified in triplicates in 384-well plates using the Power SYBR green master mix (ThermoFisher) and a ViAA7 Real Time PCR System (ThermoFisher Scientific) with a set of primers that amplify a human specific mitochondrial fragment. A second set of primers was used to amplify a specific ultra-conserved non-coding element (UNCE) as an invariant endogenous control to correct for sample-to-sample variations and errors. Relative quantification was determined by the Δ ΔCt method. Non-injected embryos were amplified in parallel and the frequency of human cells was estimated using genomic DNA standard curves created using human-mouse serial dilutions of hPSC and mouse cells genetically matched to the tested samples.

For detection of human RNA transcripts in human-murine chimeric feti, 2μg of total RNA that were isolated from all samples of cross-species fetuses and placentae were retro-transcribed using the superscript IV VILO master mix (ThermoFisher Scientific). For real-time PCR amplification, diluted (1:20) cDNA samples were admixed to the TaqMan Fast Advanced Master Mix (ThermoFisher Scientific) and Taqman gene expression assays (ThermoFisher Scientific). Each sample was simultaneously amplified in triplicates using human specific (GAPDH: Hs99999905_m1, ACTB: Hs99999903_m1) and human-mouse non-specific (GAPDH: Hs02758991_g1 ACTB Hs03023880_g1) primer-probe sets. All samples that displayed human genomic DNA above non-injected control embryos were cross-validated for mRNA analysis and the frequency of human cells was estimated using mRNA/cDNA standard curves that were created using human-mouse serial dilutions of hPSC and mouse cells genetically matched to the tested samples. Real time fluorescence detection was performed using a ViAA7 Real Time PCR System (ThermoFisher Scientific).

For all genomic DNA and mRNA experiments, each individual 384 well plate included a minimum of 3 stage-matched non-injected control mouse embryos and a 12-14 point serial dilution of human-mouse cells to create standard curves for each individual plate and eliminate experimental variations.

#### *In vitro* expansion of puromycin resistance transgene-expressing human hematopoietic and neural cells from human-murine chimeric feti

For *in vitro* expansion of puromycin-resistant hematopoietic lineages, E12.5 and E14.5 embryo fetal livers were separated, gently pipetted to disaggregate, and plated onto human fibronectin-coated plates (10 ug/mL/well) in endothelial growth medium 2 (EGM2, Lonza). After cultures were established in EGM2, the medium was switched to StemSpan SFEM (StemCell Technologies) supplemented with 50 ng/mL Flt3-ligand, 50 ng/mL TPO, 50ng/mL c-kit ligand (SCF), 20ng/mL IL2, 20 ng/mL IL6, 50 ng/mL GCSF, 3 units/mL EPO and 0.5 ug/mL puromycin. Medium was replaced every 2-3 days and analyzed 6 weeks after *ex-vivo* culture by flow cytometry (GFP, human CD34, hHLA) and immunofluorescence (HNA) according to the protocols described above.

For *in vitro* expansion of puromycin-resistant neural lineages, heads of chimeric E12.5 embryos were minced, digested in 0.05% Trypsin and plated on Matrigel-coated plated with PSC neural induction medium (NIM, ThermoFisher Scientific, A1647801). After adherent cells reached 60-70% confluency, NIM was supplemented with 0.5 ug/mL puromycin. Cells were passaged every 7-10 days with continuous puromycin throughout 6-week *ex-vivo* expansion. Puromycin concentration was increased to 1 μg/mL once the explants were established to further select human cells. Occasionally, retinal pigmented epithelial (RPE) cells were observed within neural cultures (not shown). Human cell identity was validated by immunofluorescence to detect HNA and NFM according to the protocol described above.

#### Flow cytometric analysis of human-murine chimeric fetal cells

Single cell preparations of E7.5 embryos and hematopoietic explants for direct flow cytometric analysis of GFP expression were prepared using either 0.05% trypsin-EDTA or Accutase for 5 min at 37°C.

### Bioinformatics

The NIH and ArrayExpress accession numbers of the data that was analyzed in this manuscript include: GSE141639 (bulk RNA-seq of primed and TIRN-SC ^3^, E-MTAB-3929 (scRNA-Seq of human embryos; Petropoulos et al, 2016), GSE197265 (Ribo-seq and RNA-seq of human embryos; Zou et al, 2022), GSE74111 (RNA-seq of Parp1KO mESC (Liu and Kraus, 2017), GSE95515 (DUX4 ChIP-seq in hiPSC; Hendrickson et al, 2017). Annotations for transcription factors (Lambert et al, 2018), for PARP1 and TNKS substrates (Ayyappan et al, 2021, Li et al, 2017, Nie et al, 2020, Bhardwaj et al, 2017) were compiled from published datasets.

#### Bulk RNA-Sequencing studies

Isogenic hPSC samples were prepared as described above, washed in PBS and cell pellets (1-2 million cells) were snap-frozen. The mRNA isolation and strand specific mRNA libraries were performed by the Genetic Resources Core Facility, Johns Hopkins Department of Genetic Medicine, Baltimore, MD. Briefly, mRNA poly-A selection and RNA library were prepared using NEBNext Poly(A) mRNA Magnetic Isolation Module (NEB#E7490) and NEBNext Ultra II Directional RNA Library Prep Kit for Illumina (NEB#E7760). Stranded mRNA libraries were sequenced on an Illumina Novaseq 6000 instrument using 50bp paired-end dual indexed reads. Raw data in Figure 1 was published (Park et al, 2022) with NIH Gene Expression Omnibus accession number GSE141639 and was realigned to hg38. Reads were aligned to GRCH38 using STAR (version 2.7.10a) Python software (Dobin et al, 2013). An index for the hg38 genome was prepared using the Homo_sapiens.GRCh38.dna.primary_assembly.fa fastq file and the human hg38 Ensembl v109 annotation gtf file (Homo_sapiens.GRCh38.109.gtf). Read alignment was executed using STAR with the following options: –outSAMtype BAM Unsorted SortedByCoordinate –quantMode TranscriptomeSAM GeneCounts. Summarized experiment objects were created using the R packages ‘Rsamtools’, ‘GenomicFeatures’ and ‘GenomicAlignments’: summarizeOverlaps (features=exonsByGene, reads=bamfiles, mode=“Union”, singleEnd=FALSE, ignore.strand=FALSE, fragments=TRUE). Differential expression analysis was executed using the R package DESeq2 (Love et al, 2014). Ensembl IDs were converted to gene symbols using the AnnotationHub package and current Ensb.Hsapiens objects (e.g., v110). Transcription factors annotations were obtained from a published database (Lambert et al, 2018). We performed k-mean clustering of RNA-seq data of human preimplantation embryos using average FPKM counts^1^ and the ‘pheatmap’ R package (kmeans_k =6). We defined a preZGA gene cluster as the combined gene selection identifying 1C, 2C and 4C samples, as well as 8C, ICM and hESC clusters specific to the corresponding cell-staged embryos. Volcano plots were created using the R package ‘enhanced Volcano’. Scatter plots and MA plots were generated using ‘ggplot2’. Euler diagrams were generated using the ‘eulerr’ R package. For heatmaps, a matrix of mean-subtracted regularized log counts was generated using ‘DESeq2’. Batch effects were removed using the package ‘limma’ removeBatchEffect command and heatmaps were constructed using the ‘Complex Heatmap’ R package. Differential expression of transposable elements was executed using the TEtranscripts Python package (Jin et al, 2015) using --sortByPos --format BAM --mode multi and the GRCh38_rmsk_TE.gtf and hg38_rmsk_TE.gtf gtf files.

<u>Parp1 KO mESC RNA-Seq data was obtained from</u> NIH Gene Expression Omnibus accession number <u>GSE74111 (Liu and Kraus, 2017).</u> Reads were aligned to GRCm39 using the STAR software similarly to the human data. An index was prepared using the Mus_musculus.GRCm39.dna.primary_assembly.fa fastq file and the mouse GRCm39 Ensembl v111 annotation gtf file (Mus_musculus.GRCm39.111.gI). A count matrix was directly constructed using STAR read counts. A matrix of mean-subtracted regularized log counts was generated using ‘DESeq2’. Orthologous human genes were annotated using BioMart.

#### Single cell RNA sequencing

For single cell RNA-seq studies, 10X Chromium (10X Genomics) barcoding, library construction and sequencing (Single Cell 3’ v3) were performed by the Genetic Resources Core Facility, The Johns Hopkins Department of Genetic Medicine, Baltimore, MD. Concurrent RUES02 cultures in E8 (primed) and LIF-3i (TIRN; TIRN +iDUX4 after 24 hours supplementation with 2μg/mL doxycycline) were gently collected using a mixture 1:1 Versene/Accutase and passed through a 70μm cell strainer. Viable cells were enriched using the MACS Dead Cell Removal Kit (Miltenyi Biotec) and prepared in PBS, 0.4% BSA. Cell quality check (cell viability >95%, cell diameter) was verified before preparing libraries. Library preparation was performed according to the manufacturer’s instructions (Chromium Next GEM Single Cell 3’ Gene expression). A low sequencing depth MiSeq run was used to assess quality control (Q30 Bases >96%). 10X libraries were sequenced on an Illumina Novaseq 6000 instrument (28×8×91 run configuration).

A custom hg38 reference genome was built using the Cell Ranger (10X Genomics) Linux software to align sequenced reads to a more inclusive up-to-date Ensembl Id library (v109), including protein-coding genes of interest that are currently annotated as non-coding RNAs (e.g., *TPRXL*) with the code: cellranger mkgtf Homo_sapiens.GRCh38.109.gtf Homo_sapiens.GRCh38.109.filtered.gtf --attribute=gene_biotype:protein_coding --attribute=gene_biotype:lincRNA --attribute=gene_biotype:antisense --attribute=gene_biotype:processed_transcript --attribute=gene_biotype:transcribed_processed_pseudogene. The reference genome was built with cellranger mkref --genome=GRCh38 --fasta=Homo_sapiens.GRCh38.dna.primary_assembly.fa --genes=Homo_sapiens.GRCh38.109.filtered.gtf --memgb=32 --nthreads=20. FASTQ files were aligned to this custom hg38 reference using the 10X Genomics Cloud Analysis platform (Cell Ranger Count v7.0.1 pipeline). Raw data and demultiplexed aligned non-normalized Cell Ranger output files are provided under GEO accession number TBA. The human embryo data^17^ was obtained via the SRA toolkit prefetch program (Accession # PRJEB11202) and realigned to hg38 using the STARsolo software (Kaminow et al, 2021). As mentioned above for bulk RNAseq, the index was prepared using the Homo_sapiens.GRCh38.dna.primary_assembly.fa fastq file and the Homo_sapiens.GRCh38.109.gtf gtf file for Ensemblv109 annotations. STAR parameters were: --runDirPerm All_RWX --readFilesCommand zcat $SORTEDBAM --soloType SmartSeq --readFilesManifest ../$TAG.manifest.tsv --soloUMIdedup Exact --soloStrand Unstranded --soloFeatures Gene GeneFull --soloOutFileNames output/ features.tsv barcodes.tsv matrix.mtx. The human embryo metadata is available at E-MTAB-3929.

Downstream analysis was performed using the Seurat (v4.4) R package (Satija et al, 2015). Seurat objects were created for each sample using Read10X (or ReadSTARsolo for the embryo data) and Create SeuratObject commands. First, Seurat objects were prepared individually by subsetting low quality cells, cell multiplets and dying cells by filtering using ‘nFeatureRNA’ and the percentage of mitochondrial genes (“percent.mt” feature as defined by PercentageFeatureSet(object, pattern = “^MT-“). After log normalizing and scaling the data (‘NormalizeData’ and “ScaleData’ functions), we performed linear dimensional reduction (‘RunPCA’ command) and graph-based clustering (Louvain algorithm) using the ‘FindNeighbors’ (dims = 1:50) and ‘FindClusters’ (resolution = 0.5) functions. The datasets were visualized by running the non-linear dimensional reduction Uniform Manifold Approximation and Projection (UMAP) using the ‘RunUMAP’ command (dims = 1:50). The “nFeature_RNA’ and “percent.mt” features were assessed within clusters using ‘FeaturePlot’ and any cluster that included residual low quality or dying cells were discarded from the analysis. The analysis was run again on the trimmed datasets (successively ‘NormalizeData (normalization.method = “LogNormalize”, scale.factor = 10000)’, ‘FindVariableFeatures (selection.method = “vst”, nfeatures = 2000)’, ‘ScaleData’, ‘RunPCA’, ‘FindNeighbors (dims = 1:50)’, ‘FindCLusters (resolution = 0.5)’, ‘runUMAP’) to verify data consistency. To account for sequencing depth variations between all 3 samples, we normalized all 3 datasets using the ‘SCTransform’ function (vst.flavor = “v2”, return.only.var.genes = FALSE). The normalized data were merged. ‘VariableFeatures’ for the SCT slot were defined using the VariableFeatures on the SCTransformed objects without the “return.only.var.genes = FALSE” command). Follow up analysis included successively ‘RunPCA’, ‘FindNeighbors (dims = 1:50)’, ‘FindClusters (resolution = 0.5)’, ‘runUMAP’. 3D UMAP data were constructed using Seurat “RunUMAP (dims = 1:50, n.components = 3L), ‘Embeddings’ and ‘Fetchdata’ functions and imported into the plotly 4.10 R package. 2D UMAP were graphed using Seurat ‘DimPlot’ and ‘FeaturePlot’ functions. For violin plots, the data was extracted from the Seurat objects using ‘FetchData’ and plotted using the ‘ggplot2’ package. The human embryo data was processed similarly. Trophectoderm and endodermal cells were excluded by selecting the identifiers “not applicable” and “epiblast” in the “Characteristics.inferred.lineage” metadata from the authors. Gene sets distinguishing ‘E3’, ‘E4-E5’ and ‘E5-E6 epiblast’ were obtained using the ‘FindMarkers’ function (avg_log2FC >log2(2), p_val_adj <0.05). Gene sets were curated using top differentially expressed genes in bulk RNAseq analysis of TIRN/TIRN DUX4 cells. The average expression of modules of interest was calculated by the function ‘AddModuleScore’. The code used to prepare each Figure can be provided upon request and will be uploaded into Github following publication.

#### Whole proteome analysis

Proteomic studies were performed by the Translational Proteomics Laboratory at the Johns Hopkins University, Baltimore, MD. A first set samples included parallel RUES02 hESC cultures in primed E8 (n=3), TIRN LIF-3i (n=3) and TIRN + iDUX4 after 24 hours supplementation with doxycycline (n=3) conditions. A second set of samples included parallel RUES02 hESC cultures in primed E8 (n=3) and primed E8 + iDUX4 after 24 hours supplementation with doxycycline (n=3) conditions. Freshly prepared samples were homogenized in lysis buffer. Pooled samples served as quality control for each experiment. Samples were labeled with 10-plex TMT with >98% efficiency. Mass spectrometry analysis was performed using the Orbital Fusion Lumos instrument (Thermo Fisher Scientific) and either easy-nano-LC 1200 or ultimate 3000 nano LC chromatography systems (Thermo Fisher Scientific). Database search was conducted using Proteome Discoverer and the UNIPROT genome UP000005640 (1% FDR). Volcano plots were created using the R package ‘enhanced Volcano’. Scatter plots and bar plots were generated using ‘ggplot2’ and Prism software. TNKS1/2 and PARP1 substrate annotations were summarized from published databases (Ayyappan et al, 2021, Li et al, 2017, Nie et al, 2020, Bhardwaj et al, 2017). Protein ubiquitination writers and erasers and histone ubiquitin readers, writers and erasers are curated from GO annotations and published protein sets (Vaughn et al, 2021).

#### Ubiquitinome studies

Ubiquitinome studies were performed by Creative Proteomics. These studies included parallel RUES02 hESC cultures in primed E8 (n=3), TIRN (n=3), primed E8 + iDUX4 after 24 hours supplementation with doxycycline (n=3) and TIRN DUX4 LIF-3i after 24 hours supplementation with doxycycline (n=3) conditions. Briefly, snap frozen samples were homogenized in lysis buffer (8 M urea, 1% protease inhibitor) by sonication. For each sample, 9.5mg of protein were digested by trypsin, peptides were purified using C18 reversed-phase columns, and ubiquitinated peptides were enriched using anti-K-ε-GG antibody beads. Mass spectrometry analysis was performed using the Ultimate 3000 nano LC chromatography system (Thermo Fisher Scientific). Data analysis was conducted using Maxquant and the UNIPROT genome UP000005640 (localization prob >0.75). Differentially ubiquitinated sites were identified with fold change > 1.5 and q value <0.05. Missing (Non-ubiquitinated) values in the primed controls were filled in using the sample minimum method (Kong et al, 2022).

#### ChIP sequencing (ChIP-Seq) studies

For ChIP-Seq, isogenic primed/TIRN hESC RUES02 samples (100×10^6^ cells per condition) were fixed using a freshly prepared solution of 11% formaldehyde, 5M NaCl, 0.5M EDTA pH8.0, 1M HEPES pH7.9 for 15min at room temperature with agitation. Fixation was stopped by adding 1/20 volume of 2.5M glycine solution. Henceforth, samples were kept on ice or refrigerated, washed with chilled 0.5% Igepal in PBS twice, with 1mM PMSF added to the second wash before snap-freezing pellets. Downstream sample preparation, sonication, immunoprecipitation, qPCR validation, library generation, QC and barcoding, and next-generation sequencing were executed by Active Motif. Duplicate samples for primed and TIRN-SC hESC were processed using the following antibodies: anti-BRD4 (Active Motif 91301), anti-H3K4me3 (39159), anti H3K27ac (Active Motif 39133), anti-H3K27me3 (39155), anti-PARP1 (Active Motif 39561), anti-NANOG (Active Motif 61419), anti-POU5F1 (Active Motif 39811), anti-SOX2 (Active Motif 39823) and anti-STAT3-phos (Cell Signaling 9145). Equal amounts of unprecipitated genomic DNA from all samples were pooled to generate input control libraries. 75-nt single-end sequence reads were generated using a NextSeq 500 sequencer (Illumina) and mapped to the human hg38 genome using VWA algorithm. BAM files were sorted using the samtools Python software ‘sort’ command and uniquely mapped reads were selected using samtools ‘view -b -F4’ and ‘view -b -q25’ commands. PCR duplicates were removed using samtools ‘markdup’. Initial peak calling was done using MACS2 software. Cut-off p-values were estimated by running a cut-off analysis using macs2 ‘callpeak --cutoff-analysis -f BAM -B --gsize=hs --tsize=75 --bw=200 -m 5 50 -n test_p’. Peak calling was run using MACS2 ‘callpeak’ with the following parameters: -f BAM -B --gsize=hs --tsize=75 --bw=200 -m 5.To handle replicates using the Irreproducible Discovery Rate (IDR) method (Huang et al, 2004), low stringent p-values were selected: H3K4me3 (p=0.001) NANOG (p=0.01), PARP1 (p=0.05), POU5F1 (p=0.01), SOX2 (p=0.001), phophoSTAT3 (p=0.01). The resulting narrowPeak files were sorted by -log10(p-value) using ‘sort - k8,8nr’ before running ‘idr –rank p.value’ to combine replicates. For protein/histone marks that bind to extended regions of the genome (i.e., BRD4, H3K27me3, H3K37ac), we utilized a p-value cut-off of 0.0001 with the following MACS2 parameters: ‘callpeak -f BAM -B --SPMR --broad --broad-cutoff 0.001 --gsize=hs --tsize=75 --bw=200 -m 5 50 ‘. Consensus broad peaks were defined using bedtools ‘intersect’. The human DUX4 ChIP data (Hendrickson et al, 2017) was obtained via the SRA toolkit prefetch program (Accession # PRJNA377315). Paired-end FASTQ files were aligned to the human hg38 genome using the bowtie2 aligner using the original publication settings: -t --sensitive-local -p 20 --no-mixed --no-discordant. SAM files were converted to BAM format and sorted using samtools ‘view’ and ‘sort’ commands. Sorted BAM files were filtered for uniquely mapped reads using samtools ‘view -b -q10’. Proper pairing statistics were verified using samtools ‘flagstat’. Peak calling was done using the MACS2 program with the following parameters: callpeak -f BAMPE -B --SPMR -q 0.05 --gsize=hs. Consensus DUX4 peaks were defined using IDR.

To compute quantitatively the differentially bound sites in NANOG, OCT4 and SOX2 ChIP-seq samples, we employed the ‘Diffbind’ Bioconductor R package. Corresponding IDR peaksets were used to create DBA objects (Diffbind::dba) for each individual factors or all three combined and provide a framework for the analysis. A binding matrix was calculated based on the read counts in the BAM files for each replicate using the function ‘dba.count’ and peaks were overridden to be re-centered at a uniformed length of 400bp. The data was normalized based on sequencing depth using ‘dba.normalize’. The primed samples were designated as reference samples with ‘dba.contrast(minMembers=2)’ before running the differential analysis using DESeq2 (‘dba.analyze’). Intervals within the hg38 human genome blacklist were removed and the ChIP input control bam file was used as control greylist reads for consensus peak selection. All bound sites were retrieved with ‘dba.report(method=DBA_DESEQ2, contrast =1 (for single factor analysis) or 4 (for SON co-binding), bCounts=TRUE, bCalled=TRUE, th=1)’. Peak annotations were added using the ChIPseeker R package and the ‘annotatePeak’ command using an Ensembldb object (v110) created with AnnotationHub. Peak overlaps with other peaks (e.g., H3K4me3, H3K27ac) were defined using the IRanges command ‘findOverlaps’. Promoters were defined as regions at +/- 3000bp from TSS. Enhancer regions were defined using H3K27ac consensus peaks. Euler diagrams were created using the R package eulerr and ChIPseeker::overlap as input. To produce heatmaps, a profile object was created in Diffbind using ‘dba.plotProfile’ that was imported into the profileplyr R package for additional annotation and customization using the command ‘generateEnrichedHeatmap’. Scatter plots and box plots were created from the dba.report output dataframe of dba objects using ggplot2. For DUX4 profiles, non-overlapipng Diffbind generated SON peaks. BAM files were indexed using samtools ‘index’ utility and the signal data was converted to bigwig using deeptools ‘bamCoverage’ using RPGC normalization: bamCoverage --binSize 10 --normalizeUsing RPGC --smoothLength 60 --extendReads 75 --centerReads --numberOfProcessors 20 --effectiveGenomeSize 2913022398. Input-substracted bigwig files were created using the deeptools “bigwigCompare” command and replicate bigwig files were averaged using the deeptools ‘bigwigAverage’ command. Input-subtracted averaged RPGC normalized profiles were visualized using the Integrated Genomic Viewer software. Line plots were created using deeptools ‘computeMatrix’ and “plotProfile” commands or alternatively, the deeptools matrix was imported in the profileplyr R package. Deeptools ‘computeMatrixOperations’ was used for relabeling samples and regions. Gene bodies were defined using the Ensembl Homo_sapiens.GRCh38.110.gene.bed bed file. The DUX4 matrix of scores (averaged RPGC) was created within Diffbind-calculated NSO gained peaks and the DUX4 data was graphed using Deeptools ‘plotProfile’.

Local motif enrichment analysis was conducted using CentriMo (Bailey and Machanick, 2012) within the MEME-ChIP suite v5.5.5. NSO and NANOG gained (FC2>2, FDR<0.05) sequences were uploaded as bed files using the UCSC human hg38 format and the “Vertebrates” and “Human and mouse (HOCOMOCO v12 CORE)” databases. Dot plots were created using ggplot2 using CentriMo E-values and log2 fold change from Deseq2 RNA-seq analysis.

### 3D rendering of ubiquitinated sites from Xray crystallography and AlphaFold predicted structures

The following PDB files were downloaded from the UNIPROT database: DNA-bound protein domains for NANOG (4RBO), SOX2 (1O4X), and PARP1 (4DQY) and were aligned to their full AlphaFold predicted structures, NAD+ (1A26) and ubiquitin (1D3Z) using the PyMOL function “cealign”

## SUPPLEMENTARY INFORMATION

### SUPPLEMENTARY TABLE

**Table S1. Differential gene and transposable element expression analysis of bulk RNAseq for eight independent isogenic TIRN-SC vs primed lines;** includes annotations for transcriptional regulators (Lambert et al, 2018) and DUX4-bound genes from hPSC DUX4 ChIP-Seq data (Hendrickson et al, 2017) ^2^.

**Table S2. Differential expression analysis of whole proteome of TIRN (T), TIRN+iDUX4 (TD) and primed hPSC+iDUX4 (PD) RUES02 cells vs isogenic RUES02 primed (P) hPSC control cells; i**ncludes cross-tabulation of RNA-protein values from concordant RNA-Seq data; with annotations for PARP1/TNKS substrates, transcriptional regulators ^1^ and early human embryo stage clustering (Zou et al, 2022 ^3^).

**Table S3. Differential gene and transposable element expression analysis of bulk RNAseq for TIRN (T) vs primed (P), TIRN+iDUX4 (TD) vs primed (P) and primed+iDUX4 (PD) vs primed (P) RUES02 hESC;** includes annotations for transcriptional regulators^1^, DUX4-bound genes (Hendrickson *et al* ^2^), early human embryo stage clustering (Zou *et al* ^3^), and concordance of overexpressed developmental genes in TIRN-SC with the same genes overexpressed in PARP -/- mESC (Liu and Kraus ^4^).

**Table S4. Single cell RNA-Seq annotated gene module sets used for Figure 3**.

**Table S5: TIRN-SC NSO ChIP-Seq and Motif data.** Includes ChIP-Seq identification of genomic locations and annotations for NANOG, SOX2, OCT4 (NSO) co-binding sites; as well as associated H3K27Ac, H3K4me3, H3K27me3, BRD4 marks for *all GAINED* co-occupied sites. Tables also include motif analysis of the most significant (FDR<0.05, fold change>2) NSO co-binding sites, NANOG-only sites, and SOX2-only sites for these significant TIRN GAINED sites. Tables also list associated genes with transcriptional start sites (TSS) within 10 kb of for ALL TIRN NSO GAINED sites, as well as the subset of GAINED NSO TIRN sites with the highest differentially bound significance (i.e., FDR<0.05, fold change (FC)>2).

**Table S6. Intensities and differential protein analysis of ubiquitinated peptide sites for TIRN, TIRN+iDUX4 and primed+iDUX4 stem cells vs primed control cells with cross-tabulation of ubiquitinome-proteome.** All tabs are annotated for known PARP1/TNKS substrates ^5^, transcriptional regulators ^1^, and early human embryo clustering ^3^.

**Table S7. Materials and Reagents.** Includes lists of all hESC and hiPSC lines, antibodies, inventory of bioinformatics samples, and PCR primers used in these studies.

### SUPPLEMENTARY MOVIE

**Movie S1. Confocal microscopic Z-stacks of representative human-mouse fetal chimeras after blastocyst injection of TIRN-SC.** Immunodetection of HNA^+^ human cells in an E9.5 human-murine fetal chimera with co-detection of neural β-Tubulin III (TUJ1) that resulted from injection of TIRN-SC RUES1 hESC into a murine blastocyst. Also includes control images from non-injected E9.5 control embryo to validate HNA/ β -Tubulin III (TUJ1) specificity.

**Movie S2. Confocal microscopy Z-stacks of representative human-mouse blastocyst chimeras after 8C mouse embryo injection of TIRN-SC.** (A) Phase contrast imaging and immunofluorescent staining of HNA^+^ human cells (red) at 4 and 24 hours (blastocyst stage) after injection of TIRN-SC RUES02 hESC into a murine 8C-16C embryo. Nuclei are stained with DAPI (blue). Immunostain for a control non-injected embryo is shown. (B,C) Phase contrast imaging and immunofluorescent staining of HNA^+^ human cells (red) at 48 hours (hatching blastocyst stage) after injection of TIRN-SC RUES02 hESC into murine 8C-16C embryos. Nuclei are stained with DAPI (blue) and trophectoderm cells are stained with CDX2 (green). For embryo #3 **(D)** GATA6^+^ cells (green) are labeled instead of CDX2.

**Movie S3. 3D modeling of differentially ubiquitinated sites in NANOG and SOX2 from X-Ray crystallographic and AlphaFold predicted structures**. Glycine-lysine conjugation sites are in red, predicted ubiquitin moieties are in purple.

### SUPPLEMENTARY FIGURES

**Figure S1.**
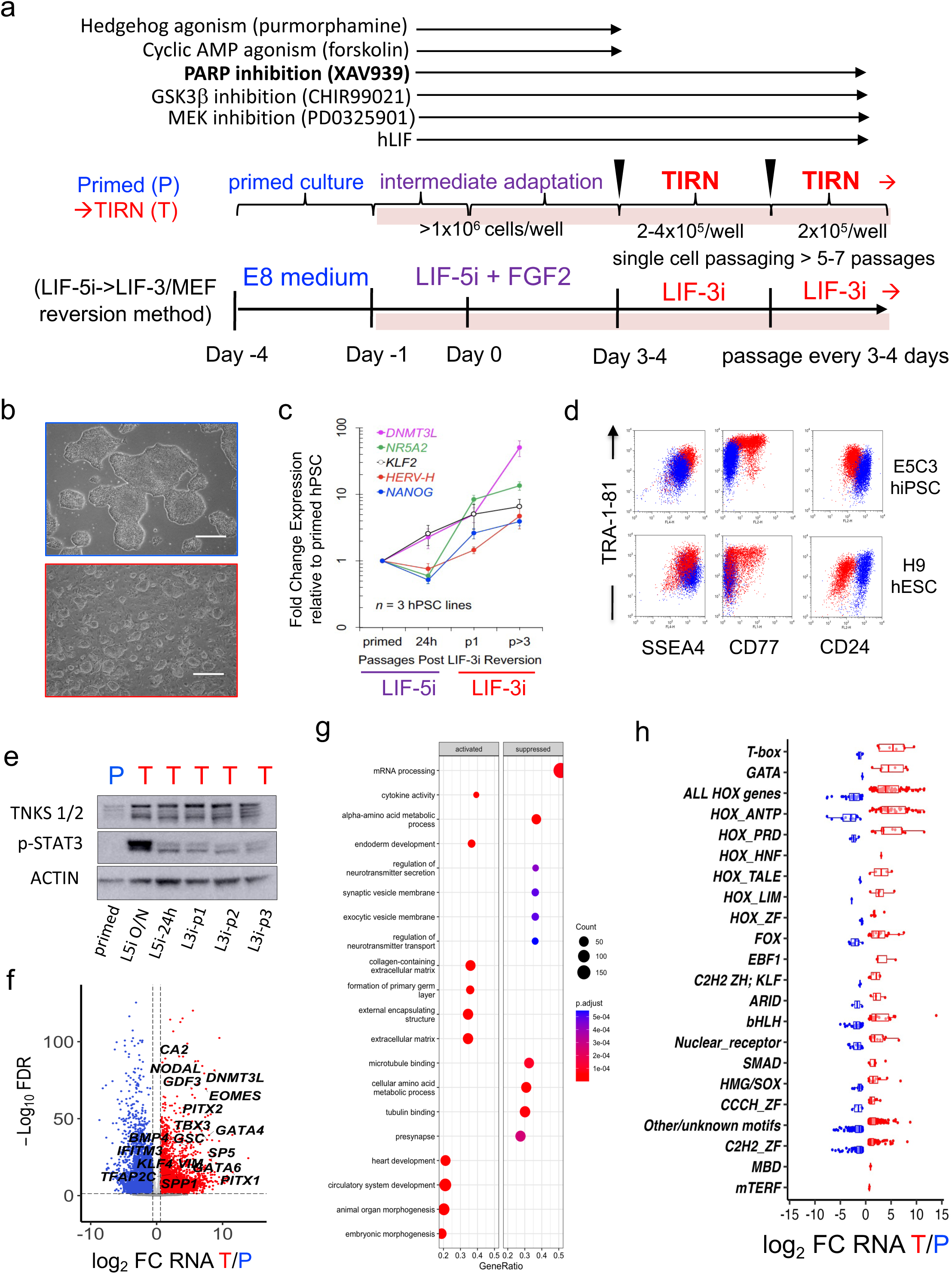
Chemical TIRN reprogramming of primed, conventional hPSC. (**a**) Summary schematic of the TIRN reversion method using LIF-5i->3i ^6-8^. (**b**) Photomicrographs of representative primed (top) vs TIRN (bottom, after 9 passages in LIF5i->3i conditions) hESC (RUES02). Scale bar = 500 μm. Following brief LIF-5i adaptation of primed hPSC and several passages in LIF-3i TIRN medium containing only XAV939, the GSKβ inhibitor CHIR99021, and the MEK inhibitor PD0325901, a broad genetic repertoire (**Table S7**) of independent conventional, bFGF-dependent, primed hESC and hPSC with a monolayer colony morphology become tolerant to bulk single-cell passaging and acquire a typical dome-shape morphology. (**c**) qRT-PCR analysis of kinetics of relative mRNA expression of naïve hPSC markers during early passages of LIF5i->LIF-3i TIRN reversion. Gene expression is normalized to ACTB and isogenic primed hPSC controls. Kinetic analysis during LIF5i->LIF-3i TIRN reversion revealed rapid upregulation of mRNA transcripts of human preimplantation naïve epiblast-specific genes (*e.g*., *DNMT3L, HERV-H, NR5A2, KLF2*, and *NANOG*) in TIRN-SC relative to their primed isogenic counterparts. Mean results are shown for three isogenic pairs of genetically independent primed and TIRN-reverted hPSC lines (*i.e*., H9 hESC; E5C3, 6.2 cord blood-hiPSC) following adaptation in LIF-5i, and 2-3 subsequent passages in LIF-3i. (**d**) Flow cytometric detection of changes in cell surface marker profiles of representative primed vs TIRN-SC derived from hESC (H9) and cord blood-hiPSC (E5C3). Profiles show merged flow cytometric expression level events of indicated cell surface proteins in isogenic primed (blue) and TIRN (red) hPSC. TIRN-SC expressed higher levels of ICM embryonic markers CD77, keratan sulphate-associated surface antigens (*i.e.,* TRA-1-60, TRA-1-81), and stage-specific embryonic antigens (*i.e*., SSEA3, SSEA4), but expressed lower levels of the primed, post-implantation epiblast marker CD24 ^9^. (**e**) Representative Western blot analysis of rapid upregulation of TNKS1/2 and phosphorylated STAT3 proteins during the first 3 passages of LIF-5i->LIF-3i TIRN reversion. ACTIN expression was used as control. (**f**) Volcano plot showing log2 fold change (FC) RNA expression between TIRN-SC (red) and primed (blue) hPSC (n=8 independent cell lines; mean data is shown). Colored dots show differentially expressed genes (FC>1.5, p< 0.05; delineated by dashed lines); non-significantly changed genes are greyed out. Highlighted are key pre-lineage blastomere and naïve ICM-specific genes expressed in both TIRN-SC and human preimplantation embryo cells (*e.g., DNMT3L, GSC, EOMES, GATA6, GATA4, NODAL, LEFTY2, GDF3, SP5, DPPA3 (STELLA), BMI1*, *CRIPTO, NR5A2, and TFAP2C* ^10^. Notably, NR5A2, TFAP2C, EOMES, GSC, and GATA6 are important regulators of lineage segregation to epiblast, primitive endoderm, and trophectoderm from pre-lineage blastomeres. (**g**) GSEA pathway analysis of differential protein expression between TIRN (red) and primed (blue) hPSC. (**h**) Bar plots of log2 FC RNA expression of pioneer factor gene families overexpressed in TIRN (T, red) vs primed (P, blue); see also **Table S1**.

**Figure S2.**
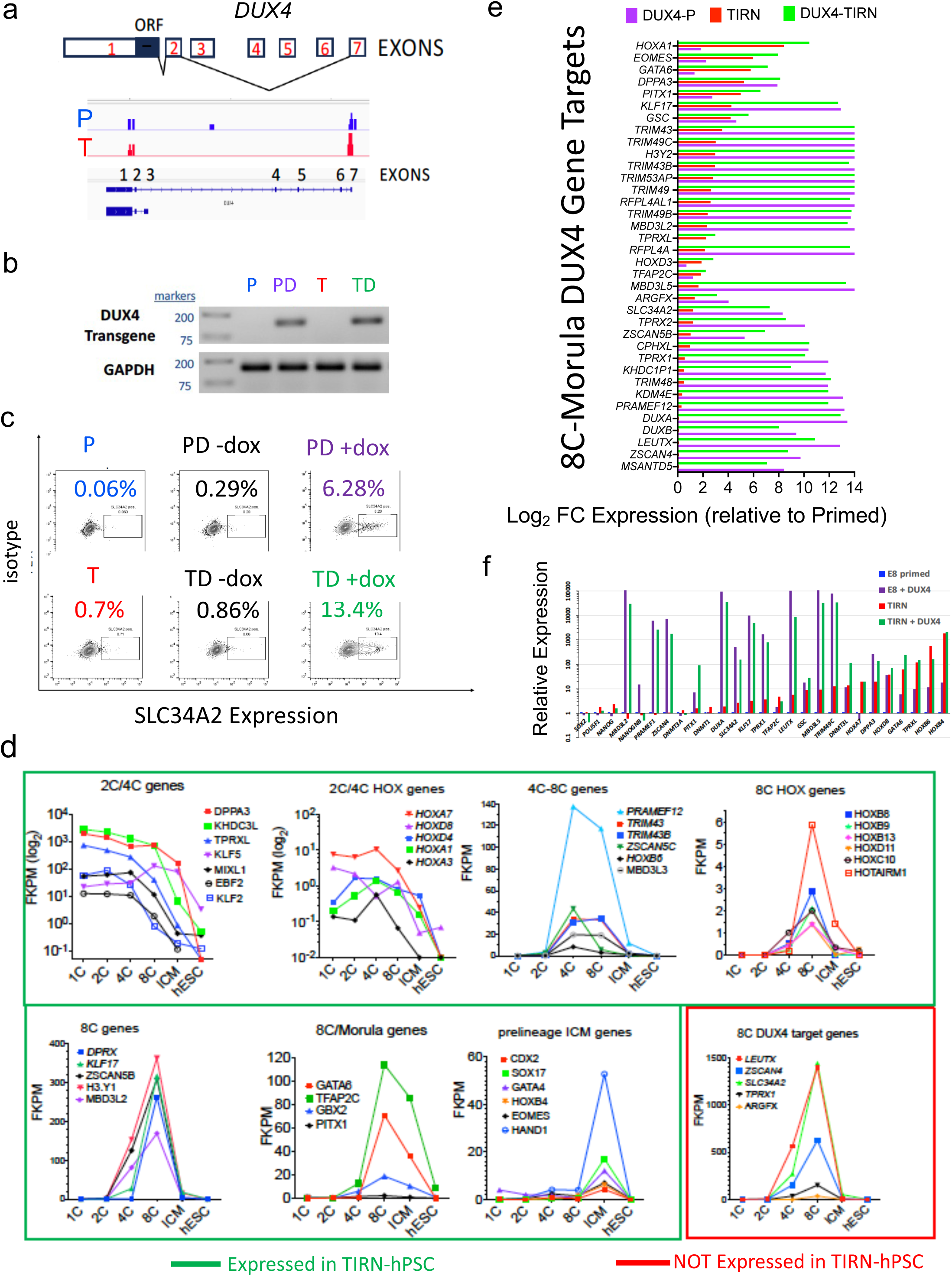
Generation of primed and TIRN-SC lines with a construct for the ectopic, inducible expression of the ZGA-priming pioneer factor DUX4. (**a**) **Endogenous *DUX4* expression in primed vs TIRN-SC.** RNA-seq profiles of the endogenous *DUX4* locus in primed (P, blue) vs. TIRN-SC (red) showing similar, low levels expression of germline DUX4 transcripts. Top schematic indicates that the detected germline DUX4 splice variants from RNA-Seq included exon 1 (the protein-coding ORF), exon 2, and exon 7. (**b**) **RT-PCR validation of transgenic (lentiviral) iDUX4 expression** following 24 hrs doxycycline inductions in isogenic primed (P), Primed +iDUX4 (PD), TIRN (T), and TIRN +iDUX4 (TD) cells by RT-PCR using transgene-specific primers. Detection of GAPDH RNA was used as control. (**c**) **Flow cytometry detection of surface expression of the 8C-specific NaPi2b transporter SLC34A2** ^11^ in primed+iDUX4 (purple) and TIRN+iDUX4 (green) after 24h doxycycline. TIRN-SC tolerated up to 24h of iDUX4 induction and resulted in over 2-fold higher fraction of SLC34A2 than isogenic primed cells. (**d**) **Expression of a large repertoire of 2C-4C, 8C, and pre-lineage morula/ICM genes in TIRN-SC.** Ribo-seq read data (log2 FPKM) from Zou et al ^3^ for 2C/4C/8C/pre-lineage expression kinetics during early human embryo development. Results show representative riboprobe 2C/4C/8C/morula factors that *were expressed* in TIRN-SC (green boxes), as well as representative, additional 8C-specific DUX4 targets (red box) that were expressed only after iDUX4-activation in TIRN-SC. (**e**) **Expression of DUX4 targets in TIRN and TIRN +iDUX4 cells.** Log2-transformed relative RNA-Seq expression of 8C-morula genes (from k-mean clustering of human embryo data ^3^) and DUX4-bound gene targets ^2^ in primed-iDUX4 (purple), Shown are ratios of TIRN (red) and TIRN+iDUX4 (green) cells / isogenic primed hPSC control expression. Although TIRN-SC already expressed a large repertoire of 8C DUX4 target genes, even in the absence of transgenic iDUX4, a smaller subset of DUX4 target genes (*e.g*. *LEUTX*, *TPRX1*) were activated robustly only after iDUX4. (**f**) **Real-time qRT-PCR validation** of RNA-Seq expression data in (**e**) of 8C DUX4 targets in primed-iDUX4 (purple), TIRN (red) and TIRN+iDUX4 (green) cells, relative to isogenic primed controls.

**Figure S3.**
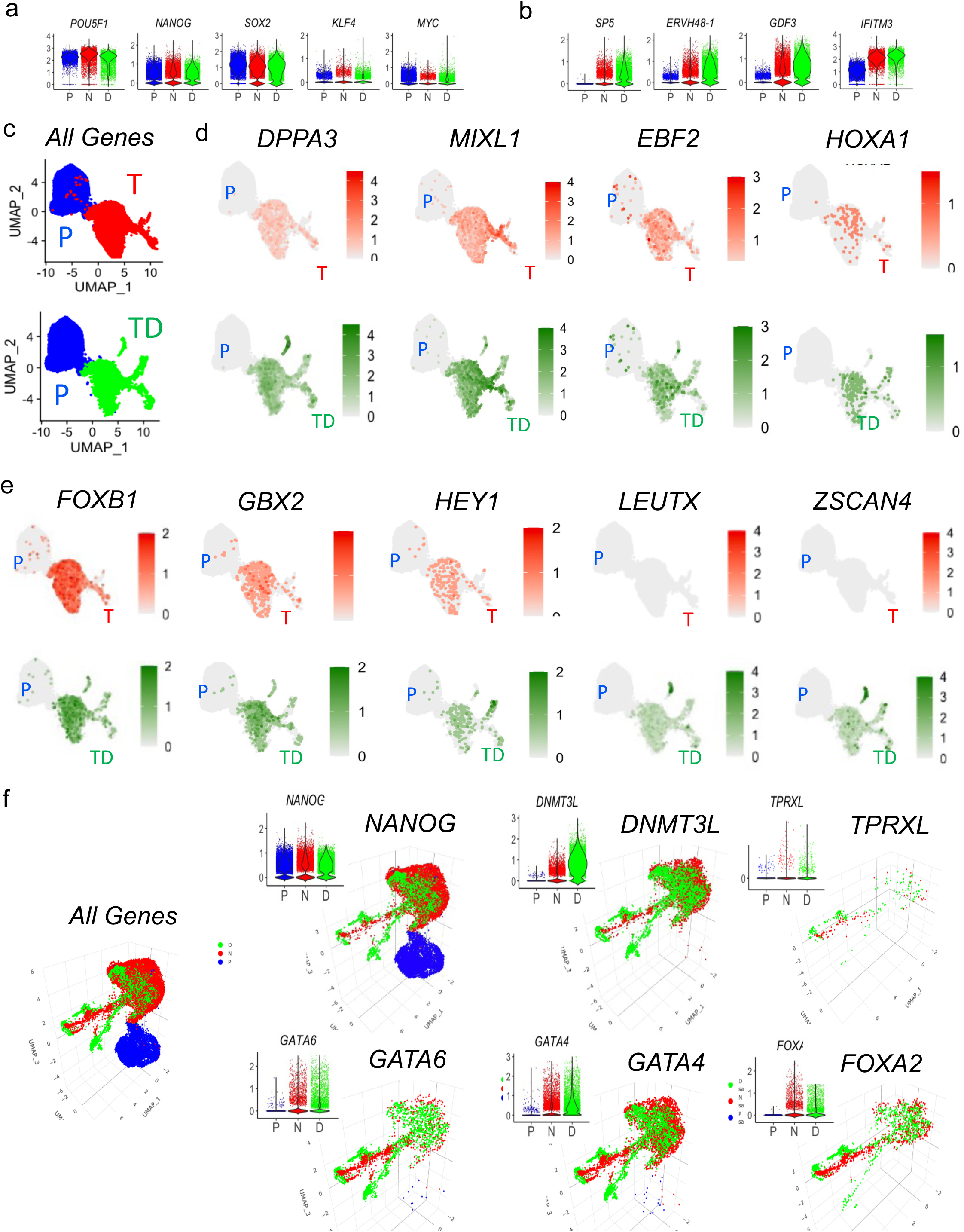
TIRN-SC were homogenous, single cell populations co-expressing 2C-4C, 8C, and pre-lineage morula genes: Single-cell RNA-Seq (scRNAseq) analysis of TIRN cells +/- iDUX4 activation. Violin plots show log-normalized corrected scRNAseq expression levels for (**a**) core pluripotency factor genes, and (**b**) naïve epiblast-specific genes. **(c, d, e**) 2D UMAP plots colored for (**c**) TIRN (red) vs primed (blue) and TIRN+iDUX4 (green) vs primed (blue) cells or showing normalized expression levels of select (**d**) 2C-4C-specific and (**e**) 8C-specific genes. (**f**) 3D UMAP plots showing cells from TIRN (red), TIRN+iDUX4 (green) and primed (blue) scRNA-seq analysis from embeddings exported using Seurat and visualized using plotly. 3D UMAP plots and accompanying Violin plots show single cell expressions and corresponding normalized expression levels for select 8C-to-morula-specific genes for either primed (P; blue) and TIRN (T; red) or TIRN+iDUX4 (TD; green) cells.

**Figure S4.**
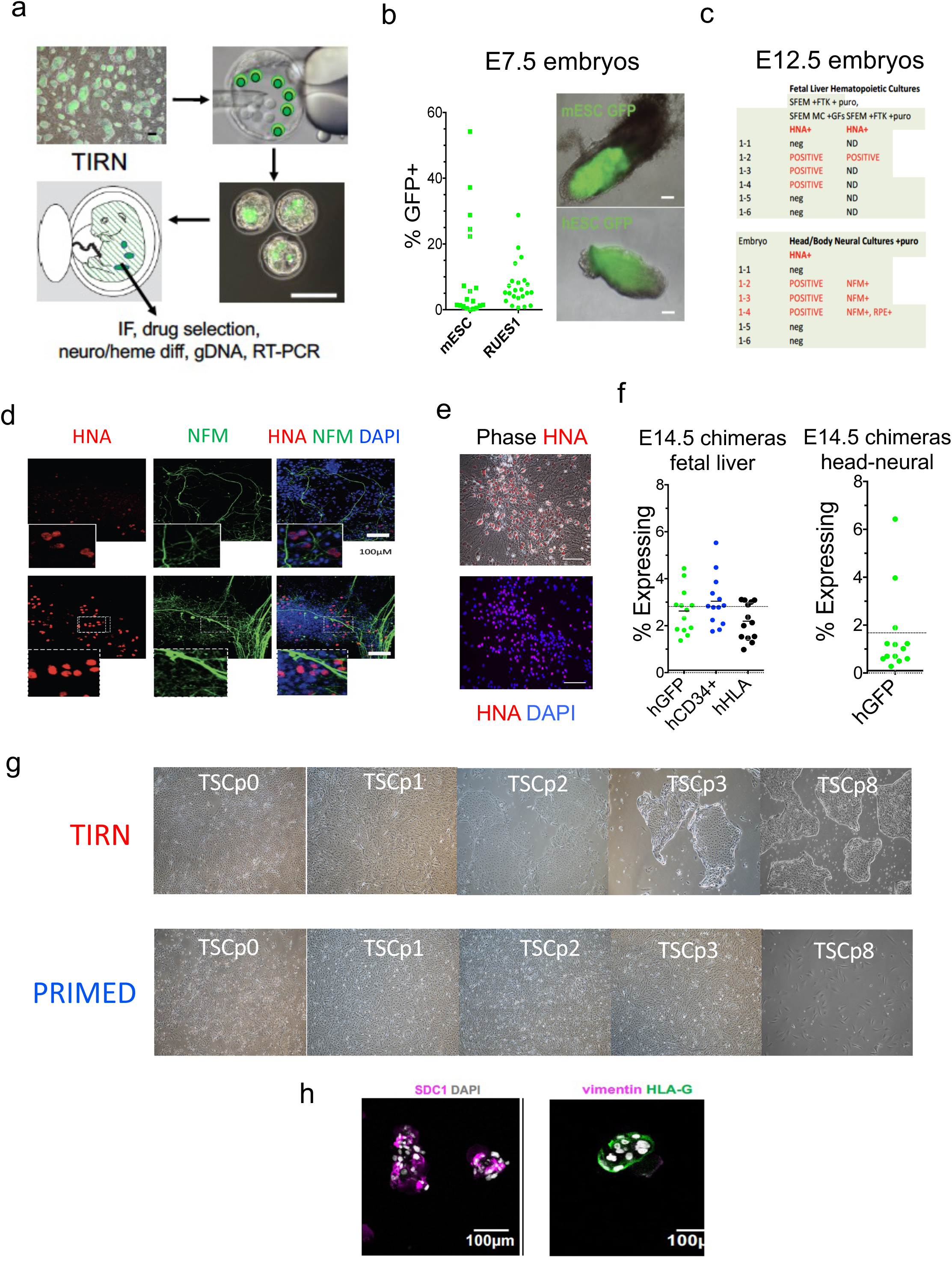
Multilineage contribution of human lineages in murine-human embryonic chimeras following injection of TIRN-SC into murine blastocysts. (**a**) Experimental design to assess and validate human-murine interspecies chimeras (**b**) Flow cytometry quantification of GFP^+^ cells in E7.5 human-murine chimeric embryos following injection of GFP^+^ human TIRN-SC or GFP^+^ control mESC. Merged phase/fluorescence microscopic images of representative E7.5 chimeras are shown. (**c**) Summary table of HNA^+^ characterization and *ex vivo* expansion of human cells from E12.5 chimeric embryo explants (fetal liver, head) following puromycin selection. (**d**) Representative immunofluorescent detection of HNA^+^(red) neurofilament (NFM^+^; green) human cell outgrowths from head explant of E12.5 human-mouse chimeric fetus. (**e**) Image of human HNA^+^ (red) neural outgrowths isolated by puromycin-resistance culture of E12.5 human-murine chimeric fetal head. (**f**) Summary of FACS analysis of GFP, human CD34 and human CD45 expression in *ex vivo* hematopoietic and neural cell culture explants isolated from human-murine fetal livers and heads, respectively, following puromycin cell culture. (**g**) Immunofluorescent staining of in vitro self-renewal of TIRN-TSCs from TIRN vs primed cells at each indicated passage, and (**h**) and *in vitro* differentiation toward SDC1+ syncytiotrophoblasts and HLA-G^+^ extravillous trophoblasts.

**Figure S5.**
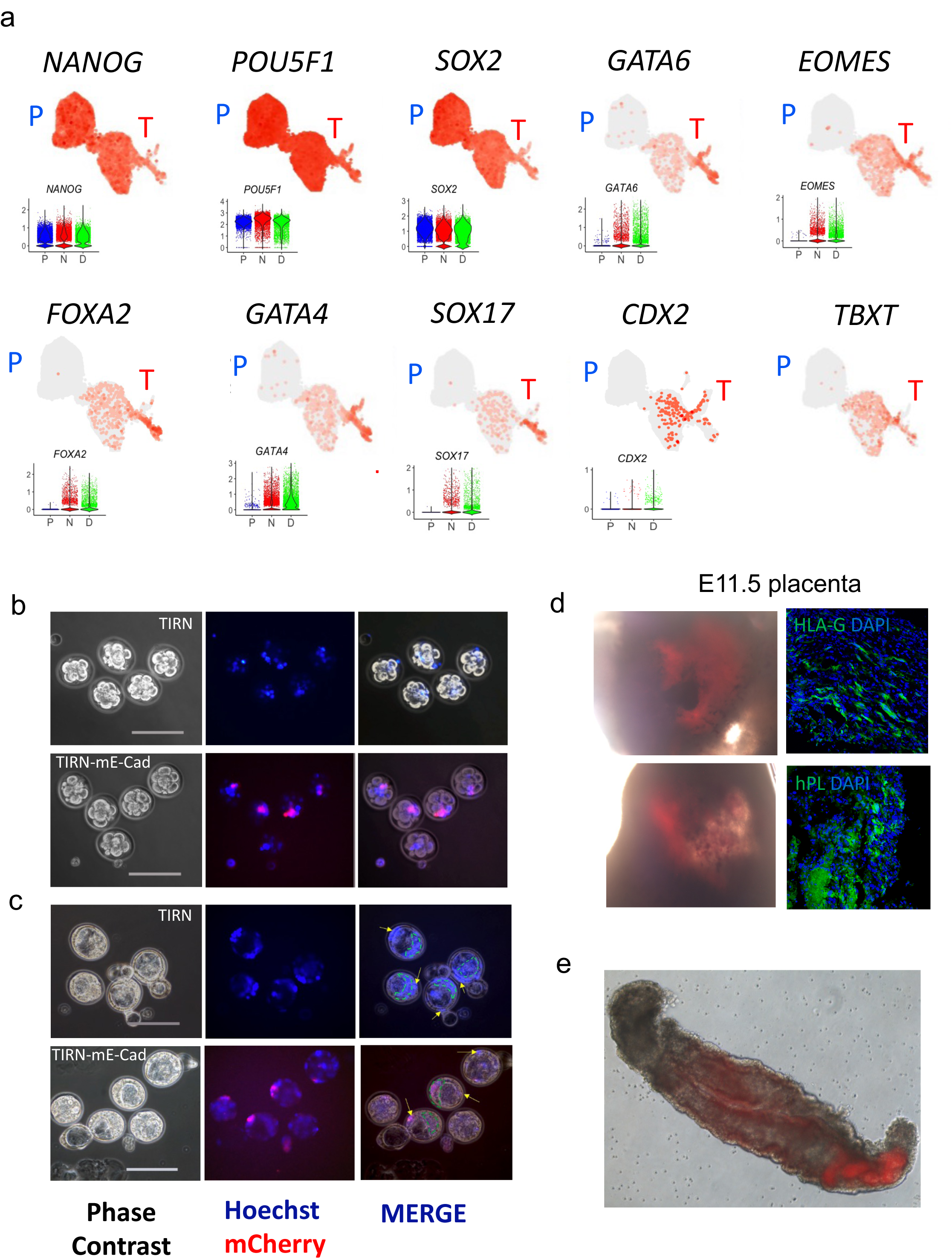
Direct contribution of blastomere-like TIRN-SC to both embryonic and extra-embryonic tissues in human-murine chimeric blastocysts and feti. **(a)** UMAP plots of log-normalized corrected counts showing *simultaneous, homogenous, single-cell expressions* for epiblast-specific (*NANOG, POU5F1(OCT4), SOX2*), primitive endoderm-specific (*GATA6*, *FOXA2, GATA4, SOX17*) and trophectoderm-specific (*GATA3, CDX2, EOMES*) genes in primed (P) vs TIRN (T) cells. (**b,c**) Merged live detection of Hoechst-labeled TIRN +/- mECad cells at 4 (**b**) and 24 (**c**) hours following injection of 5-7 TIRN-SC into 8C-16C murine embryos demonstrating rapid multi-lineage TE and PE compartment integration at 5-7 sites of murine blastocysts; in an equipotent manner. (**d**) Fluorescent detection of tdTomato+ cells (red) in the placenta of E11.5 human-murine chimeric fetus with immunofluorescent detection (green) of human specific HLA-G and human placenta lactogen (hPL); following injection of TIRN-SC into 8C-16C murine embryos. (**e**) Representative example of an abnormally developed E11.5 human-murine chimeric embryos following injection of tdTomato^+^ TIRN-SC (red) into 8C-16C murine embryos.

**Figure S6.**
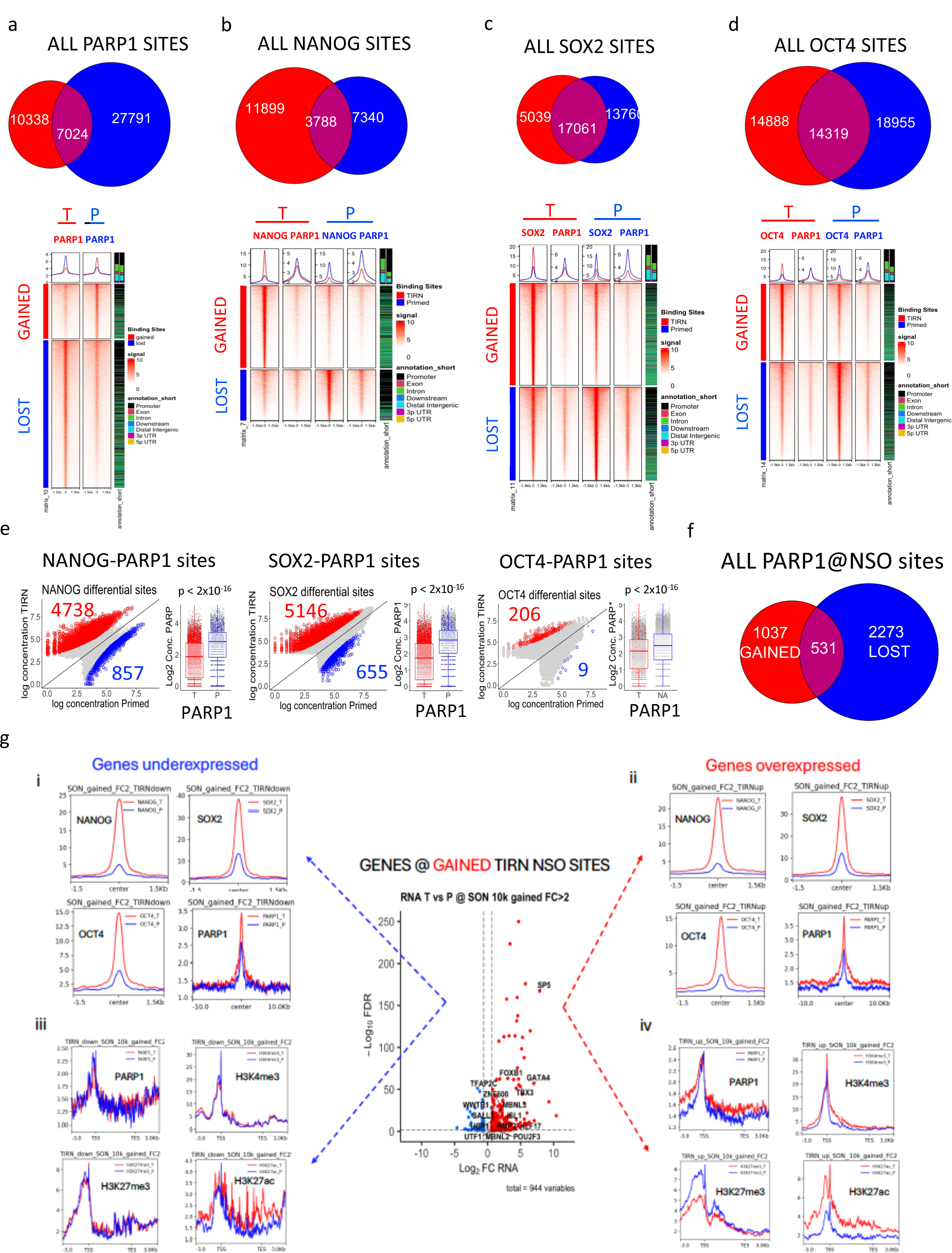
ChIP-Seq analysis of TIRN-SC chromatin. (**a, b,c, d**) Euler diagrams showing overlap of PARP1, NANOG, SOX2, and OCT4 binding sites (400 bp regions) in isogenic TIRN (red) vs primed (blue) hESC RUES2. All 400 bp sites were normalized and input-corrected using Diffbind software. Heatmaps show normalized read signals centered at non-overlapping PARP1, NANOG, SOX2 or OCT4 bound peaks with top line plots summarizing average signal for each ChIP-Seq experiments (average of 2 replicates) at TIRN (red line) vs primed (blue line) sites. Genomic distribution of peaks was annotated in the heatmaps using profileplyr. (**e**) Scatter plots for averaged (n=2) log-transformed concentrations of NANOG, SOX2 and OCT4 at NANOG, SOX2 and OCT4 sites. Differentially bound sites are indicated for primed (blue) and TIRN (red) samples. Adjacent box plots show the PARP1 concentrations at NANOG, SOX2 and OCT4 sites in TIRN vs. primed cells (differential sites are colored). (**f**) Euler diagram showing the number of overlapping PARP1, NANOG, SOX2, and OCT4 co-binding sites in isogenic TIRN (red) vs primed (blue). (**g**) Volcano plot of RNA-seq differential expression analysis for genes significantly bound by NSO in TIRN-SC (gained sites, FC>2, FDR<0.05) within 10kb of TSS. Line plots of averaged ChIP-seq signal for NANOG, SOX2, OCT4 and PARP1 centered on NSO sites in TIRN (red) and primed (blue) cells are shown for significantly (i) downregulated (FC<-1.5, adj. p value<0.05) and (ii) upregulated (FC>1.5, adj. P value<0.05) genes. Line plots of averaged ChIP-seq signal for PARP1, H3K4me3, H3K27me3 and H3K27me3 at gene bodies that are significantly bound (FC>2, FDR<0.05) NSO sites in TIRN (red) and primed (blue) cells are shown for significantly (iii) downregulated (FC<-1.5, adj. p value<0.05) and (iv) upregulated (FC>1.5, adj. p value<0.05) genes.

**Figure S7.**
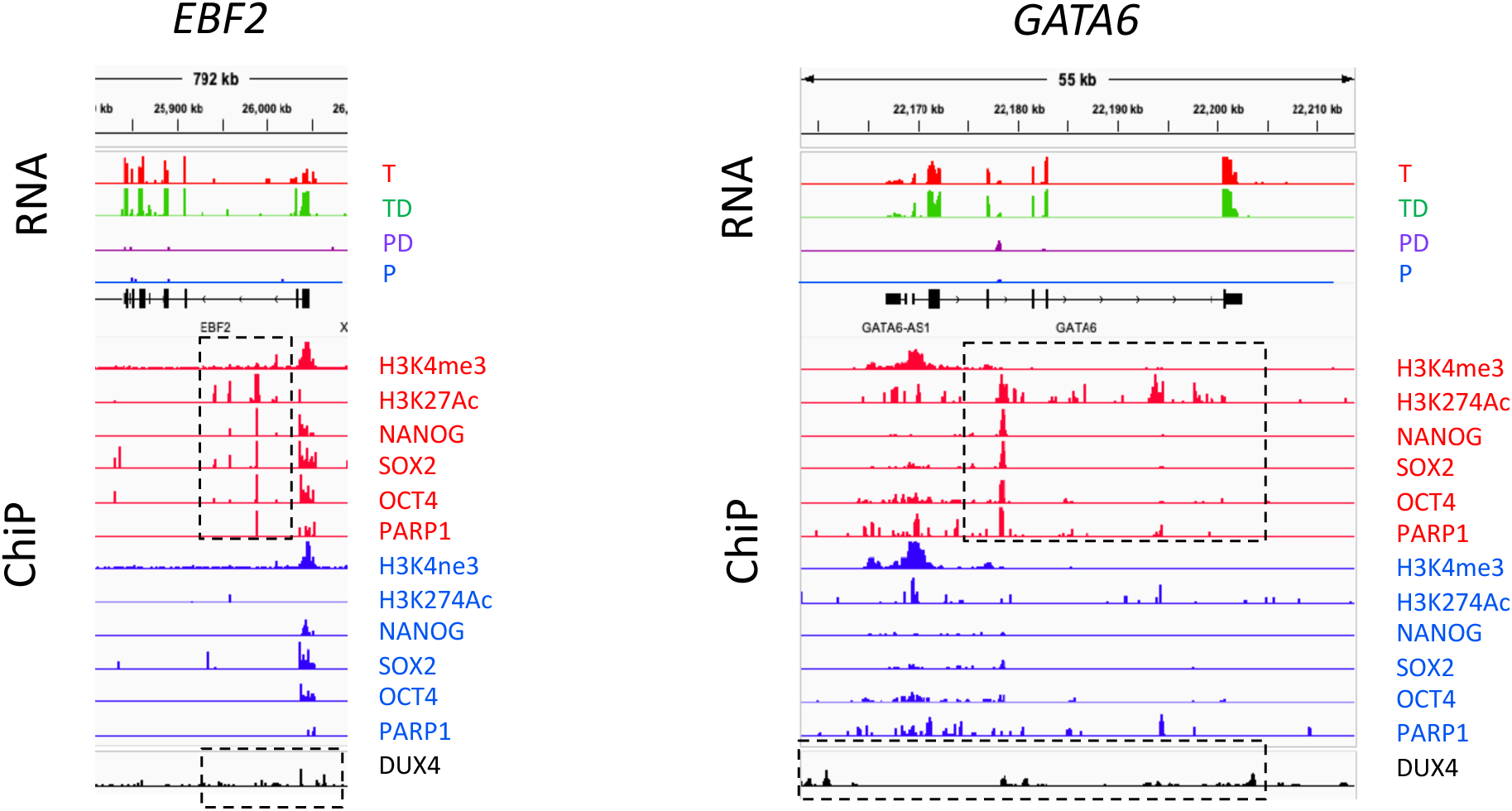
Additional examples of ChIP-Seq read profiles of 8C-specific genes at NSOP regulatory sites. Additional Integrative Genomics Viewer snapshots of RNA-seq and ChIP-seq tracks for 8C-specific gene loci for *EBF2* and *GATA6*, showing RNA expression signal in TIRN (T), TIRN+iDUX4 (TD), primed + iDUX4 (PD) and primed (P) samples and the averaged input-subtracted RPGC-normalized binding of H3K4me3, H3K27ac, NANOG, SOX2, OCT4 and PARP1 in TIRN-SC (red) and primed (blue) samples. Averaged, and genome-aligned DUX4 signal from published DUX4-ChiP data ^8^ is shown (black). Boxed regions highlight novel NSOP-co-binding putative enhancer regions with H3K27Ac and H3K4me3 enrichment.

**Figure S8.**
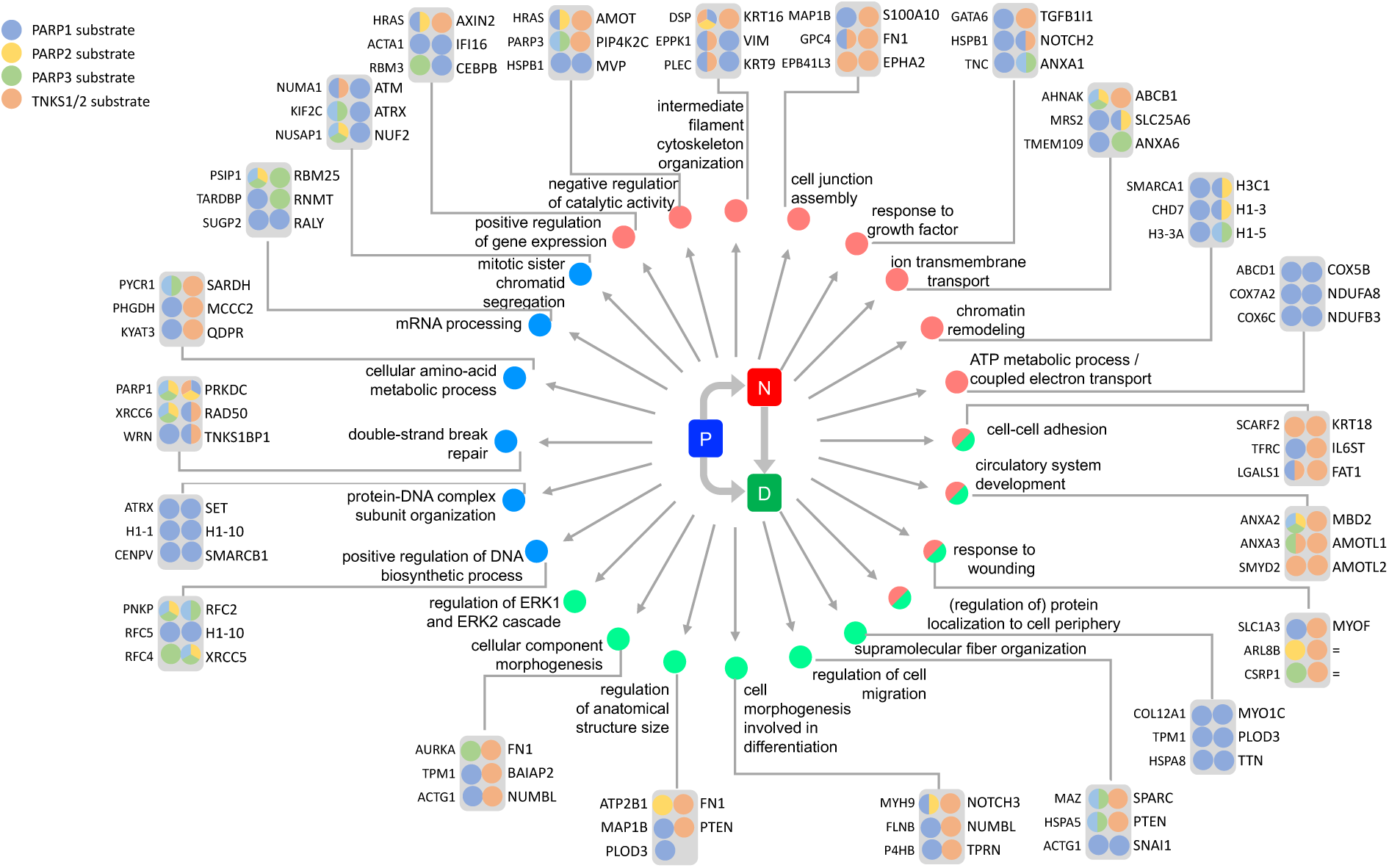
TNKS1/2-PARP1/2 substrate-driven proteomic GSEA pathways in Primed hPSC vs. TIRN and TIRN + iDUX4 stem cells. Summary diagram of GSEA of primed, TIRN, and TIRN+iDUX4 proteome data highlighting known, validated TNKS1/2-PARP1/2 substrates in each category (annotated from Ayyappan, V. *et al* ^5^).

**Figure S9.**
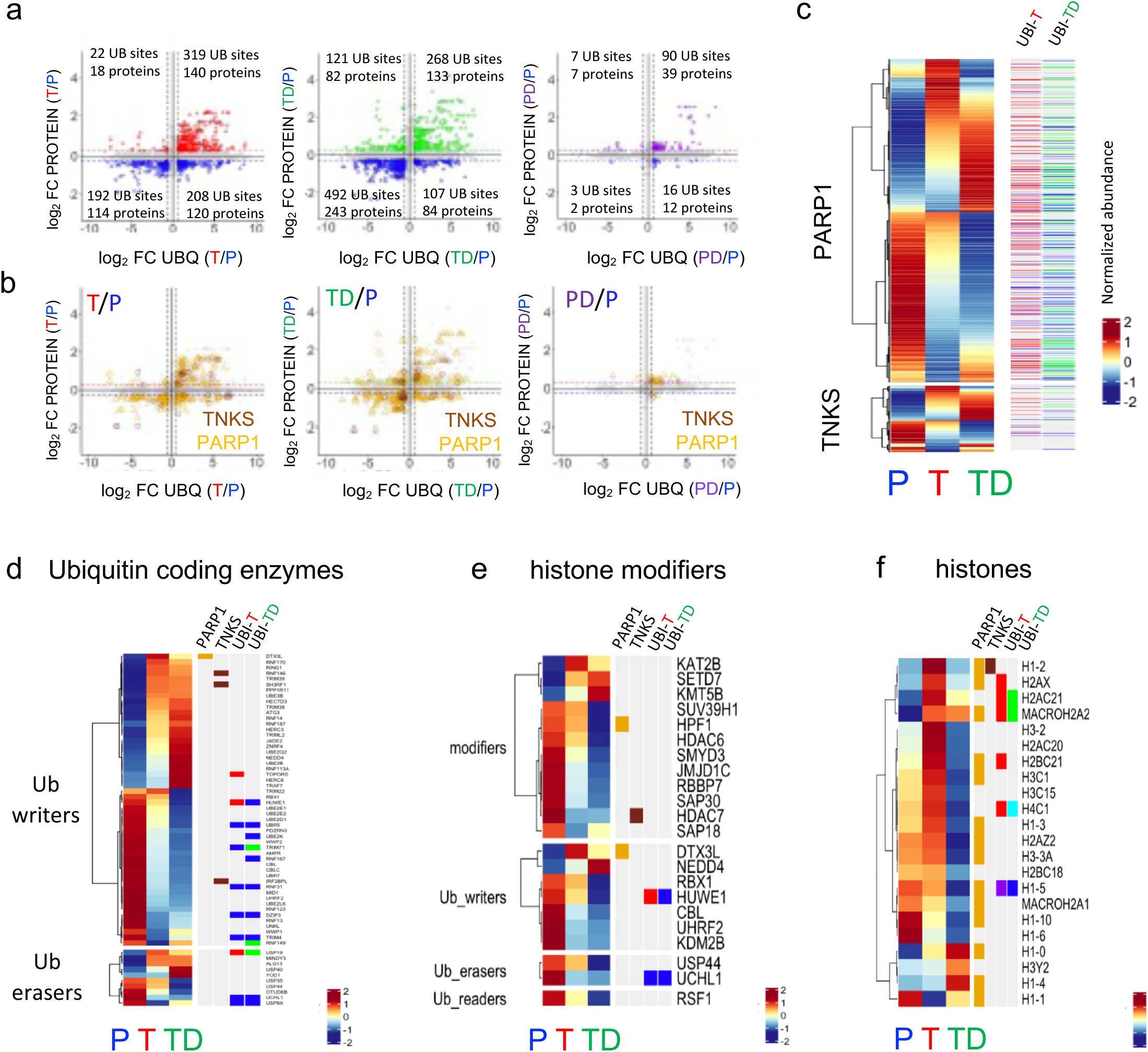
Paired whole proteome and ubiquitinome analyses of Primed, Primed + iDUX4, TIRN, and TIRN + iDUX4 stem cell lines. (**a**) Cross-plots of ubiquitinome sites (x-axis) and proteome (y-axis) relative expression in TIRN (T) vs primed (P), TIRN+iDUX4 (TD) vs primed (P), and primed+iDUX4 (PD) vs primed (P) cells showing the most differentially expressed/ubiquitinated proteins (dashed line boxes). (**b**) Cross plots of ubiquitinome sites (x-axis) and proteome (y-axis) relative expression in TIRN (T) vs primed (P), TIRN+iDUX4 (TD) vs primed (P), and primed+iDUX4 (PD) vs primed (P) cells highlighting the most differentially expressed/ubiquitinated PARP1 (gold triangles) and TNKS1/2 (maroon circles) substrates. (**c**) Heatmaps of PARP1/TNKS substrates in primed (P), TIRN (T) and TIRN+iDUX4 (TD) stem cells with annotations for differentially ubiquitinated proteins (p<0.05, FC>1.5 for TIRN in red and TIRN+iDUX4 in green, and FC<-1.5 for primed in blue. Heatmaps of differentially expressed proteins (FC>1.2, p<0.05) for (**d**) ubiquitin coding enzyme machinery (**e**) histone modifiers, and (**f**) histones in primed (P), TIRN (T) and TIRN+iDUX4 (TD) with annotations for PARP1/TNKS substrates and differential ubiquitination in TIRN (UBI-T, up=red, down=blue) and TIRN+iDUX4 (UBI-TD, up=green, down=blue).

## Notes

### Competing Interest Statement

The authors have declared no competing interest.

