## Supplemental movies for "Proteogenomic Reprogramming to a Functional Human Totipotent Stem Cell State via a PARP-DUX4 Regulatory Axis": MOVIE S3. Ubiquitination Site copy 2.pptx

### Slide 1
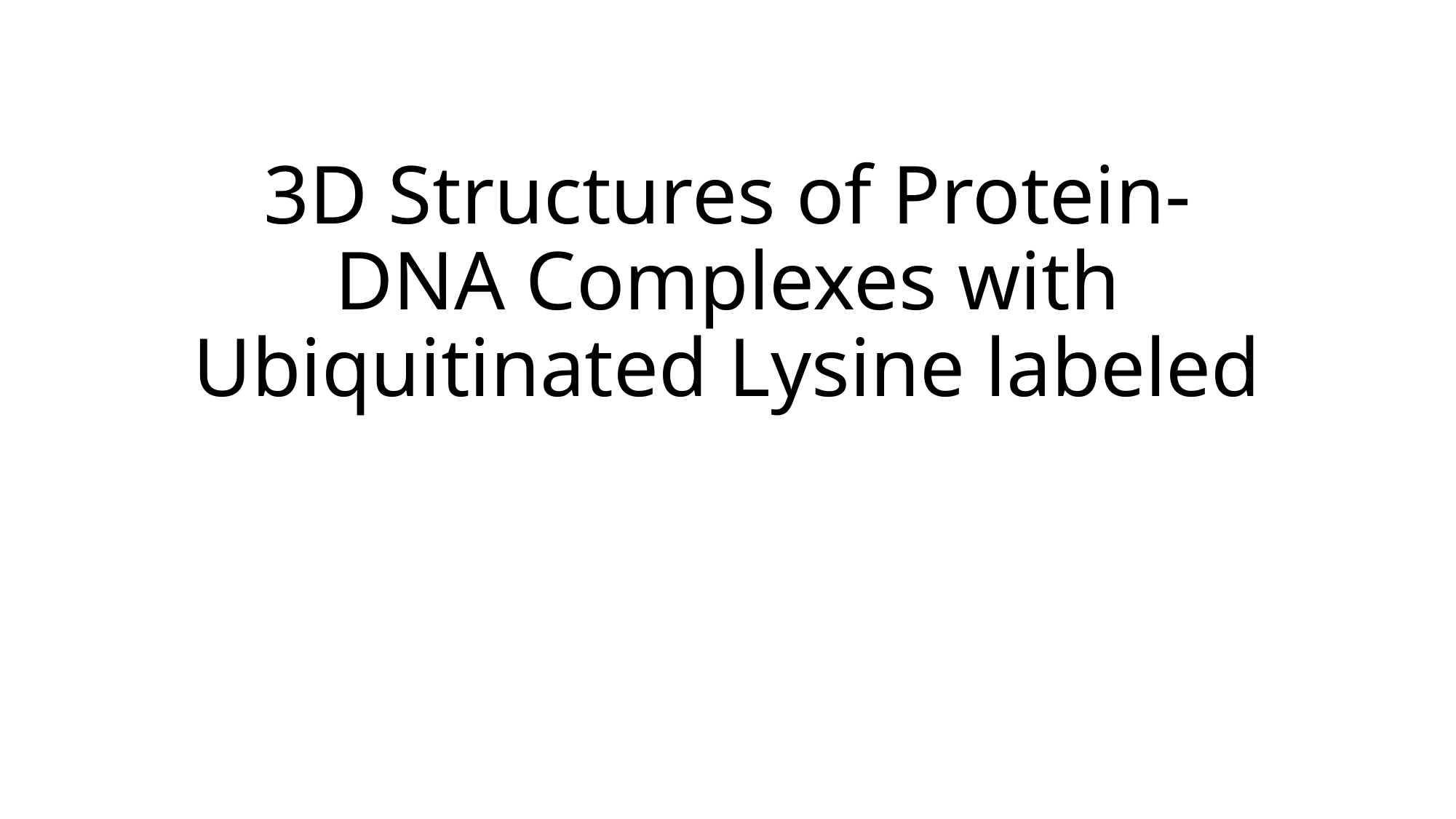

#### 3D Structures of Protein-DNA Complexes with Ubiquitinated Lysine labeled

### Slide 2
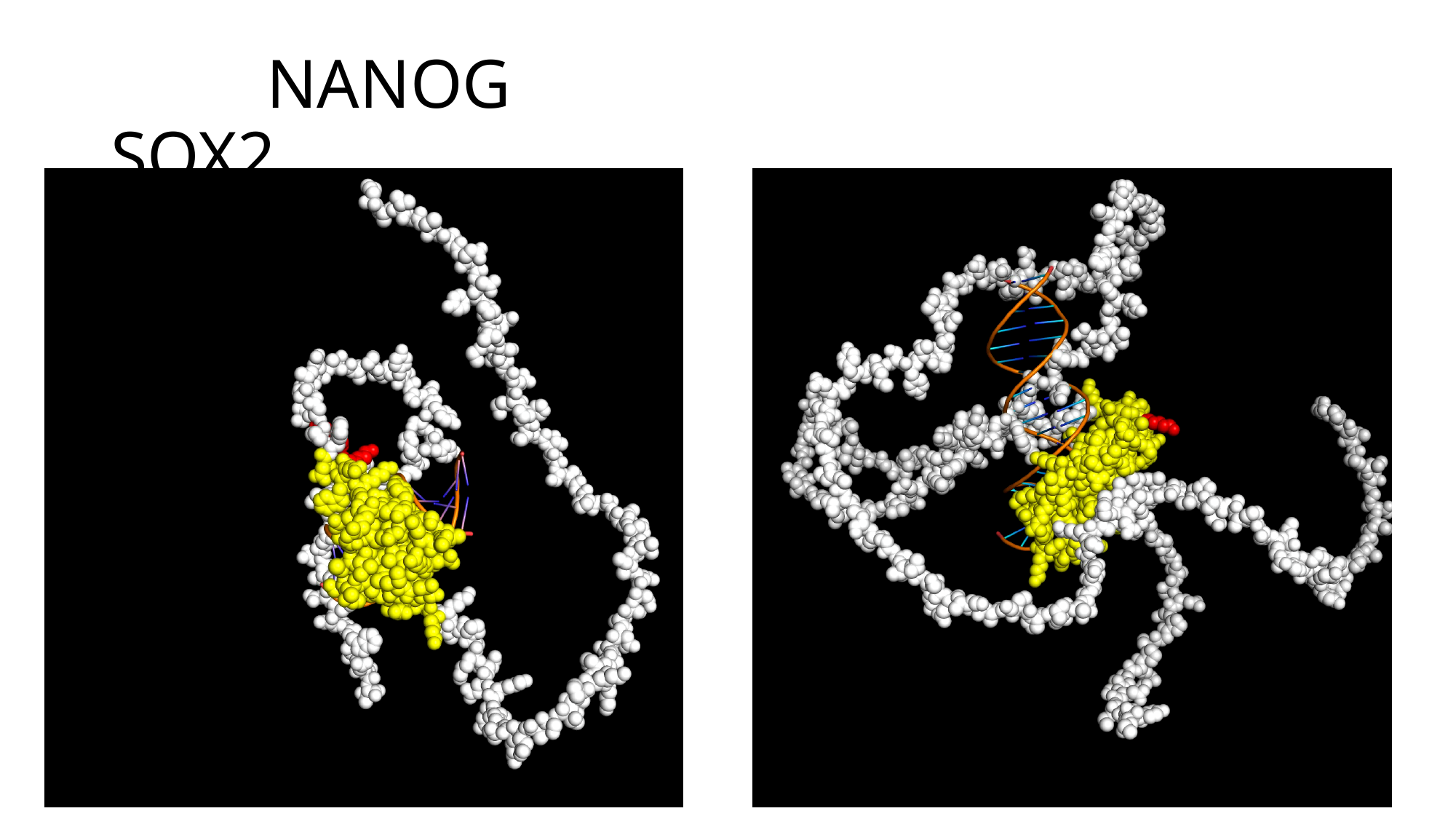

#### NANOG SOX2

### Slide 3
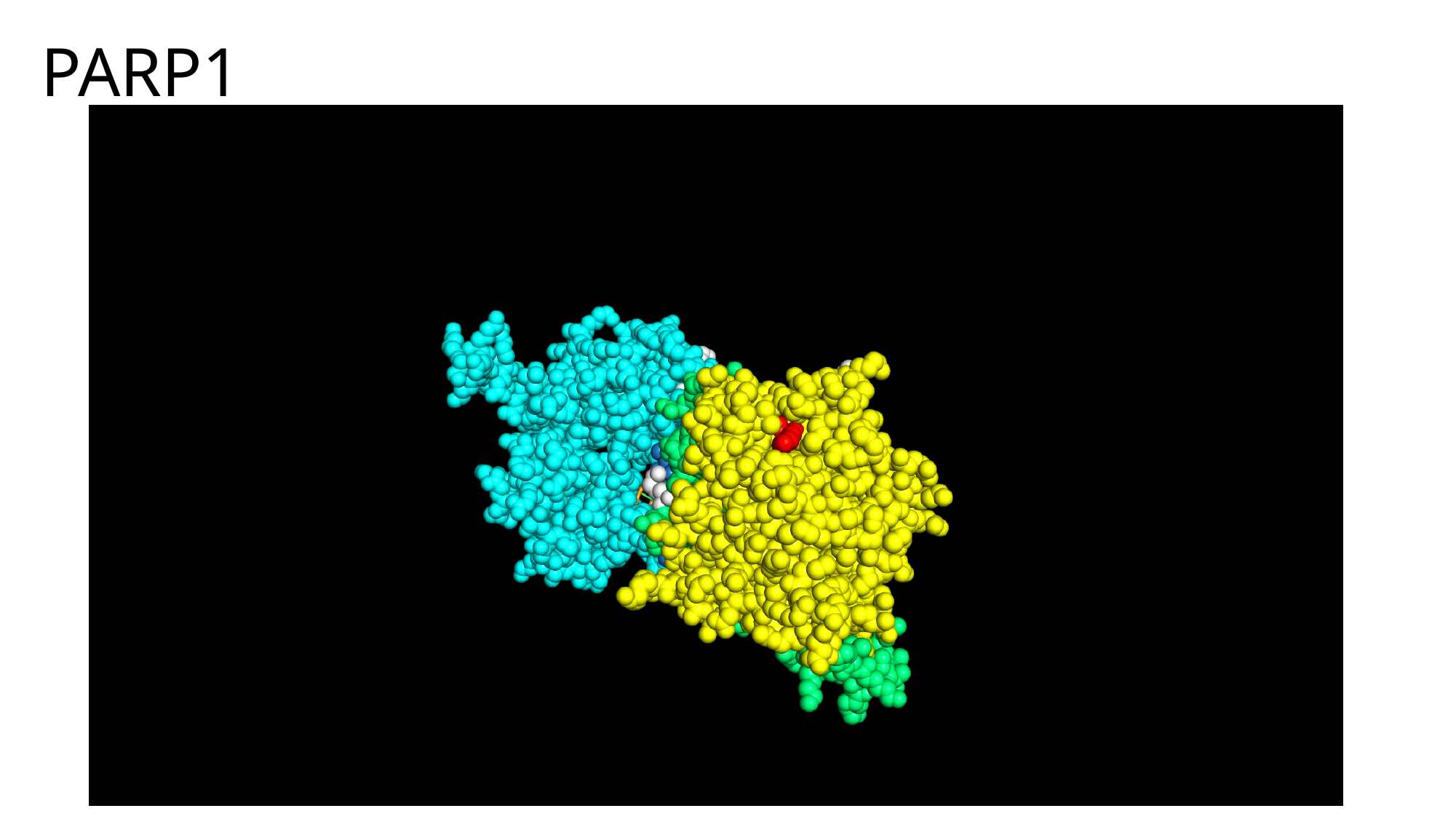

#### PARP1

### Slide 4
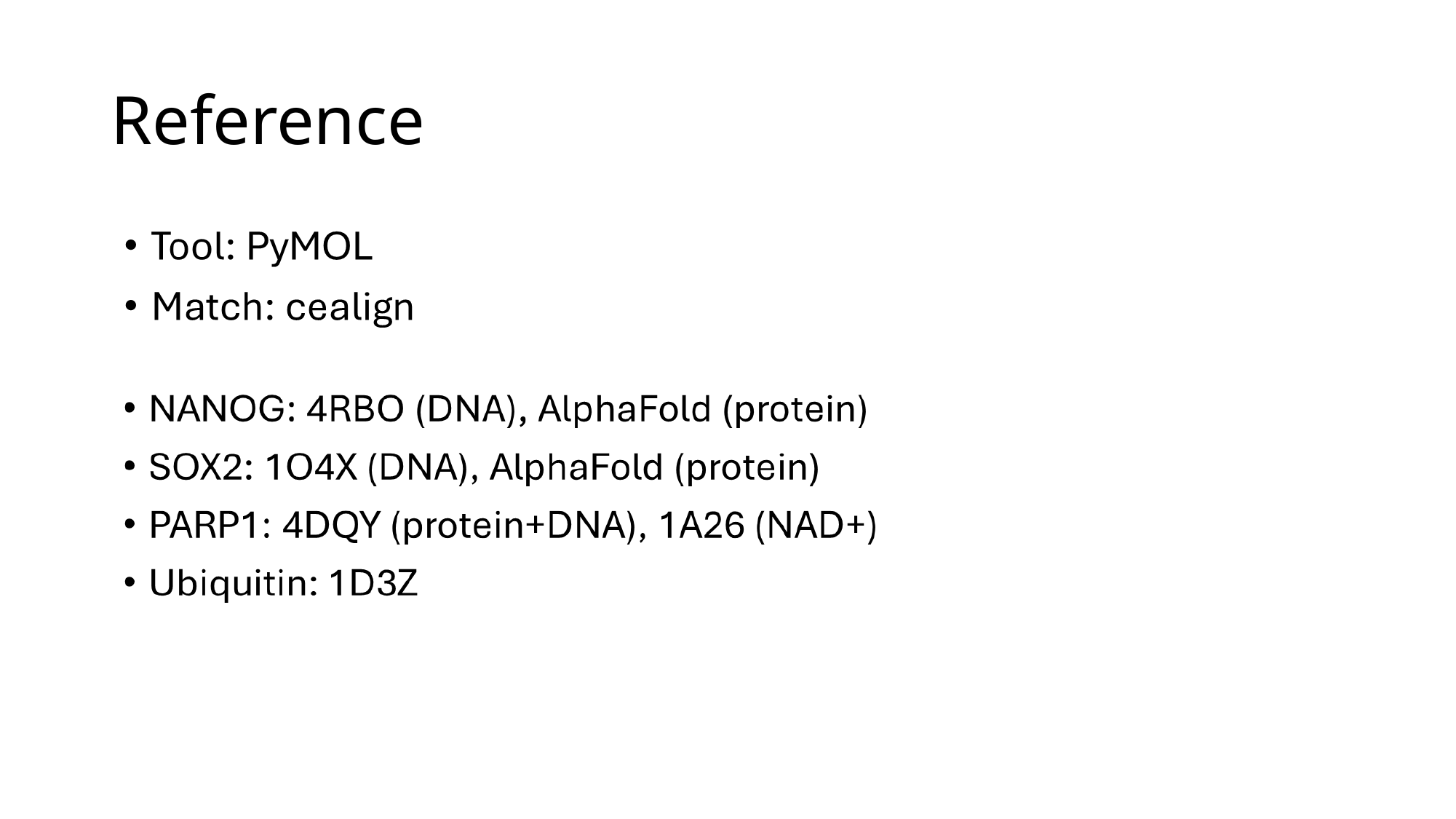

#### Reference
