## Supplemental movies for "Proteogenomic Reprogramming to a Functional Human Totipotent Stem Cell State via a PARP-DUX4 Regulatory Axis": MOVIE-S1-animation embryos1-3 copy.pptx

### Slide 1
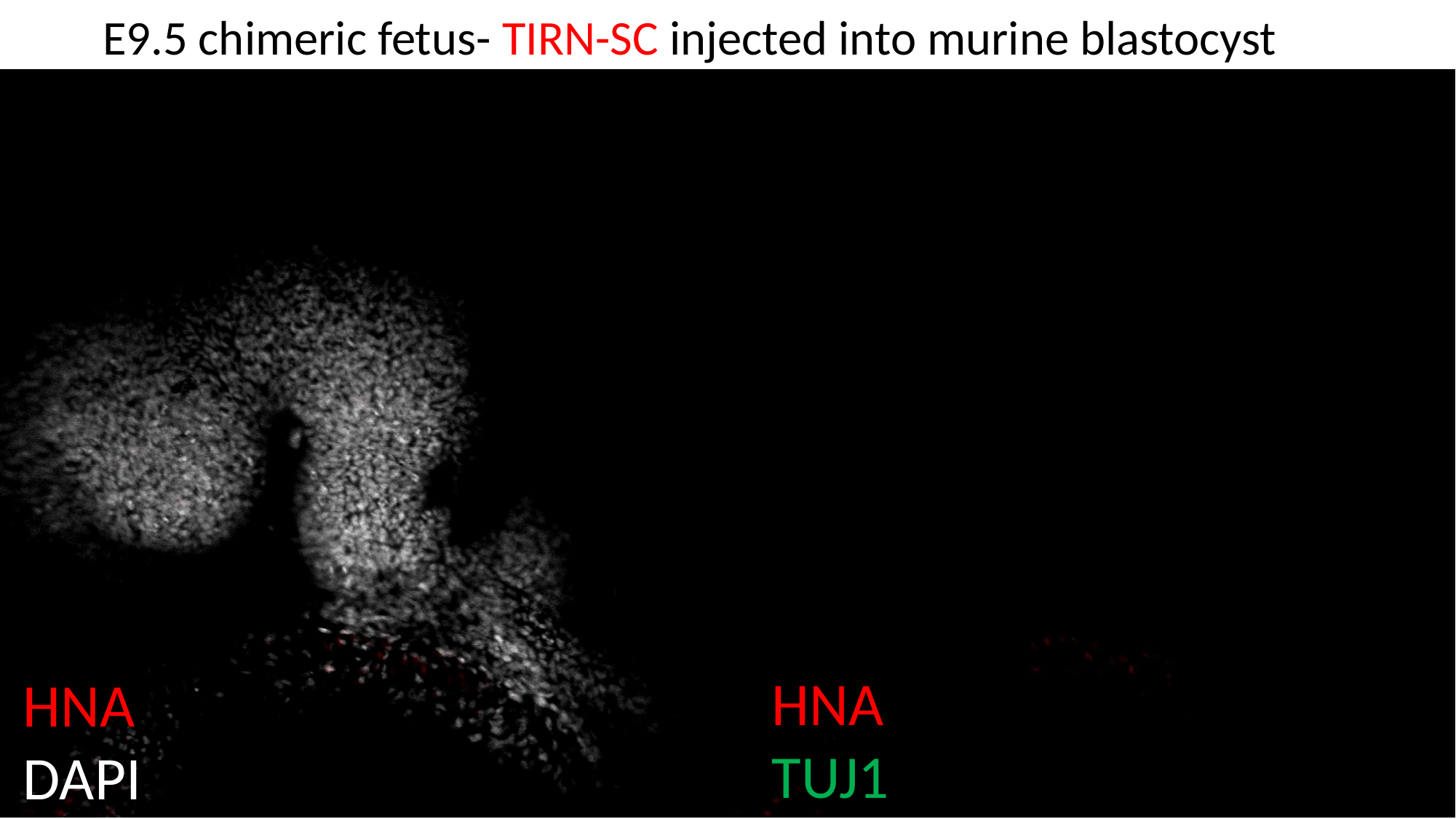

E9.5 chimeric fetus- TIRN-SC injected into murine blastocyst
HNA
TUJ1
HNA
DAPI

### Slide 2
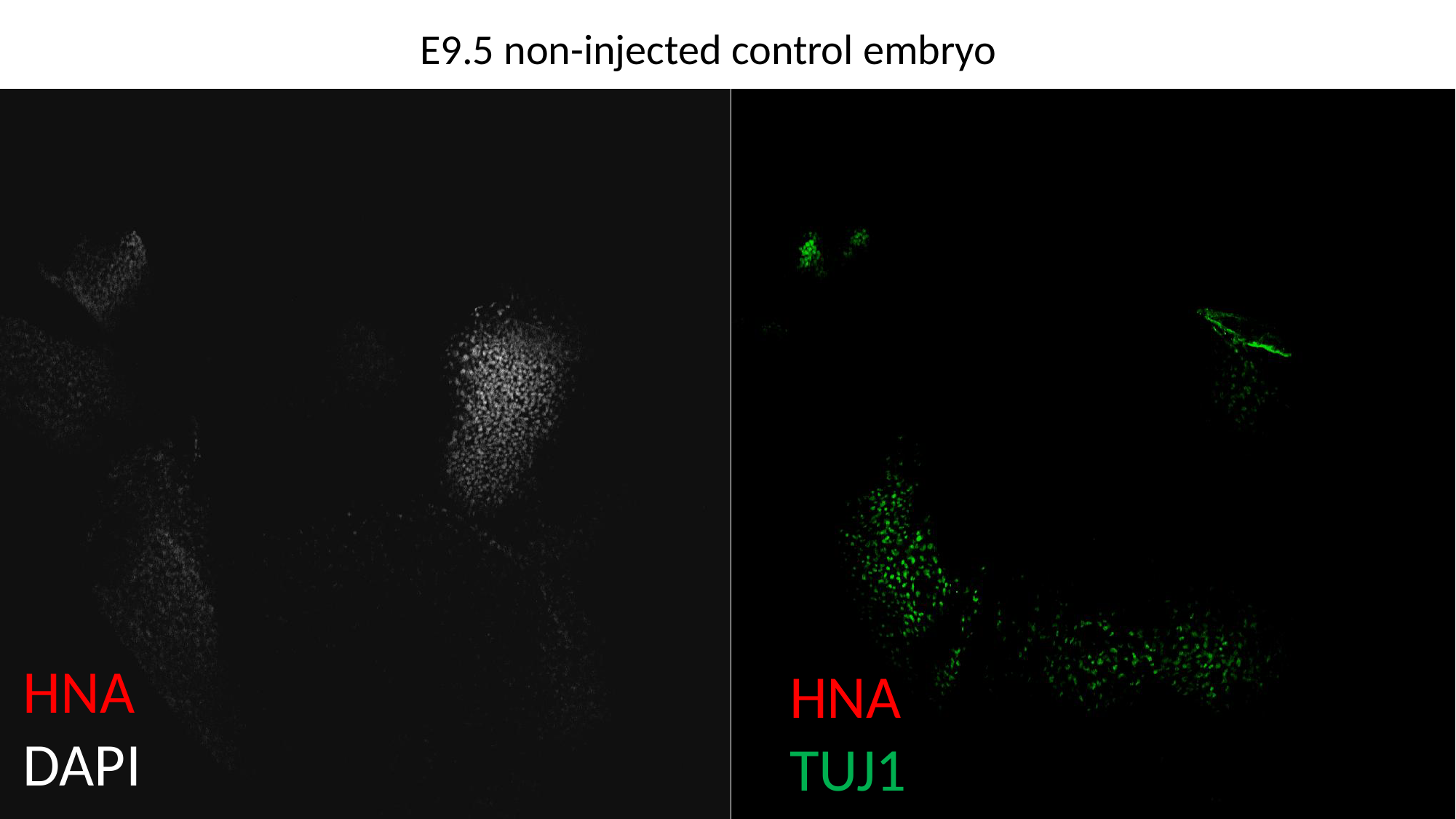

E9.5 non-injected control embryo
HNA
DAPI
HNA
TUJ1
