## Supplemental movies for "Proteogenomic Reprogramming to a Functional Human Totipotent Stem Cell State via a PARP-DUX4 Regulatory Axis": MOVIE-S2-animation embryos2 copy.pptx

### Slide 1
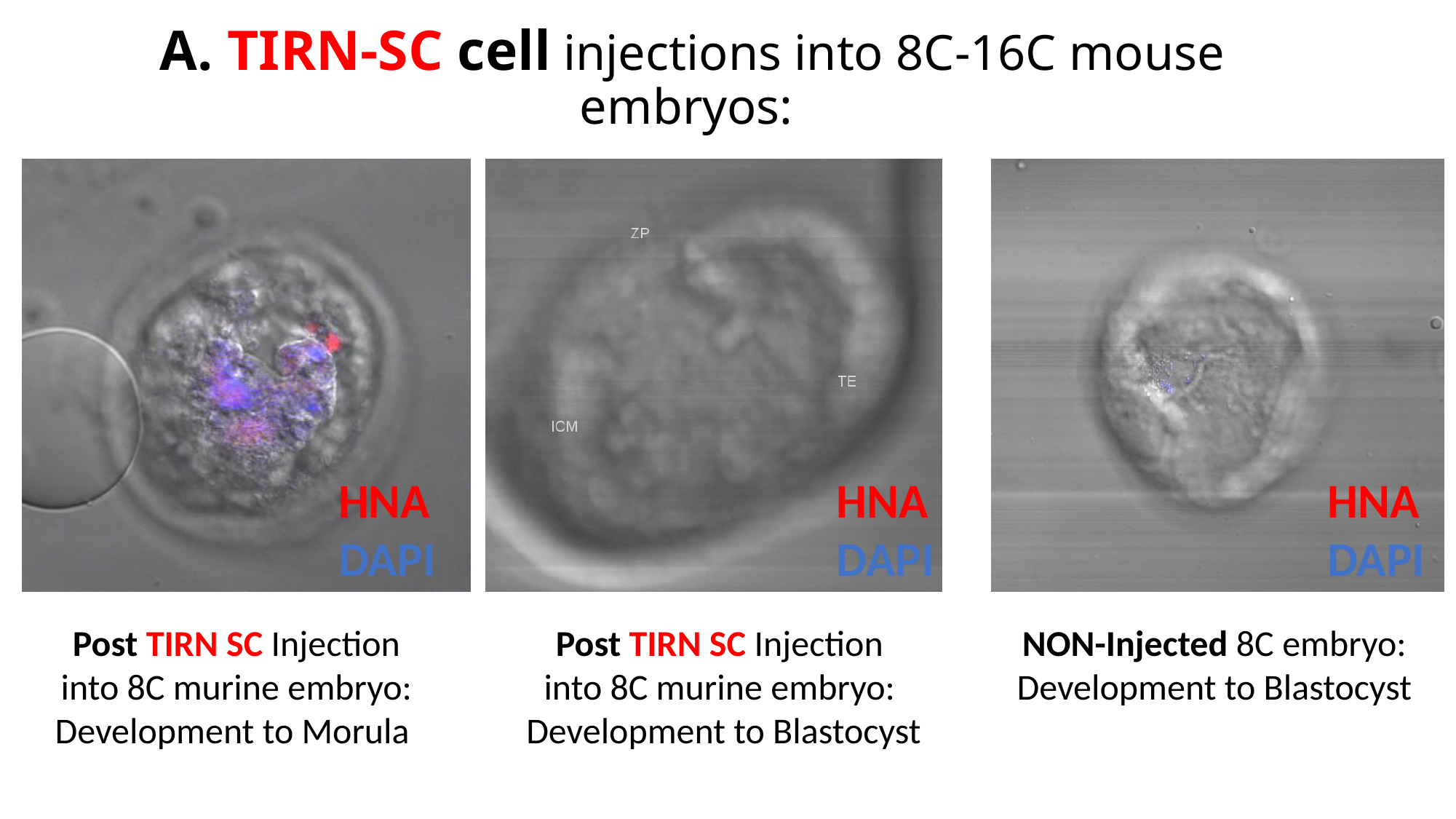

#### A. TIRN-SC cell injections into 8C-16C mouse embryos:
HNA
DAPI
HNA
DAPI
HNA
DAPI
Post TIRN SC Injection into 8C murine embryo: Development to Morula
Post TIRN SC Injection
into 8C murine embryo:
Development to Blastocyst
NON-Injected 8C embryo: Development to Blastocyst

### Slide 2
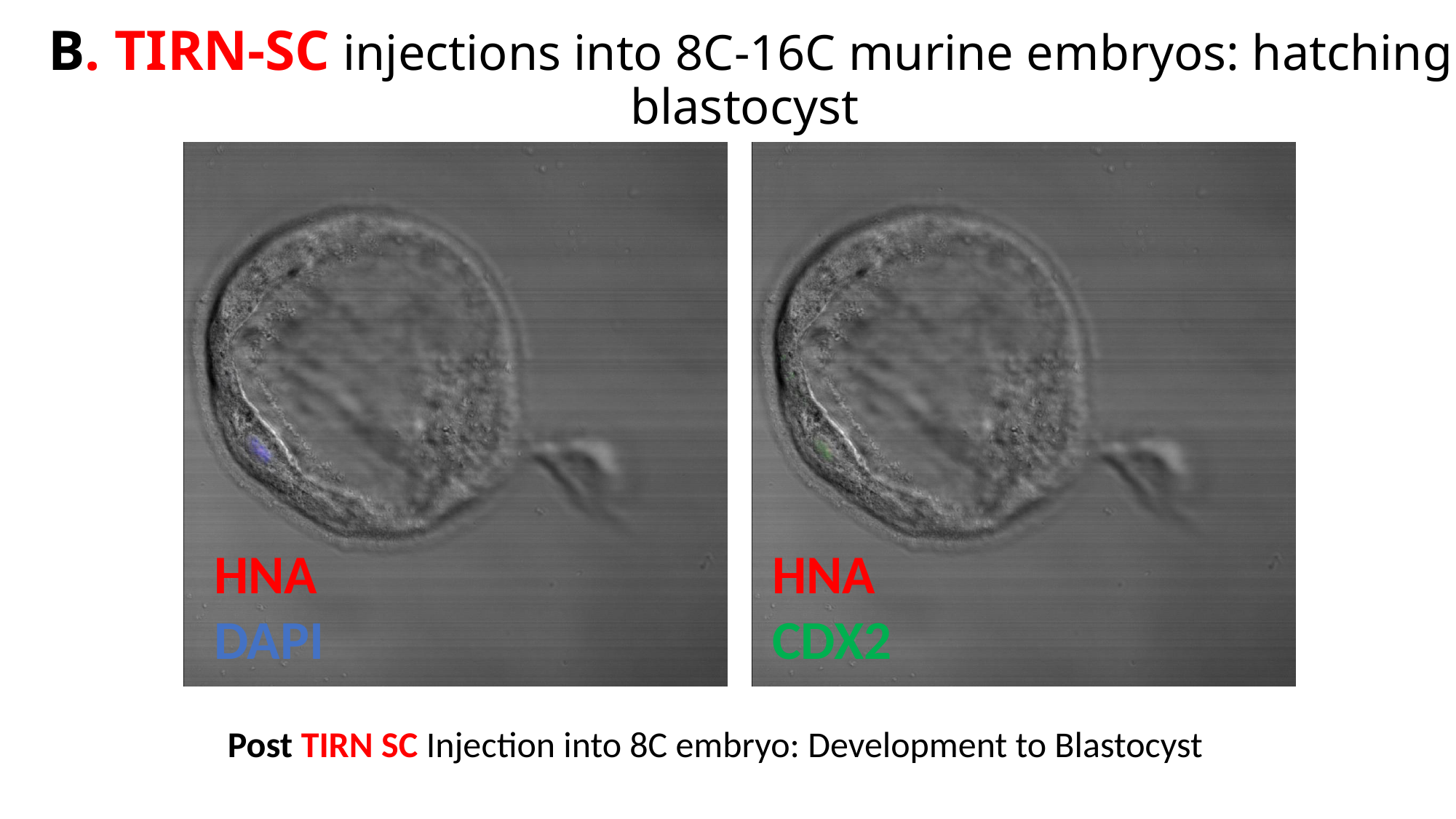

B. TIRN-SC injections into 8C-16C murine embryos: hatching blastocyst
HNA
DAPI
HNA
CDX2
Post TIRN SC Injection into 8C embryo: Development to Blastocyst

### Slide 3
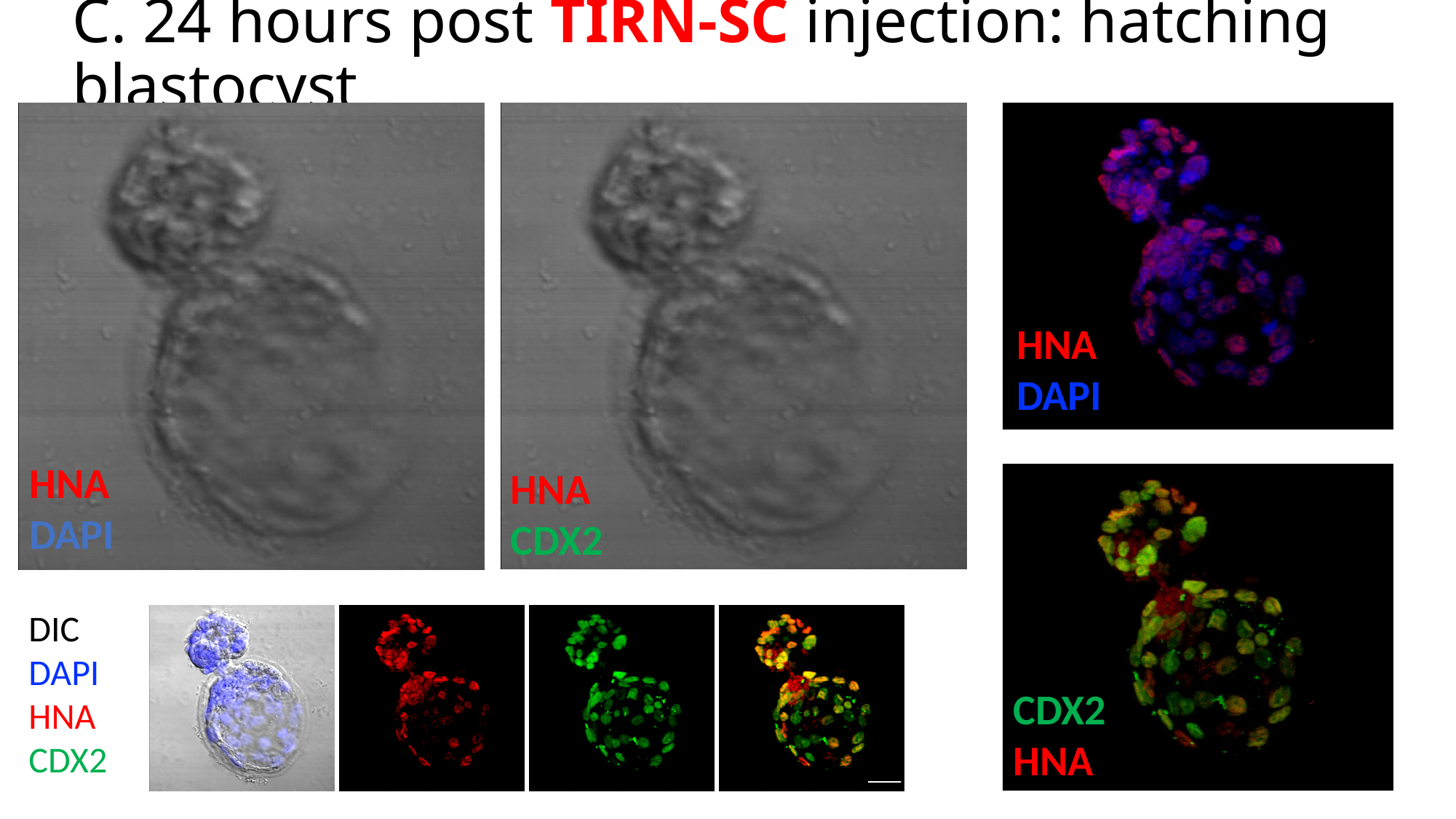

#### C. 24 hours post TIRN-SC injection: hatching blastocyst
HNA
DAPI
HNA
DAPI
HNA
CDX2
DIC
DAPI
HNA
CDX2
Max intensity
Z stack
CDX2
HNA

### Slide 4
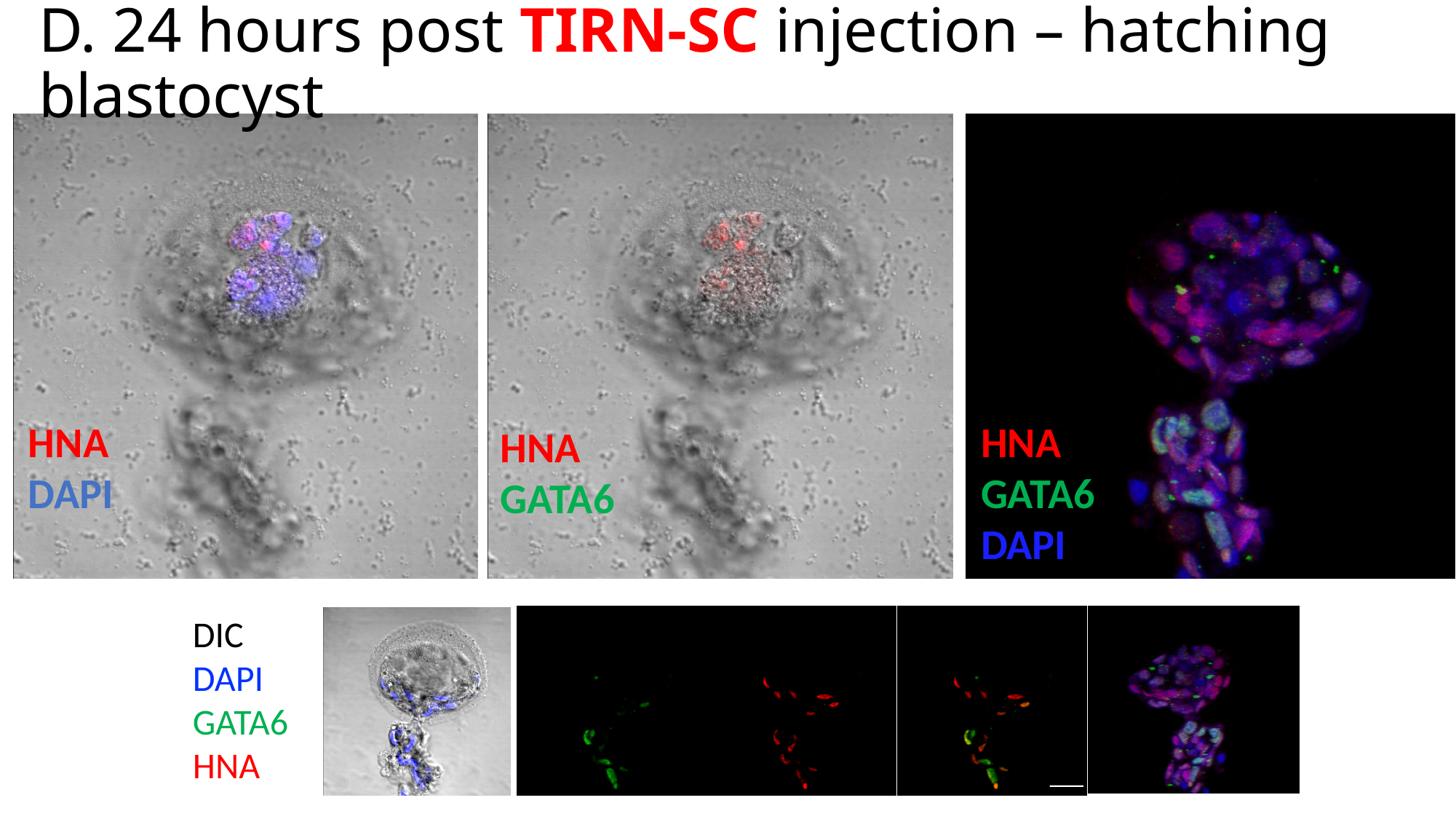

#### D. 24 hours post TIRN-SC injection – hatching blastocyst
HNA
DAPI
HNA
GATA6
DAPI
HNA
GATA6
DIC
DAPI
GATA6
HNA
From 3d
